# Principles of subclonal gene dosage across human cancer

**DOI:** 10.64898/2026.03.15.711906

**Authors:** Sólrún Kolbeinsdóttir, Vasilios Zachariadis, Muyi Yang, Luuk Broeils, Christian Sommerauer, Huaitao Cheng, Xinsong Chen, Yingbo Lin, Matías Marín Falco, Johanna Hynninen, Sampsa Hautaniemi, Yizhe Sun, Olli Lohi, Merja Heinäniemi, Suzanne Egyhazi Brage, Dhifaf Sarhan, Nikolas Herold, Johan Hartman, Hildur Helgadóttir, Felix Haglund de Flon, Anna Vähärautio, Martin Enge

**Affiliations:** Department of Oncology-Pathology, Karolinska Institutet, Solna, Sweden; Paediatric Oncology, Astrid Lindgren Children’s Hospital, Karolinska University Hospital, Stockholm, Sweden; Princess Máxima Center for Pediatric Oncology, Utrecht, Netherlands; Oncode Institute, Utrecht, The Netherlands; Department of Medicine, Karolinska Institutet, Stockholm, Sweden; Research Program in Systems Oncology, University of Helsinki, Helsinki, Finland; Department of Obstetrics and Gynaecology, University of Turku and Turku University Hospital, Turku, Finland; Department of Laboratory Medicine, Karolinska Institutet, Huddinge, Sweden; Center for Cancer Eradication Research, Faculty of Medicine and Health Technology, Tampere University and Tays Cancer Center, Tampere University Hospital, Tampere, Finland; Institute of Biomedicine, School of Medicine, University of Eastern Finland, Kuopio, Finland; Department of Women’s and Children’s Health, Karolinska Institutet, Solna, Sweden; Department of Oncology, Karolinska university Hospital, Solna, Sweden; Foundation for the Finnish Cancer Institute, Helsinki, Finland

## Abstract

Intratumor heterogeneity drives disease progression, but even though subclonal copy number variation (CNV) is a major contributor to this heterogeneity^1^, its impact on cell phenotype is not fully understood. Here, by applying high-quality joint whole genome sequencing and mRNA profiling in single cells (DNTR-seq^2^) to solid tumors and leukemias from 57 patients, we have analyzed the *in vivo* transcriptional effect of subclonal CNVs within a tumor. We found that gene dosage is generally additive in low and moderate copy states, but that cancer-type-specific compensation is common, and core promoter elements are associated with reduced additivity. We find that different classes of subclonal CNV impose varying degrees of transcriptional constraints on a cell, with highly amplified megabase-size regions associated with a strong effect both in *cis* and *trans*, while arm level CNVs generally exert a milder effect. We also describe a previously unappreciated class of tumors with *transient clonality*, where every cell is genetically highly distinct. We find that transient clonality is common in ovarian cancer and soft tissue sarcoma, that it is preceded by a whole genome duplication event, and that gene dosage in these tumors affects transcript abundance to a similar degree as in cancer with stable subclones.

## Main

At its core, cancer is a disease of the genome, where single nucleotide variations (SNVs) and structural aberrations such as copy number variants (CNVs) give rise to altered gene function or altered expression patterns, which manifests as inappropriate growth in a cell-type-specific process that is only partially understood. As highlighted by recent single-cell studies, this is an ongoing process relying on intratumor heterogeneity^3–8^. Activating or deleterious point mutations typically have dramatic and well characterized effects on cell state^9^; for example, activating mutations in the *RAS* proto-oncogenes drive cell division^10^ while deleterious mutations in the *TP53* tumor-suppressor inhibits cell death signaling^11^. Similarly, highly impactful structural variants resulting in gene fusions or promoter hijacking are often well understood. For example, the *ETV6::RUNX1* fusion in B-ALL reverses *RUNX1* transcription factor activity from transcriptional activation into repression^12^. The functional consequences of CNVs, which are ubiquitous in cancers, are on the other hand not well understood. Studies using large-scale bulk data sets have found wide differences in dosage sensitivity between genes, ranging from an approximately additive effect of copy number on transcript abundance to complete dosage compensation^2,13–18^. What determines this difference is not known – while there are well established examples of direct and indirect negative feedback regulatory mechanisms^19,20^, it is unclear how pervasive these are and how they interact with gene dosage.

Previous work in this field is based on bulk data, which comes with methodological challenges: comparisons are done between tumors with distinct genetic histories and backgrounds, and averages made across multiple cell states and subclones. Therefore, recent studies instead attempt to use single-cell methods allowing analysis of subclonal differences within a patient, which should remove problems associated with bulk analysis. Copy number differences have been inferred from transcriptional profile^21–24^, either alone or by integrating the scRNA-seq with a second single-cell whole genome sequencing (scWGS) dataset. However, these approaches come with their own biases, since their power to detect or match a CNV is dependent on the dosage sensitivity and expression levels of the genes it encompasses. In practice, studies have been limited to detecting large-scale copy number changes that appear in relatively many cells^16,25^. Previous true multiomic methods have not offered sufficient throughput, but with miniaturization of plate-based approaches such as DNTR-seq^2^ and well-DRseq^26^ it is now possible to perform large-scale multiomic sequencing of the genome and transcriptome, bypassing the need for integration or inference and allowing direct measurement of how CNVs affect phenotype. Here, we use DNTR-seq to perform such a large-scale study, profiling the joint single-cell genome and transcriptome of 57 patients from six different cancer types. We find that different classes of CNVs have variable effects on transcription, and that while higher gene dosage generally leads to an increase of transcript levels, the magnitude of the increase is affected by core promoter sequence, and tissue specific exceptions are common. We also identify a class of tumors with fully transient clonality, common in soft tissue sarcomas and ovarian cancers, where most cells are genetically unique with large-scale copy alterations distinguishing them from every other cell, indicating an extremely high rate of genetic change. Thus, by directly analyzing subclonal differences *in vivo* in patient samples across solid tumors and leukemias, our study reveals general principles by which structural variants affect transcription.

### Joint single-cell WGS and mRNA-seq across six cancer types

In order to study the effect of genotype on phenotype we collected tissue samples from 57 cancer patients. We selected six cancer types with contrasting profiles of genomic aberrations. Pediatric acute lymphoblastic leukemia (ALL) and acute myeloid leukemia (AML) have relatively non-complex genetics, with a small number of known driver events. Breast cancer (BC) and melanoma (MEL) are characterized by high mutational load and complex rearrangements, while ovarian high-grade serous carcinoma (OC), and sarcoma (SRC) are heterogeneous but often present with a very high level of structural variation **(Fig 1a)**.^27–29^ A single-cell suspension was prepared from each sample, and cells were sorted directly into 384-well plates where DNTR-seq^2^ was performed. In DNTR-seq, nuclear DNA is processed by direct tagmentation to achieve high-quality single-cell whole genome sequencing (scWGS) at an ultra-low depth of coverage, while cytoplasmic mRNA from the same cell is analyzed in parallel using a smart-seq based method^30^. Using a high-quality full-length mRNA-seq protocol enables gene fusion and enhanced SNV detection, in addition to estimating mRNA abundance. The choice of method has several important implications for the robustness of the analysis. First, the two modalities from the same cell are separated as an initial step while the cell nucleus is still intact, ensuring that there is no cross-contamination between them, which is a concern in one-pot methods^31^. Second, sorting cells into individual wells with rigid singlet gating ensures that cell doublets do not contaminate the data, which would otherwise cause serious issues in the analysis. Third, joint analysis of the mRNA-seq data allows us to remove all cells in S-phase. These cells are otherwise a source of artifacts when inferring CNVs from scWGS data since partially replicated genomes will appear as distinct copy states. Both mRNA and WGS modalities were processed using ASCENT^32^ which enables CNV-based clone detection, high resolution segmentation, and copy number calling of ultra-low coverage WGS data as well as standard alignment and transcript counting for the joint mRNA data. Based on the mRNA-seq data, ASCENT also performs fusion gene calling and detection of candidate oncogenic SNVs in genes with sufficient coverage. After strict quality filtering, the dataset consisted of 23 873 single cell transcriptomes, approximately 500 cells per patient. Visualization of the transcriptomic data in a UMAP revealed that non-cancer cells cluster by cell type and independently of patient identity, as expected in the absence of strong batch effects **(Fig 1b)**. Most individual patient samples appear as clearly distinct clusters, but for some cancer types, patients tended to be more similar, in some cases forming clusters made out of several patients. This tendency was particularly prominent for pediatric ALL and AML, and to a lesser extent visible for OC.

**Figure 1.**
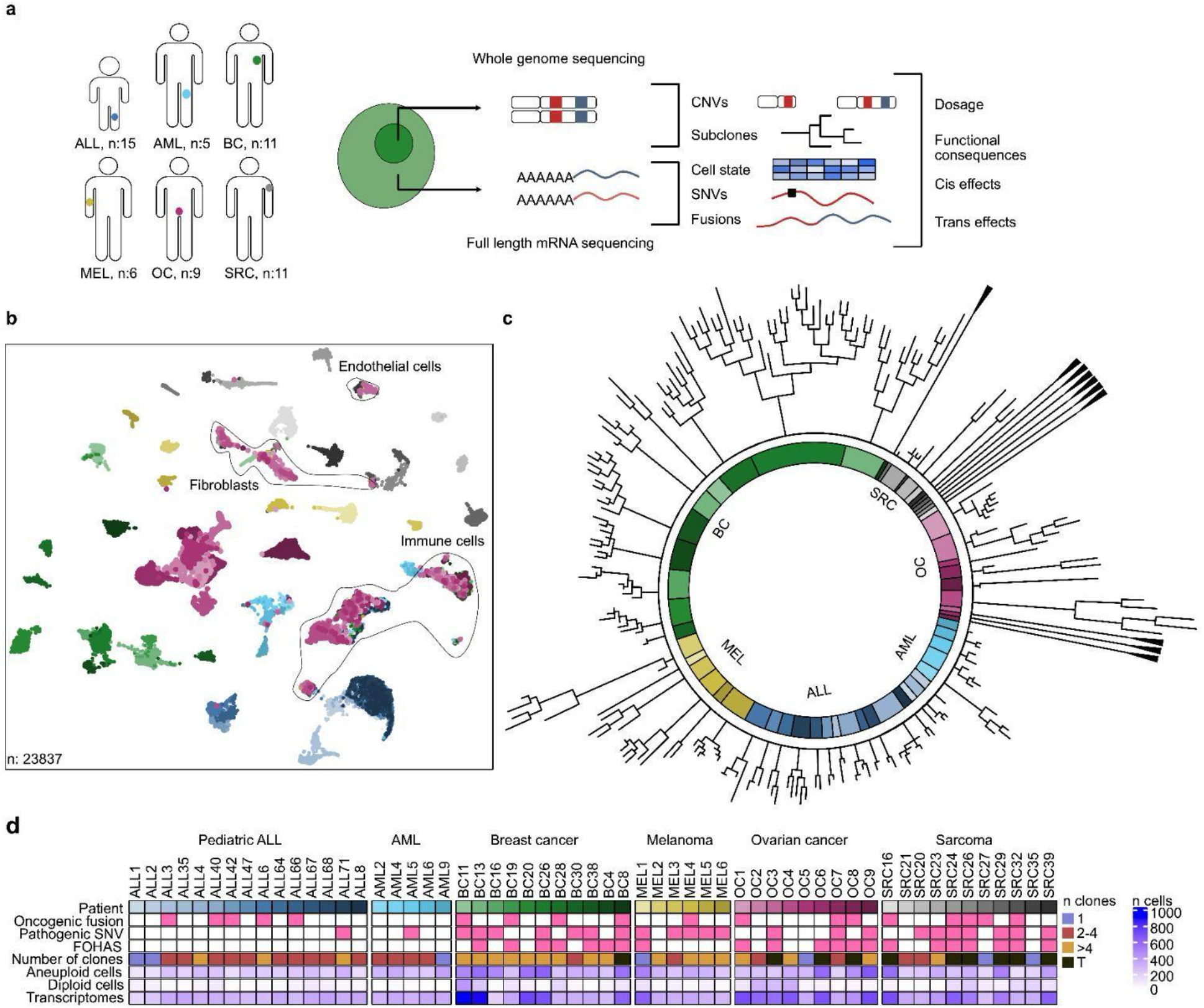
Joint single-cell WGS and mRNA-seq across six cancer types. **a**: Patient samples spanning four solid tumor types and two leukemia types were analyzed, generating joint scWGS and mRNA-seq data from 500 post-QC cells per patient. ALL: Pediatric Acute Lymphoblastic Leukemia, AML: Acute Myeloid Leukemia, BC: Breast Cancer, MEL: Melanoma, OC: Ovarian Cancer, SRC: Sarcoma. **b**: 2D UMAP projection of single-cell transcriptomes, colors indicate cancer type (base color) and patient (shading). Colors correspond to the key in Fig 1d. **c**: Phylogenetic tree based on allele-specific copy numbers of all single-cell genomes sequenced. Trees are rooted in the reference diploid genome. Triangles indicate samples where no clonal structure was found. Branch lengths correspond to genetic distance (minimal number of events). Sample colors correspond to key in Fig 1d. **d**: Dataset overview, indicating which patients have a fusion involving an oncogene, a pathogenic SNV in a COSMIC gene, focal highly amplified segments (FOHAS), how many clones they are composed of (T for transient) and number of cells passing DNA (split into aneuploid and diploid) and RNA QC.

Cancer types exhibited differences in CNV-based subclonal complexity in line with previous literature, with phylogenetic analysis indicating that leukemias harbor few events whereas BC and OC were defined by multiple events and high degree of subclonality **(Fig 1c)**. BC had the highest number of subclones, and was the only cancer type where every sample was subclonal, whereas AML at the other side of the spectrum, had no samples with more than four subclones (**Fig 1c** and **Suppl. Fig 1a**). BC, SRC and OC also included a subset of patients with very high intra-patient genetic heterogeneity, but with no detectable clonal structure. These cases did not exhibit stable clonality where the same CNV profile is detected in several cells and where genetic alterations of subclones build upon each other in a hierarchical fashion, but their CNV profiles were instead consistent with a *transient* clonality where the cells gain and lose large chromosomal regions at a high rate (annotated by a triangle shape in **Fig 1c**). Further, by analyzing the full-length mRNA-seq data, we found that samples from 18 of the patients harbored fusions involving an oncogene, and we detected pathogenic SNVs in oncogenes and tumor suppressor genes in 24 patients. Focal highly amplified segments (FOHAS) were found in the more complex epithelial tumors and all of the cancers with transient clonality (**Fig 1d** and **Table 1**).

We analyzed each individual patient to reveal a wide range of CNVs (**Extended Data Figure**). Two representative examples, MEL5 and BC13, are shown in Figure 2. MEL5 is a metastatic melanoma with average CNV complexity (**Fig 2a**), where we detected five subclones that all had high average ploidy, suggesting an early whole genome duplication (WGD) event with subsequent chromosome losses. BC13 is a HER2 negative, estrogen receptor (ER) and progesterone receptor (PR) positive BC with a complex structure of 13 stable subclones, and separated into two main clades by phylogenetic analysis **(Fig 2b**). Transcriptomic profiling of the cells of each patient revealed non-cancer cell clusters as well as several clusters of aneuploid cells (**Fig 2c, 2d**).

**Figure 2.**
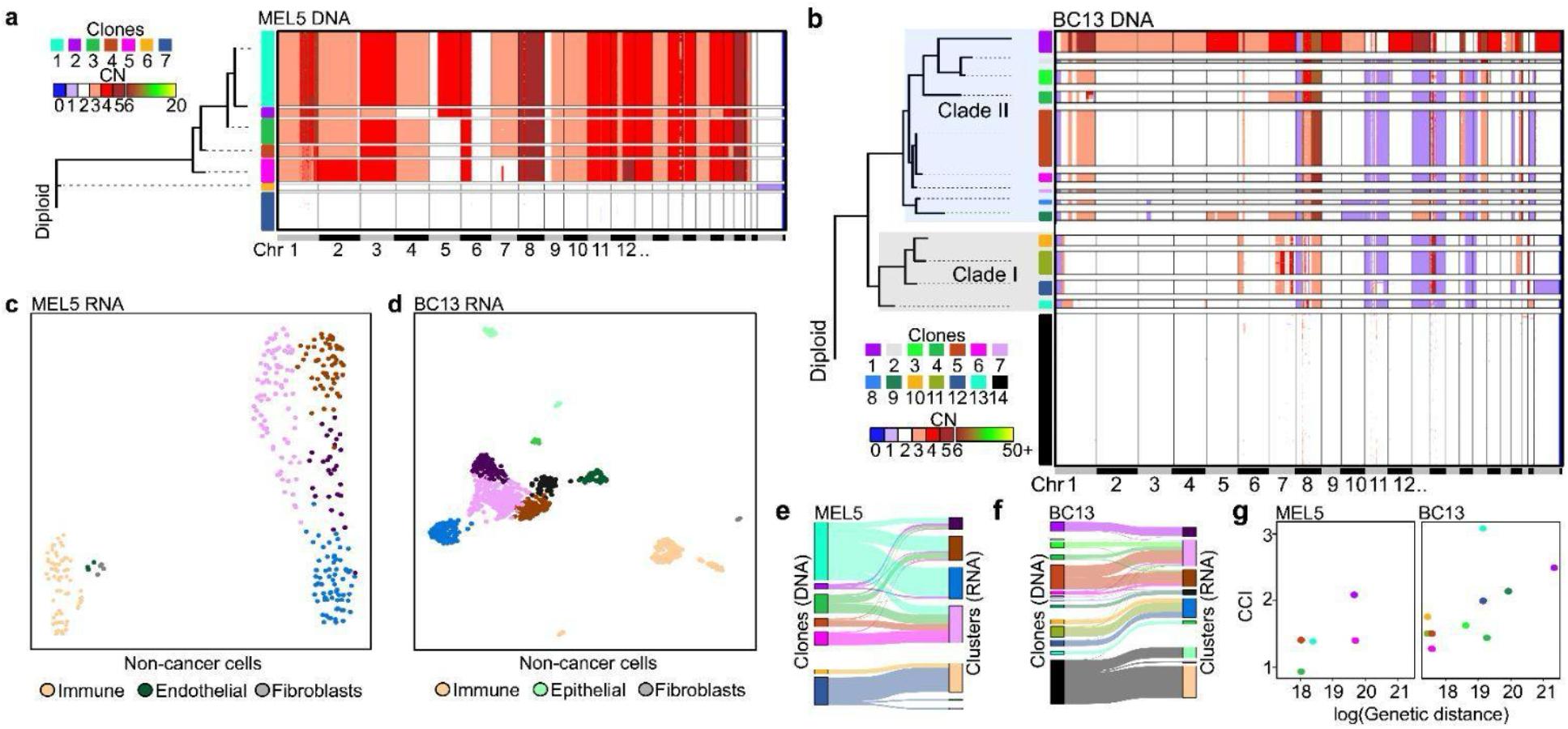
Transcriptional constraints imposed by CNVs can be highly variable. **a**: Single-cell CNV profiles from a melanoma patient (MEL5, n=262). Each cell is a row in the heatmap, horizontal scale shows genomic coordinates. Copy numbers (CN) are indicated by color. Cells are sorted into clones (vertical block colors). Phylogenetic tree of subclones (left panel) was calculated based on maximum parsimony of allele-specific CN. **b**: Single-cell CNV profiles from a breast cancer patient (BC13, n=545), as in a **c**: UMAP of single-cell transcriptomes from MEL5 (n=302). Transcriptome-based clusters are shown as colors. Non-cancer cell identities were taken from clustering all non-cancer cells. **d**: Same as c for BC13 (n=891). Dark-green cluster consists of cycling cells. **e**: Alluvial plot of genetic identity (clones, colors as in a), and their correspondence to transcriptomic identity (clusters, colors as in c) for MEL5. Distances between clones are scaled to phylogenetic distances, RNA cluster distance is equal except between the tumor and diploid cells. Non-cycling cells with full information (where WGS and mRNA passed QC) are shown. **f**: Same as e, for BC13. **g**: Clonal constraint index (CCI) of all clones (colors as in a and b) where minimum 7 cells had full information, plotted against the genetic distance of each clone to its genetically nearest neighbor.

One straightforward way to investigate genetically imposed constraints on transcription is to analyze to which extent unsupervised clustering of the transcriptional data recapitulates the clonal structure. For MEL5, transcriptomic-based clusters only partially mirrored the clonal structure, with most clones fairly evenly distributed between clusters but clone 4 and 5 clustering to a specific phenotype (**Fig 2e**). These two clones had in common a relative loss of chromosome (chr) arm 1q, but they also had large-scale differences between them, including a sub-chromosome arm amplification on chr 7, and whole chromosome gain of chr 2. Interestingly, a large number of additional non-cancer cells which had lost a single X chromosome were also identified in MEL5, with the mRNA revealing that they were immune cells (**Fig 2a, 2c, 2e**). The genetically complex breast cancer case BC13 had stronger correlation between clonal structure and transcriptional clusters, and its two main genomic clades were completely nonoverlapping in transcriptomic space (**Fig 2b, 2d, 2f**). Additionally, two subclones had exceptionally distinct transcriptional phenotypes - a clone from clade II which had undergone WGD giving it a very different copy number profile to the other subclones, and the ancestral clone of clade I, with relatively modest differences to other subclones of the clade. In contrast, the three remaining subclones in clade I were transcriptionally very similar despite genetic differences encompassing either whole chromosome loss of chr X and partial loss of chr 11 and 19, or an additional amplification on chr 7.

In order to compare the degree of genetic constraints between clones in an unbiased fashion across different levels of genetic and transcriptional complexity, we developed a more systematic approach. We calculated a clonal constraint index (CCI) for each clone, an estimator of the likelihood that transcriptionally similar cells belong to the same clone. CCI was calculated as a confusion matrix based on the probability that nearest-neighbors in transcriptional space of subsampled data originated from the same clone, compared to expected (see *Methods* for details). CCI scores are high when clonal traits constrain transcriptional phenotype strongly compared to the influence of random noise and non-clonal cell states, and approximately zero when the transcriptional effect of subclonality is weak. Importantly, a CCI score of zero does not exclude the possibility that there are transcriptional effects, but it suggests that the effect is relatively insignificant compared to other factors. In the case of MEL5 and BC13, every clone had a CCI score higher than zero, indicating that all of the clones exerted detectable constraint on the phenotype in these patients. Overall, the amount of genetic material with a unique copy state in a clone correlated with its degree of transcriptional constraint in both MEL5 and BC13, with larger genetic distance to the nearest-neighbor clone being associated with higher CCI (**Fig 2g**). However, we also commonly observed exceptions, where large-scale genetic events had little impact on transcriptional profile. Our data highlights that even in these highly controlled comparisons – within patient samples and therefore comparing subclones in the same tissue context and genetic background – the transcriptional constraints imposed by large-scale genetic alterations can be highly variable.

### Genotype-dependent transcriptional constraints vary by CNV class and are driven by both cis and trans effects

To address the fundamental question of how strongly genetically distinct subclones determine transcriptional phenotype, we next analyzed the transcriptional impact of distinct subclones across all data. In our data set, genetic changes shared within a patient recapitulate the full history of past tumor evolution, whereas subclones capture the genetic heterogeneity at the sampled timepoint, a snapshot of ongoing tumor evolution. As expected, we found that subclonal differences explained a minor but highly significant part of the transcriptional profile of cells (**Suppl. Fig 2a**). If *in vivo* subclonal genetics represent current genetic heterogeneity, their jointly measured transcriptional profiles could be used to quantify the functional consequences of this heterogeneity. A subclonal CNV profile that does not alter the transcriptional state in a meaningful way is unlikely to contain a driver event. In contrast, a highly distinct transcriptional profile in a subclone implies underlying genetic changes that impose strong phenotypic constraints and likely functional consequences. To quantify the constraints that a specific genotype places on its transcriptional phenotype, we calculated the CCI for each subclone across the full dataset (**Fig 3a**). We found that in some cases, such as ALL68, ALL66, ALL42, AML2 and BC38, subclonal traits imposed no measurable constraints on transcriptional phenotype. In others, such as BC28, BC13 and MEL5, all subclones constrained the phenotype. Lastly, we found cases with highly differential constraint levels, such as BC29 and MEL2, where certain subclones showed strong transcriptional constraints while others did not. MEL2 is an interesting example, since it harbored four subclones with weak transcriptional constraints and one outlier subclone with exceptionally high CCI (**Fig 3b**). The strongly constrained clone carried a highly amplified region of chromosome 3, additional breakpoints on chr 2 as well as several other CNVs across the genome separating it from the other four clones. Differential gene expression analysis between the expression profiles of MEL2 subclones revealed several affected genes directly overlapping the copy-altered regions (effects in *cis*), for example *PCDHB5* which was lowly expressed in subclones that carried a heterozygous deletion. We also observed altered gene expression of genes mapping to non-affected genomic regions (effect in *trans*), such as *ECM1* which had strong clone-specific expression despite no evidence of a CNV. Among genes that were not significantly differentially expressed was *MITF*, which was highly expressed in all clones, with no discernable increase in the clone where it was amplified (**Fig 3c**). *MITF* is a “master regulator” oncogene in melanoma, and it seems likely that its expression is stabilized by feedback regulation in MEL2. These results suggest a complex relationship between CNVs and expression where cis, trans and gene regulatory feedback interact to influence the transcriptional profile.

**Figure 3.**
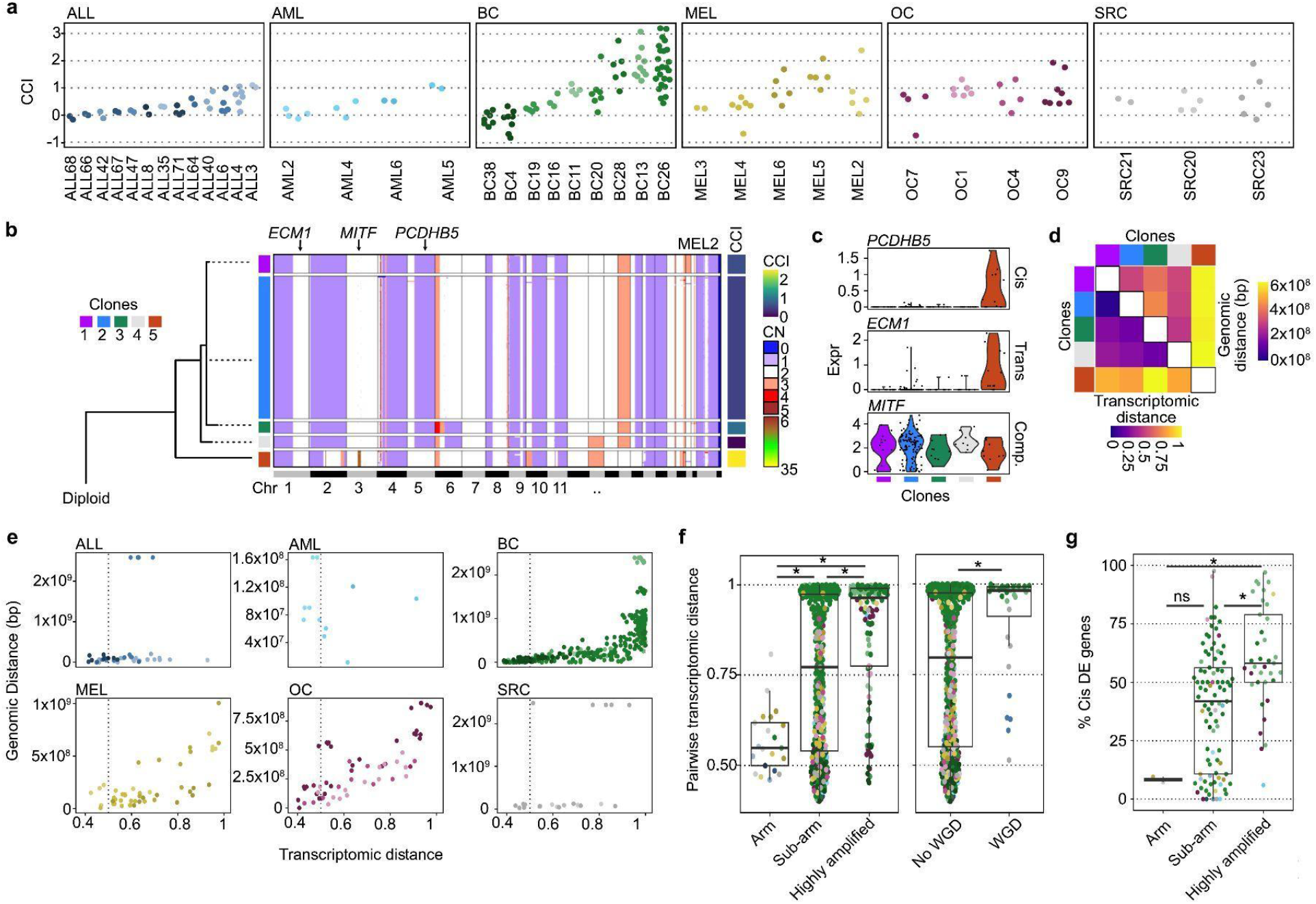
Differential genetic constraints of cell states in stable subclones. **a**: CCI scores across all patients, of every subclone which had at least 7 cells which passed mRNA QC in all patients that had at least 2 subclones. **b**: Single-cell CNV profiles from a melanoma patient (MEL2, n=174), only cells from clones with CCI scores are shown. Heatmap shows each single cell as a row and copy numbers are indicated by color. Cells are sorted into clones (vertical block colors). Phylogenetic tree (left panel) was calculated based on maximum parsimony of allele-specific CN. Color bar at the right side of the heatmap indicates CCI score per clone. **c**: Expression levels of three genes grouped by subclone in MEL2, labeled based on if they represent a *cis* or *trans* effect, or a gene subject to dosage compensation in clone 5. **d**: Pairwise distances in transcriptomic space between the subclones of MEL2 in lower triangle and pairwise distances in genomic space in the upper triangle. **e**: Pairwise comparison of genomic distance against transcriptomic distance between all clones, across all patients (color, as indicated in Fig 1d). Vertical dotted line indicates 0.5. **f**: Pairwise transcriptomic distance between clones separated by different chromosomal aberrations. Left panel: CNV class. Arm=pairs separated by only intact arm-level copy changes, sub-arm=pairs separated by segments with breakpoints within chromosome arm, but with three or less copy difference, highly amplified=pairs with segments differing by more than three copies. Right panel: average absolute ploidy difference. No WGD=average difference<0.9, WGD=average difference>=0.9. Clone pairs are colored by tumor type. Star indicates p<0.05 (Wilcoxon rank sum test). **g**: Percentage of differentially expressed (DE) genes between pairs of clones where the genes are located in regions with differing copy number (transcriptional effect in *cis*). Clone pairs were separated by CNV class as in g. Pairs with at least 10 DE genes are shown. Star indicates significance p<0.05 (Wilcoxon rank-sum test).

Many cancers carried different *classes* of subclonal CNVs, allowing us to explore their relative effect on the transcriptional profile. In order to quantify how specific genomic changes affect the transcriptome, we analyzed pairs of subclones within a patient to determine the transcriptional and genomic distance between them. As in the earlier analysis, genetic distance was measured as the size in base-pairs of the genomic regions that differ between the pair of clones, while transcriptomic distance was estimated based on nearest-neighbor confusion (similar to CCI above, see *Methods* for details). For example, in MEL2 transcriptomic distance correlated well with genetic distance (**Fig 3d**). Generalizing this approach, we performed the pairwise analysis on all subclones in the dataset. In general, bigger genomic distance resulted in monotonically increasing transcriptional distance, although there were large individual variations. Importantly, while it was possible for two subclones to be transcriptionally highly distinct while being genetically similar, genetically distant subclones were never transcriptionally similar (**Fig 3e**). Measuring CNV events rather than genomic distance in basepairs showed the same trend (**Suppl. Fig 2b**). The pairwise analysis also allowed us to pinpoint whether the *class* of CNV influences the likelihood that it imposes a strong phenotypic constraint, by comparing pairs that differ in specific CNV classes. To do this, we classified clone pairs based on a hierarchical assessment of CNV differences. First, pairs with big differences in the number of copies (CN differences of four or more) were identified. The remainder were categorized based on if they had any sub-arm-level breakpoints, indicative of double-strand breaks, or only arm-level events which can result from chromosome mis-segregation. We found that high copy number changes had the largest and most consistent association with a transcriptionally distinct cell state, whereas transcriptional constraints associated with sub-arm level CNVs were heterogeneous, ranging from nearly-total separation to undetectable. Pairs harboring only arm level CNVs had weak transcriptional constraints (**Fig 3f, left panel**). In all clones where copy number differences were consistent with a WGD event separating the clones, cell states were highly constrained (**Fig 3f, right panel**), suggesting that WGD has specific phenotypic effects even if the *relative* copy number of most genes are not greatly altered. In line with these results, differential expression analysis between clones identified that the number of differentially expressed genes (DEGs) in pairs with only arm-level changes were low, (on average five DEGs between clones), compared to 19 and 43 for the sub-arm and highly amplified groups, respectively. Interestingly, the percentage of genes altered in *cis* followed a similar pattern, with the vast majority of DEGs mapping to non-affected regions for arm-level CNVs, while genes in *cis* usually made up more than half the DEGs in highly amplified clone comparisons (**Fig 3g**). Thus, our data is consistent with highly differing transcriptional impacts between CNV classes, where both *cis* effects on genes with directly altered copy number and secondary *trans* effects contribute.

### Gene dosage and tissue-dependent compensation

The direct effect of altering the copy number of a gene on its own transcript abundance (the *cis* effect) has been previously studied by performing mRNA-seq and WGS in parallel sections of a tumor or in cell lines. Using these systems, there is generally a positive gene dosage effect, meaning that expression levels tend to correlate positively with copy numbers, with some exceptions^13–15,33^. However, there are methodological issues with using bulk samples to study dosage response – for tumor biopsies, normal cell infiltration means that transcriptional profiles are a mix of cell moieties necessitating deconvolution-based strategies instead of direct quantitative inference^15,34^. In addition, by design these studies consider the dosage effect between distinct patients samples, which means that they are potentially confounded by differences in germline genetics and patient history. Using joint single-cell data removes many of these limitations. mRNA and CNV profiles are clean of any interfering normal or S-phase cells, and by restricting comparisons to subclonal differences the comparisons are made on a similar genetic, cancer site and cancer-historical context. To take full advantage of the possibility to make quantitative conclusions based directly on the single-cell data, we analyzed mRNA abundance of expressed genes as a function of copy state directly on the mean-normalized single-cell data, using robust linear regression and Pearson correlation (see *Methods* for details). A linear regression slope of 0 indicates that higher copy number is not associated with an increase in transcript abundance, indicating total dosage compensation, whereas a slope of 1 would indicate perfect additivity. Based on the regression analysis, the dosage effect was generally additive, with most slopes between zero and one (average of 0.65) (**Fig 4a**).

**Figure 4.**
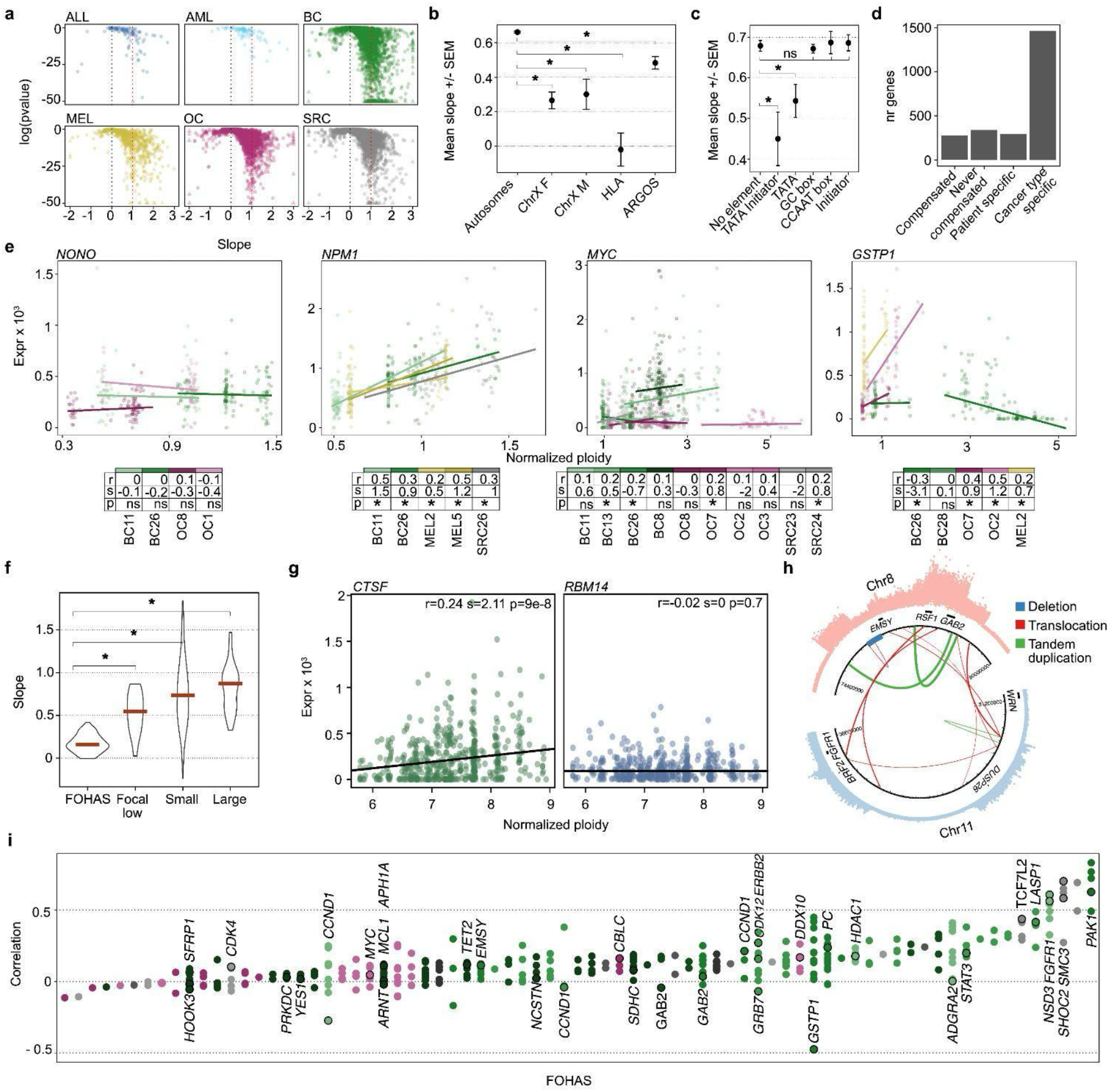
Within-patient dosage effects across six cancer types. **a**: Slope of transcript abundance as a function of copy number (normalized coefficient), from a robust linear model based on the single-cell data plotted against the Pearson p value. Cancer types are colored as in Fig 1d. Data is winsorized to the plotting area, with data outside the area indicated by triangles. **b**: Mean slope +/− standard error of the mean (SEM). Autosomes: all non-sex chromosome genes. Chr X F/M: genes on chr X in cells from female or male patients. HLA: human leukocyte antigen cluster genes, ARGOS: “Amplification-Related Gain of Sensitivity” genes previously found to be dosage compensated due to toxicity. Star indicates significance p<0.05 (Wilcoxon rank-sum test) **c**: Mean slope +/− SEM for genes grouped by promoter element or a combination of promoter elements. Star indicates significance p<0.05 (Wilcoxon rank-sum test) **d**: Gene classification based on the effect of gene dosage. *Compensated*: no dosage effect in any case, *Never compensated*: dosage effect in all cases, *Patient-specific*: compensation varies between patients within the same cancer type, *Cancer-type-specific*: compensation is consistent within cancer type. **e**: Dosage effects of individual genes exemplifying different categories. *NONO*: compensated, *NPM1*: never compensated, *MYC*: patient-specific, *GSTP1*: cancer-type-specific. Points shown are metacells made up of 5 nearest cells in ploidy space, with per-metacell average normalized CN and TPM. The slope of each line and Pearson correlation were calculated as in a, on the single-cell data. r=Pearson correlation, s=slope, p=Pearson p value. Star indicates significance p<0.05. **f**: Linear model coefficients of ploidy by gene expression. Star indicates significance p<0.05 (Wilcoxon rank-sum test) **g**: Expression and ploidy of *CTSF* and *RBM14* from a single FOHAS from BC26. r=Pearson correlation, p=Pearson p value and s=slope from robust linear regression. **h**: Structural variants in a FOHAS from BC4, determined by split-read mapping. **i**: FOHAS on x axis, color indicates which patient the segment originates from (key in Fig 1d), ordered by average Pearson correlation. Each dot is a gene on an individual segment, oncogenes and tumor suppressor genes from oncoKB are labeled in the order they appear from bottom to top.

Genome-wide analysis did not identify any large-scale local compensatory mechanisms on specific autosomes (**Suppl. Fig 3a)**. Chr X, however, showed a stronger dosage compensation (lower slope) than autosomes (**Fig 4b, Suppl. Fig 3a**), suggesting that sex chromosome dosage compensation mechanisms remain active also with aneuploid X chromosomes in cancer. Out of the autosomes, chr 6 had the lowest median slope, and the lowest median Pearson correlation. Rather than being uniform across the chromosome, this was driven by human leukocyte antigen (*HLA*) genes (**Fig 4b**), components of the major histocompatibility complexes that are responsible for antigen presentation to T cells^35^. *HLA* suppression, associated with reduced survival, is common across cancer types^36^ and often achieved through epigenetic silencing by DNA methylation^37^. We also analyzed if genes previously found to be associated with transcript dosage toxicity were dosage compensated in our data. Such so-called “Amplification-Related Gain of Sensitivity” (ARGOS) genes, which are located in commonly amplified regions but expressed at a lower level than expected^13^, had on average a lower slope than other genes, indicating partial compensation (**Fig 4b**). In contrast, we found no difference in dosage compensation between genes previously identified as oncogenes or tumor suppressor genes. (**Suppl. Fig 3b**).

Genetic elements of core promoters have been shown to dictate transcriptional behavior, with TATA or a combination of TATA and Initiator (Inr) core elements associating with larger transcriptional burst size^38,39^. We found that these same promoter elements also predict dosage sensitivity: genes that contain both a TATA and an Inr have an average slope of 0.45, and genes with only a TATA element 0.54, both significantly lower than genes with core promoters lacking these elements (slope=0.68, p < 0.005 and p<0.05, respectively). No other common promoter elements (GC-box, CCAAT-box), nor Inr alone, had a significant effect (**Fig 4c**). TATA-box promoters are generally associated with a high rate of transcription when active, and it is conceivable that such genes with a large capacity of continuous transcription are under stricter feedback control than lower capacity genes.

Among all genes consistently expressed at a robust level in our data set, we found four distinct dosage effect categories: fully compensated, never compensated, cancer-type-specific and patient-specific (**Fig 4d, Table 2**. Representative examples are shown in **Fig 4e, Suppl, Fig 3c, 3d**). Fully *compensated* genes are never detected as dosage sensitive (12% of total), *never compensated* genes are always dosage sensitive (14%).

Genes where compensated/sensitive behavior vary between patients within a cancer type are *patient-specific* (12%, among them the proto-oncogene^40^ MYC) and *cancer-type-specific* genes that were only sensitive to dosage in a subset of cancer types (62%). Thus, the majority of variation was between cancer types, suggesting that tissue-specific gene regulation networks could be a major cause behind differences.

Focal highly amplified segments (FOHAS) had the lowest slope of all segment types, indicating strong dosage compensation (**Fig 4f**). To confirm that the association holds true also in a non-linear setting, we trained a random forest classifier using expression data and found that FOHAS copy numbers were more poorly predicted, compared to other segment types (**Suppl. Fig 4a**). This might suggest a general mechanism such as that the repeated binding elements in FOHAS cause oversubscription of necessary transcription factors, or that specific genes in these regions are often under tight feedback regulation. We would expect that specific mechanisms are active for ARGOS genes. Indeed, in a FOHAS that contains the ARGOS gene *RBM14*^13^ and non-ARGOS gene *CTSF, RBM14*, was compensated for while *CTSF* was not (**Fig 4g**). FOHAS can be highly complex structures, with multiple tandem repeats and translocations (**Fig 4h**). We found that just as in the example of *RMB14* and *CTSF*, dosage sensitivity can vary widely within a FOHAS region (**Fig 4i**). Oncogenes identified on FOHAS also often varied between samples; for example, *MYC* and *CCND1* had highly patient-specific compensation patterns.

### Copy number alterations in non-cancer cells are rare and cell-type dependent

Somatic CNVs have been reported in non-cancer cells, both within the tumor microenvironment^41^ and in leukocytes where they increase in an age-dependent manner^42^. Our dataset includes non-malignant stromal populations collected together with the cancer cells, including fibroblasts, endothelial cells, and multiple types of immune cells (**Fig. 5a**). We found chr X to be the most frequently aneuploid chromosome in the non-cancer cells, with loss enriched in T-cells and mural cells across patients, and gains in endothelial cells detected at high frequency in one patient (**Fig 5b, Suppl. Fig 5a**). Fibroblasts in the tumor microenvironment have previously been associated with CNVs, and selection of cells with a gain of chr 7 has been reported^41,43^. We found CNVs in fibroblasts in multiple samples, with chr 12 gains found in two patients, chr 7 gain in four patients and chr 18 gain in three patients **(Suppl. Fig 5b**). X chromosome loss in our data was associated with a complete lack of XIST expression, indicating that the cells respond to loss by lifting X-chromosome inactivation. We also found a positive correlation between the number of X chromosomes and XIST expression with a slope of 0.67 (**Fig 5c**), in line with silencing of additional copies. Importantly, the dosage effect in non-cancer cells was similar to cancer cells, with the exception of chr X in female cells – as expected given chr X inactivation (**Fig 5d**). This precludes the possibility that non-cancer CNVs are the result of rare method artifacts and suggests that the alterations have an effect on cell state, potentially leading to selection. Larger-scale studies specifically aimed towards non-cancer cells will be needed to fully explore CNV-dependent clonal selection in non-cancer cells.

**Figure 5.**
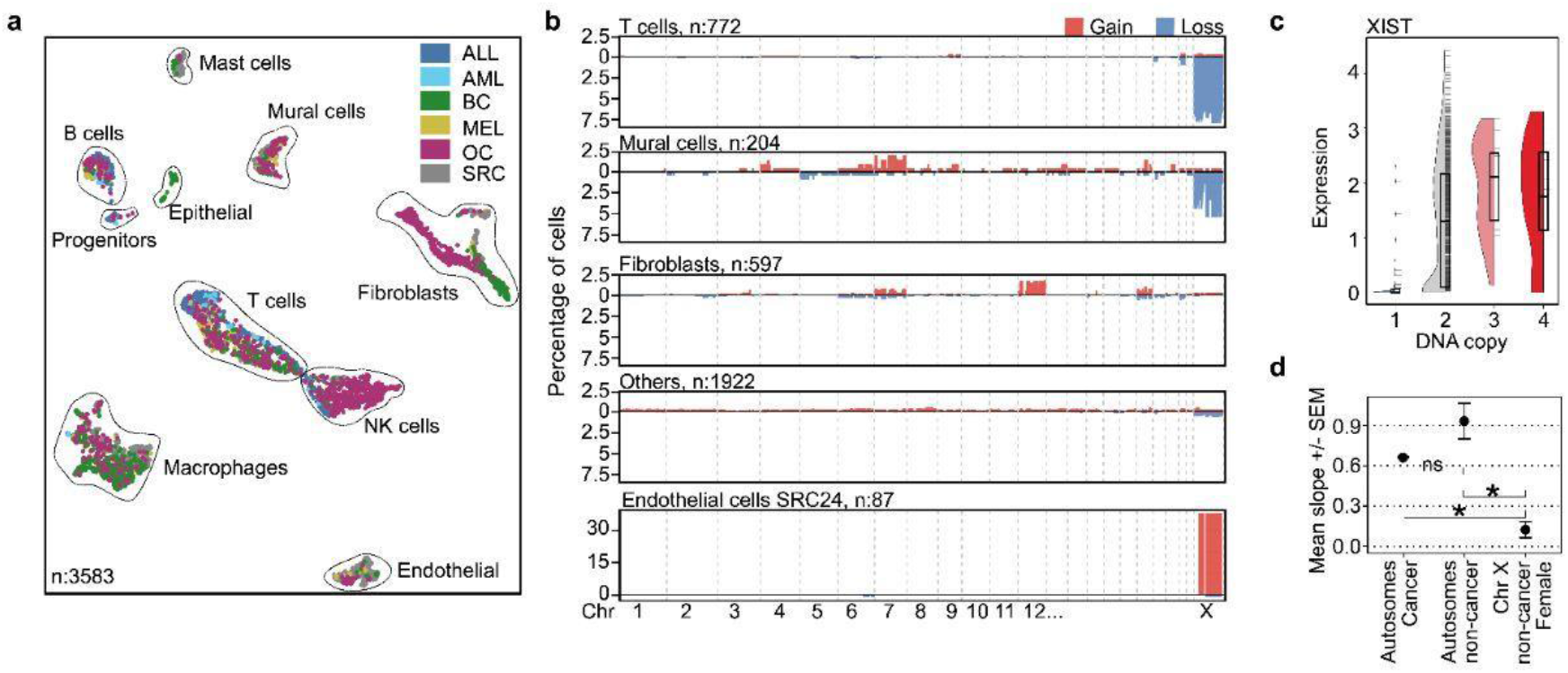
CNVs in non-cancer cells. **a**: UMAP of 3583 cells that did not belong to any cancer subclone based on scWGS. Color indicates cancer type. **b**: CNVs identified in non-cancer cells, by mRNA-based classification. **c**: Expression of XIST in all female non-cancer cells, split by copy number of chr X. Half-violin plot shows density, box extents indicate 25th and 75th percentiles with the central line indicating the median. Individual data values are shown as horizontal lines to the right of the density plot. **d**: Mean slope +/− standard error of the mean (SEM) for genes on autosomes from cancer cells and non-cancer cells as well as chr X in non-cancer female cells. Non-cancer cell gene dosage is calculated within each cell type. Star indicates significance p<0.05 (Wilcoxon rank-sum test).

### Transient clonality is a common feature of solid tumors

While most of the patients in our dataset presented a clear and hierarchical subclonal structure, ten samples lacked any detectable subclones. These included six soft tissue SRC (three myxofibrosarcomas, two leiomyosarcomas and one undifferentiated pleomorphic sarcoma), one BC (triple negative breast cancer) and three OC. The samples were categorically different from the rest in two critical ways: genetic heterogeneity was extremely high, with majority of cells having unique copy numbers in a chromosome-arm sized region or more (>50 million basepairs) (**Fig 6a**), and they had undetectable or very weak CNV-based subclonal hierarchy, i.e. CNVs were not stably inherited to form subclones. We named this type of subclonal behavior *transient clonality*. Single cells with transient clonality varied in large chromosomal regions such that the median difference between cells in transient cases was of similar magnitude or larger than the median distance between subclones in non-transient cases (**Fig 6b**). Transient clonality was not related to treatment-associated DNA damage, since the transient samples were all treatment naive. Further, they included both primary and metastatic lesions; the SRC and BC were primary tumors, while the OC samples were taken from a metastatic location (omentum or ascites).

**Figure 6.**
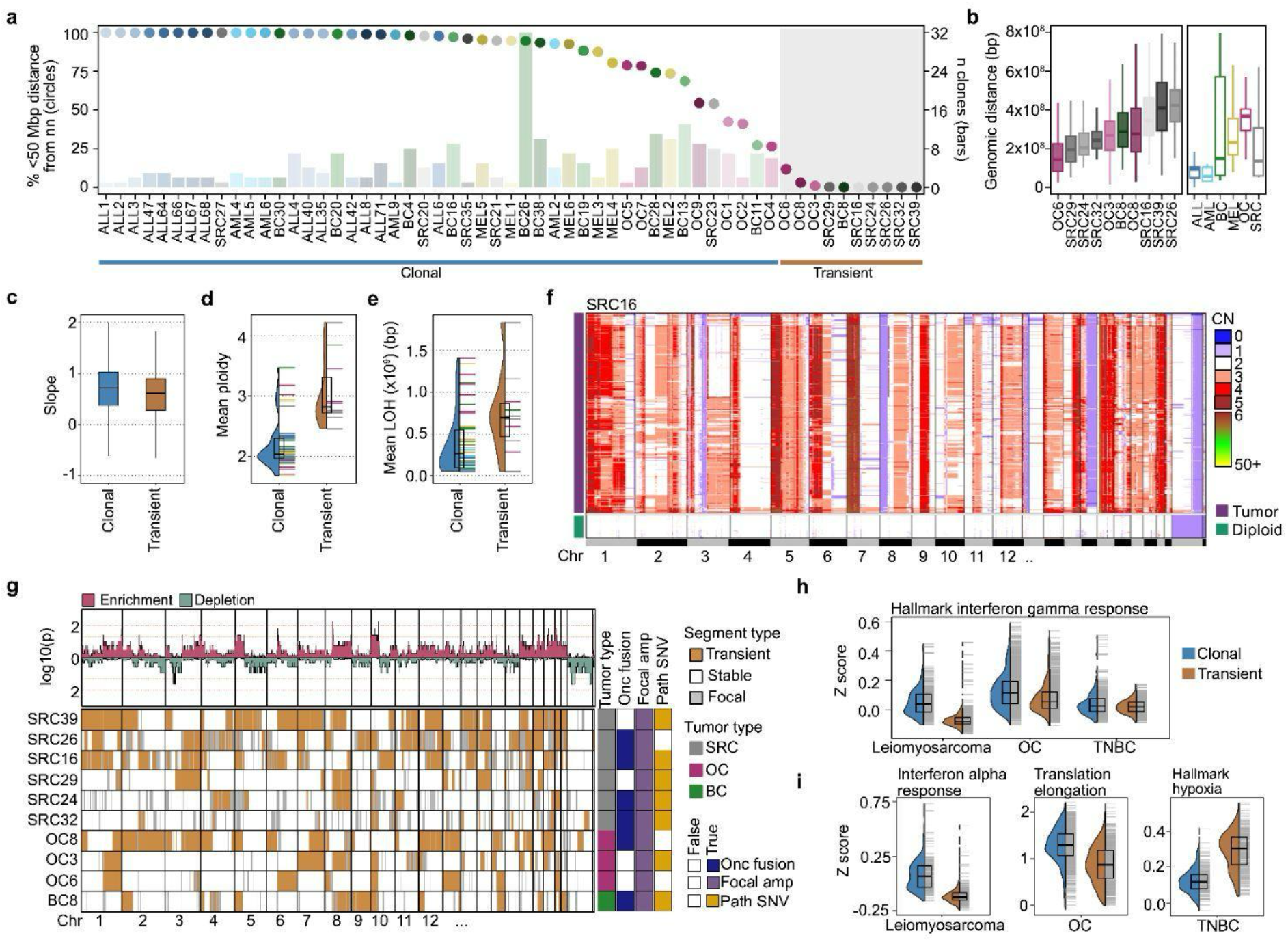
Transient clonality in solid tumors. **a**: Left y axis (circles): Percentage of cells whose distance to its nearest-neighbor cell in genetic space, is less than 50 Mbp (megabase pairs). Right y axis (bar): Nr of clones per patient. Color indicates patient. **b:** Distance between nearest-neighbor cells in genetic space, across patients. Plot shows median with boxes indicating 25th to 75th percentiles, and whiskers 1.5x the interquartile range. Left panel: Transient cases. Right panel: Genetic distance to the nearest-neighbor from a different subclone, by cancer type. **c**: Dosage effect (slope) in clonal and transient cases. Boxplots as in b. **d**: Average patient ploidy for transient and clonal patients. Half-violin plot shows density, box extents indicate 25th and 75th percentiles with the central line indicating the median. Horizontal lines on right side of density indicate individual values of average ploidy, colored by patient (as in a). **e**: Average genetic size of loss of heterozygosity in transient and clonal cases. Plot elements as in d. **f**: Single-cell CNV profiles from a transient sarcoma (SRC16, n=636). Heatmap shows each single cell as a row and copy numbers are indicated by color. Aneuploid and diploid cells are separated as blocks. **g**: Genome wide heatmap labeling transient (>20% of cells have a CN in that segment differing from median) and stable segments in orange and white respectively, and focal segments (<5Mb) in grey. Patients are annotated by presence of oncogenic fusions, a highly amplified focal CNV and a pathogenic SNV from the cosmic database. Bar plot (top) shows statistical significance of enrichment/depletion of transience compared to mean across the genome. Yellow line denotes p=0.05, red line p=0.01 (Poisson density). **h**: Hallmark interferon gamma response z scores for tumor subtypes split by genotype. Half-violin plot shows density, hinges indicate 25th and 75th percentiles, and the central line the median. Horizontal lines on the right side of the density plot indicate z-scores of individual genes. **i**: Z scores of Hallmark and Reactome pathways split by tumor subtype. Plot elements are as in h.

Using the joint mRNA-seq data, we analyzed dosage effects in transient cases and confirmed that they were comparable to clonal cases. (**Fig 6c, Suppl. Fig 6a**). This implies that copy number alterations in these chaotic genomes are likely to carry some degree of functional consequences, and positions them as being part of a class of CNV-driven lesions identified previously by scWGS^44^. Given the extremely high copy number variability, we would expect there to be a high degree of gene copy redundancy. In line with this, we found that the transient tumors had high average ploidy. Interestingly, the transient cases also had a high degree of loss of heterozygosity (LOH), with LOH regions often encompassing a majority of the chromosomes where it could be confidently called (**Fig 6d, 6e**). A plausible explanation for this observation is that transient cases evolved first by large-scale LOH, followed by one or more rounds of WGD, both predating the ongoing extensive chromosome missegregation.

While overall variation was very high, the transient cases did exhibit a varying degree of stability across chromosomes, and each sample had a clear underlying clonal structure, as exemplified for SRC39 (**Fig 6f**), OC8 and BC8 (**Suppl. Fig 6b, 6c, extended data figure**). Most of the transient cases harbored oncogenic fusions and pathogenic SNVs and all of them carried FOHAS. WGS data was available for three of the six transient sarcoma cases, and of those, two had a mutation in *TP53*. Amongst the remaining three transient sarcomas, we found two with *TP53* mutations by analyzing mRNA data. Thus, it seems that *TP53* mutation is not an absolute prerequisite to develop transient clonality. All chromosomes were affected by the transiency, but relative frequency suggested that some, such as chr X and chr 9, were less likely to be affected (**Fig 6g**). It could be speculated that the apparent genetic chaos reflects unsuccessful cell divisions, where most of the affected daughter cells are already marked for death. Therefore, we first tested if the cells were G2M arrested, if they were apoptotic, or with activated unfolded protein response pathways which could indicate heavy stress. However, there was no significant evidence of increased activation of any of these pathways, in line with transient clonality representing a stable phenotype rather than a short-lived cell state (**Suppl. Fig 7a, 7b**).

To investigate if a specific gene signature accompanied the transient cases, we performed pathway analysis. Since the tumor types where clonal transience appear are very different, a direct comparison across all samples is problematic and we therefore split the analysis by subtype (myxofibrosarcoma and pleomorphic sarcoma were left out of this analysis, since they all had transient clonality). Few pathways were shared between the subtypes, but among the significant hits were the *GOBP viral genome regulation* pathway and *hallmark interferon gamma response* (downregulated in transient compared to clonal) (**Fig 6h, Suppl. Fig 7c, Table 3**). This might indicate either an immune-cold environment or immune evasion within the transient genotype, which would be in agreement with ovarian cancer literature which has found WGD to be associated with immunosuppressive phenotypes^45,46^. In addition to the shared pathways, we found several subtype-specific effects, with *downregulation in interferon alpha response* and other immune related pathways enriched in transient leiomyosarcoma compared to clonal (**Fig 6i, Suppl. Fig 7d**). Multiple pathways related to translation were downregulated in transient OCs but upregulated in the other two transient subtypes (**Fig 6i, Suppl. Fig 7e**). The most significant triple negative breast cancer pathway was response to hypoxia, while other pathways identified included immune related pathways (**Fig 6i, Suppl. Fig 7f**). In summary, the results suggest that subtype-specific differences might exist between clonal and transient genotypes. Future work with larger targeted datasets will be needed to fully understand the causes and functional consequences of clonal transience.

## Discussion

In this study, we present the largest and most diverse joint single-cell mRNA and WGS analysis to date, examining how CNV-based subclones constrain cellular phenotypes in patient samples, across six different types of human cancer. The scale and wide variety of cancer types enables us to make general inferences of how structural variation shapes transcriptional state. Our data revealed substantial within-patient variability in how subclonal CNVs affect the transcriptome. Part of this variability can be directly attributed to the nature of the CNV classes. High-level amplifications tended to impose wide-reaching constraints in *cis* and *trans*, whereas arm-level CNVs had a very weak effect, often at a level below the intrinsic transcriptional variability between cells. However, even though their overall effect on transcriptional phenotype was weak, arm-level CNVs still affected mRNA abundance in an approximately additive fashion. Even though they could be described as passenger events in the sense that they do not drive the appearance of a novel, distinct, phenotype they are not silent and it is possible that they provide subtle phenotypic heterogeneity that affects disease outcome. We also find that large differences in ploidy, likely arising from subclonal WGD, markedly affect cell state, indicating that genome-scale structural changes have phenotypic consequences even if the relative differences between gene loci are similar before and after WGD.

The observed transcriptional constraints can be attributed to a combination of *cis* and *trans* regulatory effects, with their relative contributions depending on the underlying CNV class. When assessing dosage sensitivity, the *cis* effects of CNVs, we find that most genes exhibit some degree of tissue-specific buffering. Overall, we see a highly heterogeneous response to copy number alterations. Some of this heterogeneity might stem from missing information inherent in our experimental setup – we do not have the power to reliably identify subclone-specific SNVs or gene fusions and it seems likely that some of our unexplained variation stems from such undetected variation. Deeper sequencing along with improved computational analysis methods could help reduce the missing information. Further large-scale, cross-tumor studies are clearly needed to fully address the limited generalizability of dosage effects.

We also found rare occurrences of large-scale CNVs in non-cancer cells. Chromosomal aberrations were highly cell-type-specific with the most common alteration, loss of chr X in T-cells, occurring in 7% of the cells across several patients and cancer types. Additional aneuploidies occurred in endothelial cells and fibroblasts, but at much lower rate. Interestingly, we found several cases of patient-specific gains (in endothelial cells and fibroblasts). Aneuploidies in autosomes were highly dosage sensitive, with the exception of chromosome X, where increasing XIST levels suggest that X chromosome inactivation mechanisms counteracted gains/losses. Taken together, these results implicate different mechanisms between chr X loss and other aneuploidies. Loss of an inactive X chromosome would likely have minimal impact on cellular phenotype and its prevalence in many patients suggests that it is a relatively common event. On the other hand, patient-specific gains of autosomal chromosomes, with concomitant increase in transcript levels, suggests that selection by clonal expansion has occurred. This expansion must have been separate from the tumor cell expansion, since the patient-specific aneuploidies in non-tumor cells were distinct from CNVs in the tumor clones.

A large subset of the solid tumors in our data set presented with *transient*, rather than *stable* clonality. These cancers were common among soft tissue sarcomas and ovarian cancers and were characterized by a complete lack of clonal structure, such that no cells share a CNV profile, and that the differences between nearest-neighbor profiles is of a similar or larger scale as differences between stable subclones in non-transient cancers. Theoretically, these cases could be the result of undersampling cells from a tumor with an ultra-high number of stable clones. However, we think this explanation is unlikely. First, the number of stable clones necessary to not see any cell pairs stemming from the same clone is incredibly high (in the order of tens or hundreds of thousands). Second, the individual cells are not organized in a similar manner to subclones in a complex cancer. Rather than showing sequential gains in a hierarchical fashion, cells from clonally transient cancers tend to all follow a major copy number profile, with large gains and losses superimposed and in a seemingly random fashion. Third, the breakpoints tend to remain the same among the cells in a transient cancer, with most changes due to gains or losses of common genomic fragments. Because of these features, we suggest that transient clonality is mainly due to continuous missegregation of chromatin fragments at mitosis. Importantly, while these tumors lack a defining transcriptomic profile, their fluctuating copy-numbers still produce dosage effects at a similar level to that seen in stable clones, indicating that they are not functionally silent.

Taken together, our study demonstrates the feasibility and value of large-scale joint scWGS/mRNA-seq. We show that genetic constraints on phenotype depend strongly on the class of CNVs, with small (megabase-size) segments, that are not detectable using RNA-based inference, having particularly strong effects. We further identify tissue-specific and core-promoter dependent compensation of dosage effects. We identify transient cancers, where most cell divisions lead to non-silent errors in segregation. These findings collectively highlight the importance of direct measurement of DNA and mRNA in the same cells to understand genotype-phenotype associations in cancer.

## Methods

### Sample preparation

This study was performed in accordance with the ethical standards of the Helsinki Declaration with written informed consent from patients or guardians. Bone marrow of leukemic patients was collected at diagnosis. Red blood cells were depleted by gradient separation or red cell lysis before the samples were viably frozen in freezing medium (FBS supplemented with 10% DMSO). Samples from solid tumors were collected following surgery. Samples except for OC6-9 were cut into small pieces and single-cell suspension was prepared by enzymatic and mechanical dissociation using a tumor dissociation kit (Miltenyi, cat 130-095-929). Solid tumors were treated with DNAseI for 5 minutes. Cells were viably frozen in freezing medium following dissociation. OC6-9 samples were dissociated to single cells and frozen as described previously^47^.

### Flow cytometry sorting

Tumor samples were thawed and washed in 1xPBS. Viability and cell number were assessed in a cell countess. Samples with few viable cells or samples containing a lot of debris underwent dead cell removal by magnetic beads (Miltenyi kit). Single-cell suspensions were finally filtered on a 40*µ*M strainer, centrifuged at 350g for 5 min and resuspended in cell staining buffer (1x PBS with 2% FBS) at a concentration of 1-5 million cells/mL. Solid tumors were stained with CD45 (FITC) for negative selection according to manufacturer’s recommendations, ovarian cancers 6-9 and breast cancers 28,30 and 38 were additionally stained with EpCAM (AF700). After the antibody was washed away, 1*µ*L eFluor450 viability dye was added per 500*µ*L cell suspension and incubated on ice for 20 minutes. Ovarian tumors 1-5 were stained with CD3, CD56, CD45 and a viability dye (APC-Cy7) with the same protocol. ALLs were depleted of dead cells by magnetic bead separation and stained with CD34, CD10, CD20, CD38, CD45, and CD19 but no viability dye. AMLs were stained with CD19, CD3 (APC both of those - neg selection), CD34 and CD45, and a viability dye (eFluor450).

Gates were set to exclude debris and doublets, remove dead cells where a dead cell marker was used and to enrich for tumor cells according to previously known markers. Cells were sorted into 384-well plates containing 3uL lysis buffer composed of oligodT, dNTPs, ERCC controls, Triton X-100 and RNAse inhibitor. Plates were spun down at 2000 x g at 4°C for 5 min and snap frozen on dry ice and kept at −80°C until they were processed.

### DNTR-sequencing

DNTR protocol was performed as previously described^48^. Briefly, plates were thawed and spun down at 2000 x g at 4°C for 5 min to allow separation of the nuclear and cytosolic fraction. The cytosolic fraction was aspirated carefully and moved to a second plate. The cytosolic fraction was then either snap frozen or processed immediately by annealing the oligodT to mRNA molecules, reverse transcribing and cDNA amplification. cDNA was purified with magnetic bead purification, cDNA was tagmented, barcoded and further amplified. The nuclear fraction is treated with proteinaseK to remove histones, then tagmented, barcoded and amplified. Amplified libraries were purified using magnetic beads and sequenced. Paired end sequencing was performed, either 2x37bp or 2x150bp on nextseq550 or novaseq6000 (**Table 1**), targeting approximately 100M sequenced bases per cell.

### DNA data processing

DNA data processing was performed with ASCENT^32^ (github.com/EngeLab/ASCENT). Briefly, reads were trimmed with trim-galore to remove adapter sequences and low-quality bases. Reads were aligned to GRCH38 using bwa^49^ and duplicates were removed using Picard (https://github.com/broadinstitute/picard). Bins were filtered to remove problematic regions and cells that were confirmed to be in S or G2M phase by mRNA expression were removed. Cells were segmented at 40kb resolution using mpcf^50^. Scale factor estimation was performed, and initial clones were assigned based on the scaled copy number profiles. These clones were refined manually (see https://github.com/EngeLab/SubclonalGeneDosage) at 10kb resolution and allele specific copy numbers were calculated per clone-segment.

### RNA data processing

RNA data processing was performed with ASCENT^32^ (github.com/EngeLab/ASCENT). Briefly; reads were trimmed with trim-galore to remove adapter sequences and low-quality bases. Reads were aligned to GRCH38 using STAR^51^ (v2.7.7a). Duplicates were removed and reads were counted using featureCount. Quality control filters were applied to exclude low-quality cells: cells with fewer than 10,000 reads or 500 detected features were removed. Additionally, cells falling below the 0.005 quantile cutoff for ACTB expression were excluded from further analysis.

## Analysis

### RNA visualization

Count data was scaled and clustered based on the top 2000 variable genes, with 40 neighbors and resolution 0.5 to create a UMAP with 40 dimensions for the entire dataset using Seurat^52^ (v 5.3.1). For patient specific UMAPS cells from aneuploid clusters, based on genetic information, were selected and 20 nearest neighbors were found with resolution 1 to create a UMAP from the 20 first dimensions.

### Linear models to discern effects of different variables

Expression data of aneuploid cells originating from patients harboring at least two subclones was used to build linear models. The 1000 highest expressed genes across the dataset were analyzed and first a model with only cancer type as a covariate to explain the expression levels per gene per cell was built. Secondly cancer type and patient were used as covariates, and thirdly a model with cancer type, patient and clone as covariates. The adjusted R squared value was used for visualization.

### Transcriptomic distance

CCI scores were calculated per clone using the top 3000 most highly expressed genes per patient. Only clones composed of at least seven cells that passed mRNA quality control were included in the analysis. For each clone, 100 iterations were performed on log transformed mRNA counts, sampling five cells per clone. Pairwise distances between cells were computed based on Spearman correlation of gene expression and these distances were used to perform multidimensional scaling (MDS). For each cell the nearest neighbor was identified and compared to the expectation of a random nearest neighbor. The CCI score was calculated as log(observed/expected). For pairwise comparison the same procedure was performed, except between only two clones at a time.

### Analysis of fusion genes and SNVs in single-cell RNA-seq data

Fusion genes are identified by merging all high-quality RNA bam files per patient and running STAR-fusion^53^ (v1.13.0). The output is filtered to only include fusions that are at least 5Mb apart in the linear genome, where read evidence comes from more than 10 cells and a maximum of 1 of those cells being classified as diploid. If one of the genes in the pair is oncogenic the fusion was said to be oncogenic. SNVs were identified from mRNA by merging bam files of all tumor cells and identifying SNVs in the genome with minimum read quality 20. The output was filtered for common SNVs (1000G phase3 common SNPs (AFe4)). The candidate SNVs were categorized by running ensemble’s variant effect predictor (VEP)^54^ with buffer size 500, distance 5000, polyphen b, sift b and transcript_version. The VEP output was filtered to include only Missense_variant, stop_lost, stop_gained, transcript_ablation or NMD_transcript_variant, and alternative allele readcount be a minimum of 16 from 8 different cells. Finally, to identify pathogenic SNVs we included SNVs found in the COSMIC database that were annotated as either “possibly pathogenic” or “pathogenic” by CLIN_SIG information.

### Analysis of complex structural variants

Candidate structural variants were called by running manta^55^ (v 1.6.0) on pseudobulk bam files of all aneuploid cells per cancer. Output was filtered by removing events with less than 5 reciprocal spanning reads. Links in the predicted amplicon region and associated genomic regions were plotted, with coverage calculated directly from the merged (pseudo-bulk) scWGS. Translocations and tandem duplications were visualized as arcs with a line width proportional to the relative number of reciprocal reads, deletions were visualized as a thick line across the affected region.

### Gene dosage

Dosage was calculated at the single-cell level. Single-cell CN, derived from refined breakpoints, were mean-normalized and single cell transcriptomic data was normalized as counts per million (CPM). Per patient transcriptome filtering required genes to have an expression level >100 cpm and to be expressed in more than half the cells. Genomic per patient filtering required that on raw CN data the gene needs to exist in at least two distinct copy states in more than 8 cells and on normalized CN data, the difference between the highest and lowest copy number had to exceed 0.2. Outlier CNs were removed based on the upper 0.99 and lower 0.01 quantile. Finally only genes where the CN variance > median variance per cancer type were retained. Robust linear regression with huber loss were modeled per gene per patient and the coefficient normalized by dividing the average expression over average ploidy per patient gene. Pearson correlations were calculated on the same normalized dataset. Dosage was additive with a significant Pearson p value (<0.05) and a positive (>0) Pearson correlation. Promoter element information was downloaded from the Eukaryotic Promoter Database (EPD)^56^, the oncogene and tumor suppressor genes were annotated with the OncoKB cancer gene list^57,58^ available at (https://www.oncokb.org/cancer-genes) and ARGOS gene list was downloaded from the publication^13^.

### Highly amplified focal CNVs

Segments smaller than 5Mb with a mean normalized max CN > 15 and average mean normalized CN > 8 were classified as highly amplified focal CNVs. For comparison, segments smaller than 5Mb with a mean normalized max CN <15 and average <4 were annotated as focal lowly amplified segments. To predict the CN of highly amplified focal CNVs from gene expression, a random forest model was implemented, using the ranger (v 0.17.0) R package with settings num.trees=1000, importance=“none”, scale.permutation.imporatnce=TRUE, mtry=NULL. The model was trained on the same set of genes as were used in the gene dosage calculations.

### Non-cancer cell analysis

Non-cancer cells were identified from pre-refined clusters on the patient level. mRNA corresponding to non-cancer cells was clustered and cell types annotated based on expression of key markers. Non-cancer cells were annotated as harboring CNVs if they had an aneuploid segment larger than 5Mb. Dosage calculations were performed in the same way as for cancer cells, except the threshold for a minimum number of cells with said CNV was set to four (nine in cancer dosage analysis).

### Genetic distance

Hamming distances were calculated between all cells on the CN x bin matrix. Based on the resulting distance matrix the nearest neighbor was identified for each cell. To obtain a tumor-wide measure of genetic heterogeneity the same analysis was done, excluding the cells own clone and averaging the distances to all other clones.

### Classification of segments in transient patient samples

Segments were classified as transient if they were >5Mb and if at least 20% of the cells had a different CN than the average CN of that segment. To calculate which segments were enriched/depleted for being transient we calculated Poisson probabilities.

### Hallmark pathway analysis

Differential expression analysis was performed per tumor type or per tumor subtype on all aneuploid cells, split by if they were clonal and transient by running Seurats FindMarkers with logfc.threshold=0 and min.pct=0.1. The resulting genes were ranked based on average log2 fold change and the R package fgsea^59^ (v1.28.0) was used to perform enrichment analysis. All pathways that were not found in the subcollections CGP, IMMUNESIGDB, VAX or collections C1 and C8 from The Molecular Signature Database (MSigDB)^60^, which were downloaded using the msigdbr (v 25.1.1) R package, were used. The results were filtered for significance (adjusted p value <0.05) and ordered by normalized enrichment scores (NES). Shared pathways were those that had the same directionality of NES and were found significant in all clone vs transient comparisons across tumor types. For subtype-specific analysis only the Hallmark, REACTOME and KEGG pathways were kept.

## Data Availability

Raw data for all samples except for OCs will be available under restricted access on https://doi.org/10.48723/gr42-ja30 And OCs on EGA with ID: EGAS50000001643 Processed data and analysis scripts are available on github (https://github.com/EngeLab/SubclonalGeneDosage).

## Ethical declaration

This study was approved by the Regional Ethics Committee in Stockholm, Sweden (Dnr 2021-02718, 2019-05149, 2019-01805, 2022-05409-01, 2020-04163, KI 76-51, KI97-451), and Regional Ethics Committee in Pirkanmaa, Tampere Finland (#R13109). Ovarian high-grade serous carcinoma patients OC6, OC7, OC8 and OC9 were enrolled in DECIDER trial (**ClinicalTrials.gov** ID: NCT04846933) with an approval from the Ethics Committee of the Wellbeing Services County of Southwest Finland (VARHA/28314/13.02.02/2023)..

## Acknowledgements

Part of this work was facilitated by the Protein Science Facility at Karolinska Institutet, Stockholm and BEA, the Bioinformatics and Expression Analysis core facility, which is supported by the board of research at the Karolinska Institutet. This work was supported in part by the Swedish Research Council (2023-02912), the Swedish Cancer Society (232839 Pj), Swedish Childhood Cancer Foundation (PR2024-0108 and MT2023-0014), Sigrid Jusélius Foundation (AV), K. Albin Johanssons stiftelse (AV, MF) and Cancer Foundation Finland (AV), the European Union’s Horizon 2020 research and innovation programme under Grant Agreement No. 965193 for DECIDER and the European Union (ERC-CoG STRONGER 101125261 for AV). Views and opinions expressed are however those of the author(s) only and do not necessarily reflect those of the European Union or the European Research Council. Neither the European Union nor the granting authority can be held responsible for them. We want to thank the patients and their families and the clinicians and other hospital personnel that participated in the sample collection.

## Author contributions

ME designed the study with help from SK. ME, VZ and SK developed the experimental and computational framework for analysis. SK, VZ, MY, LB, CS, HC processed samples. XC, YL, SEB, DS, YZ, AV, NH, JH, HH, FHF, JH, OL, MH, SH and MMF provided samples and provided cancer type-specific biological insight. SK processed the data. SK and ME analyzed and interpreted the data and drafted the manuscript with input from all authors. All authors reviewed and approved the final version.

