## Extended data figure for "Principles of subclonal gene dosage across human cancer"

ALL1

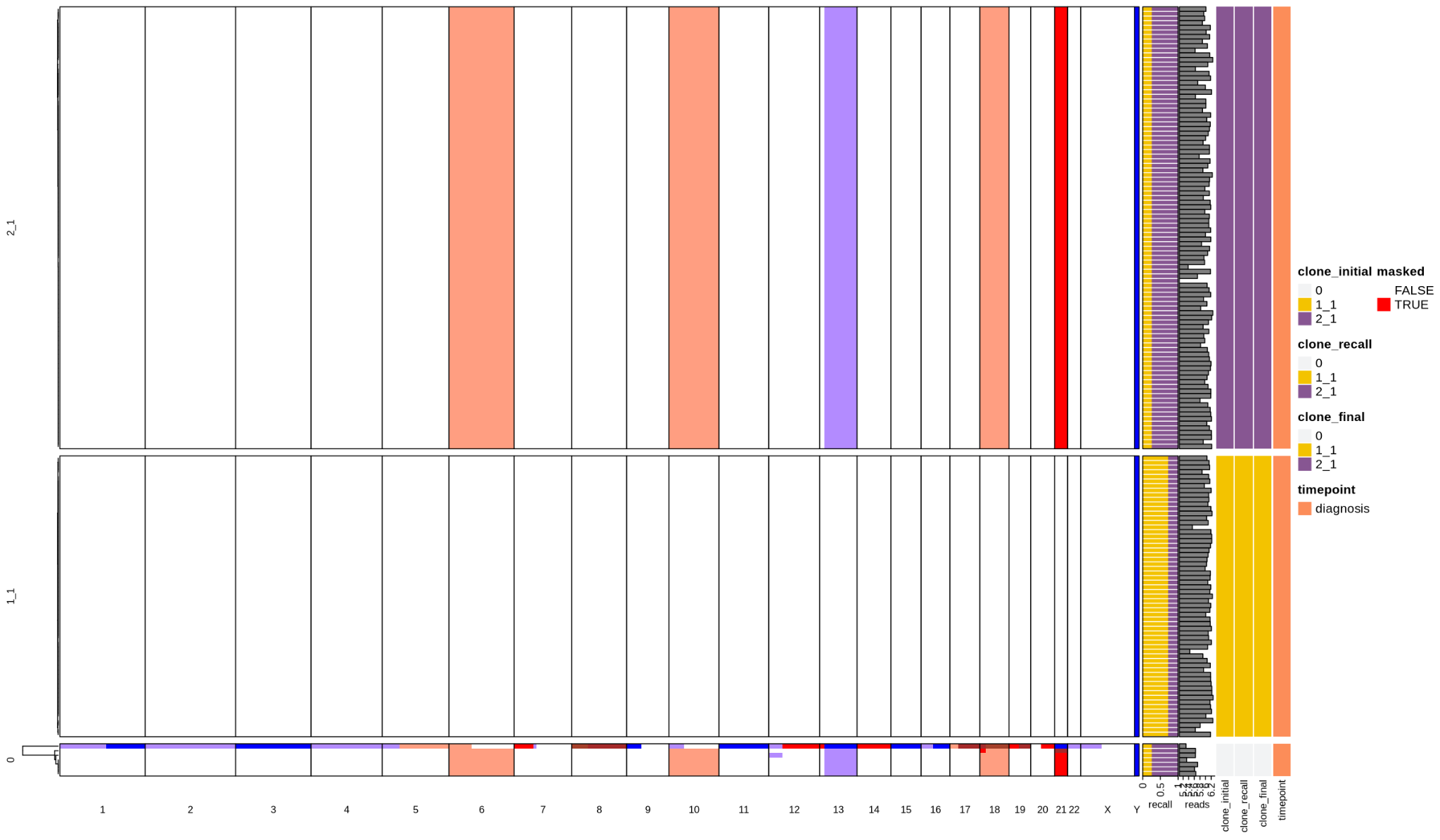

ALL2

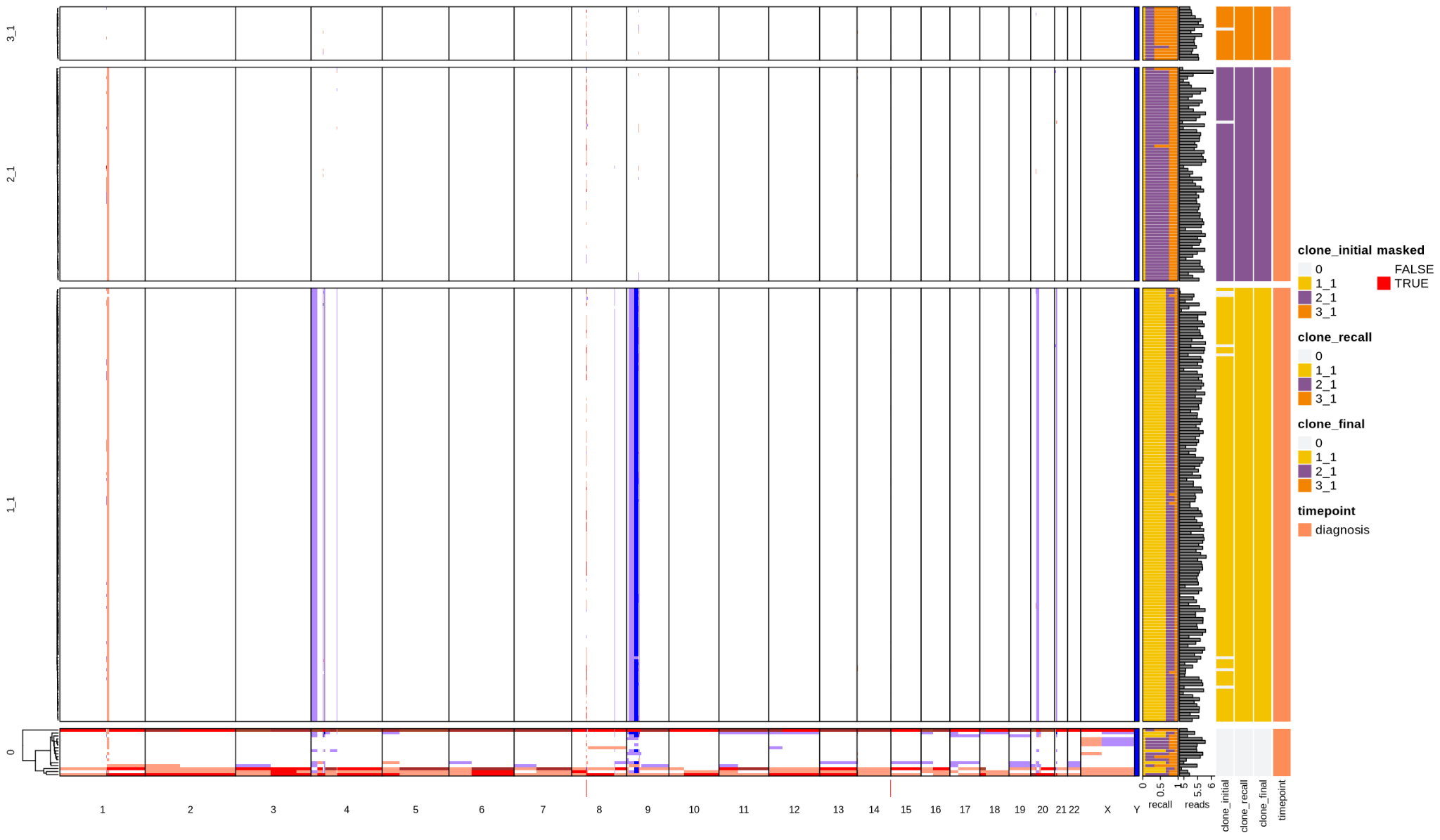

ALL3
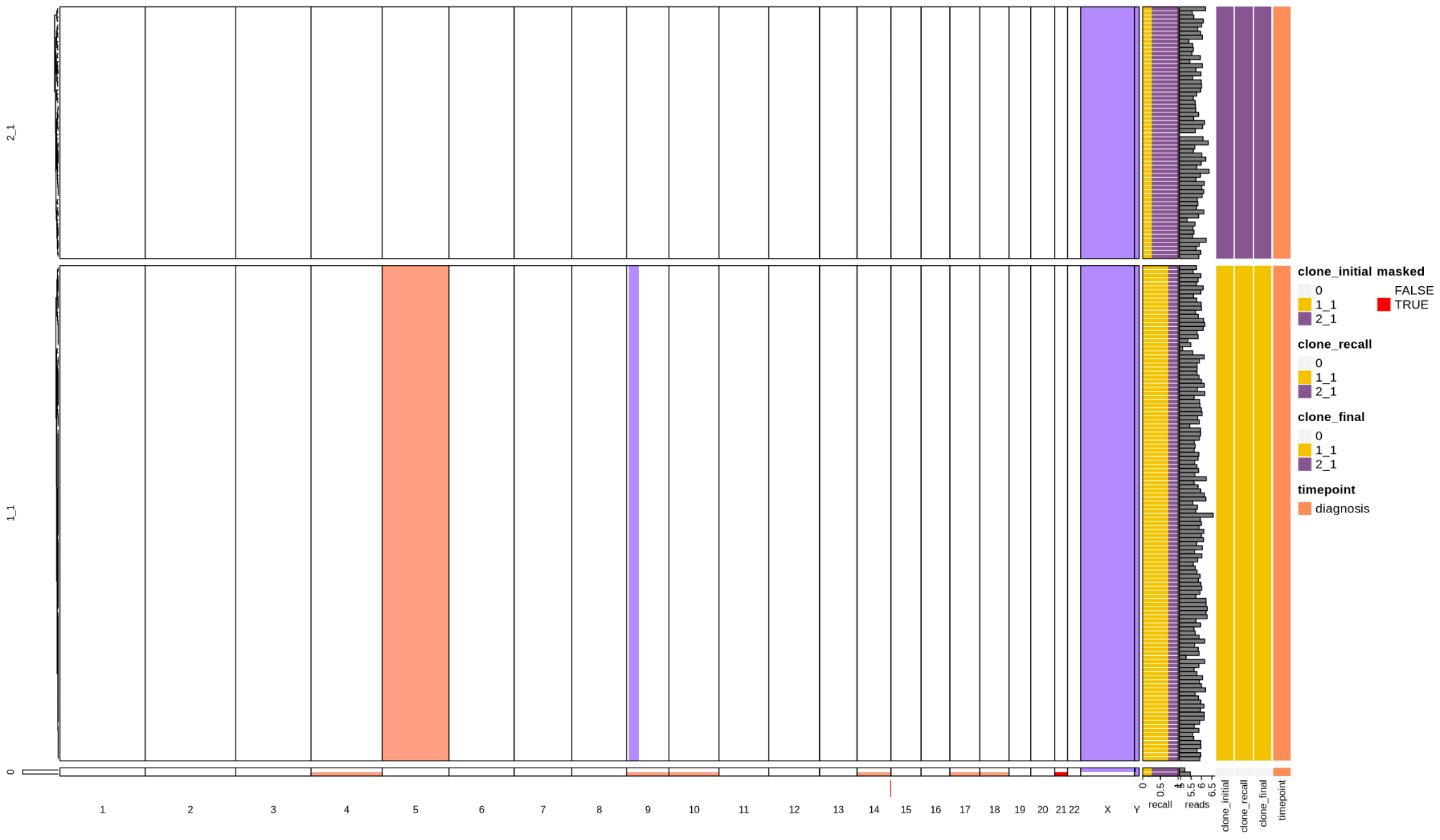

ALL4

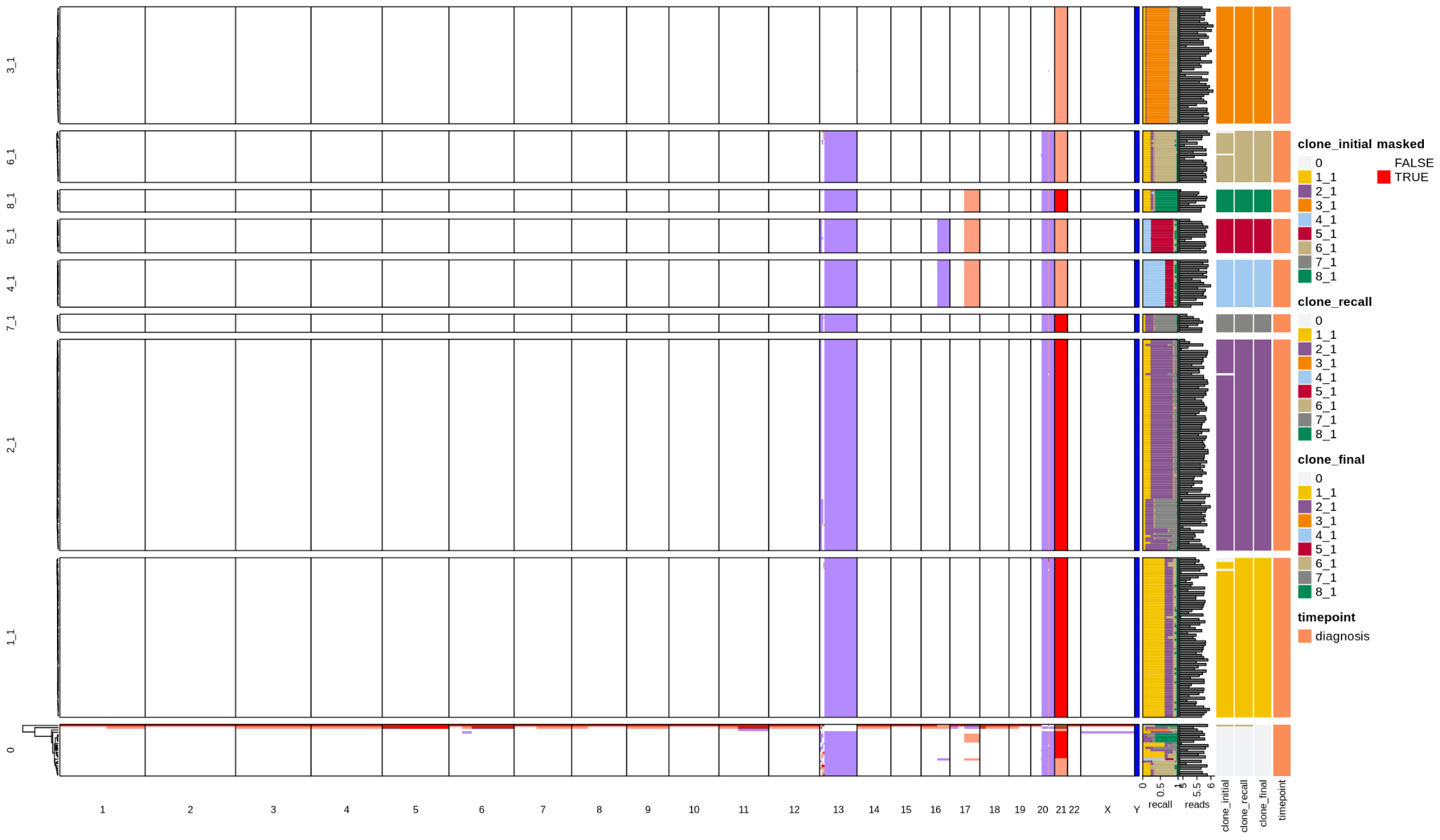

ALL6

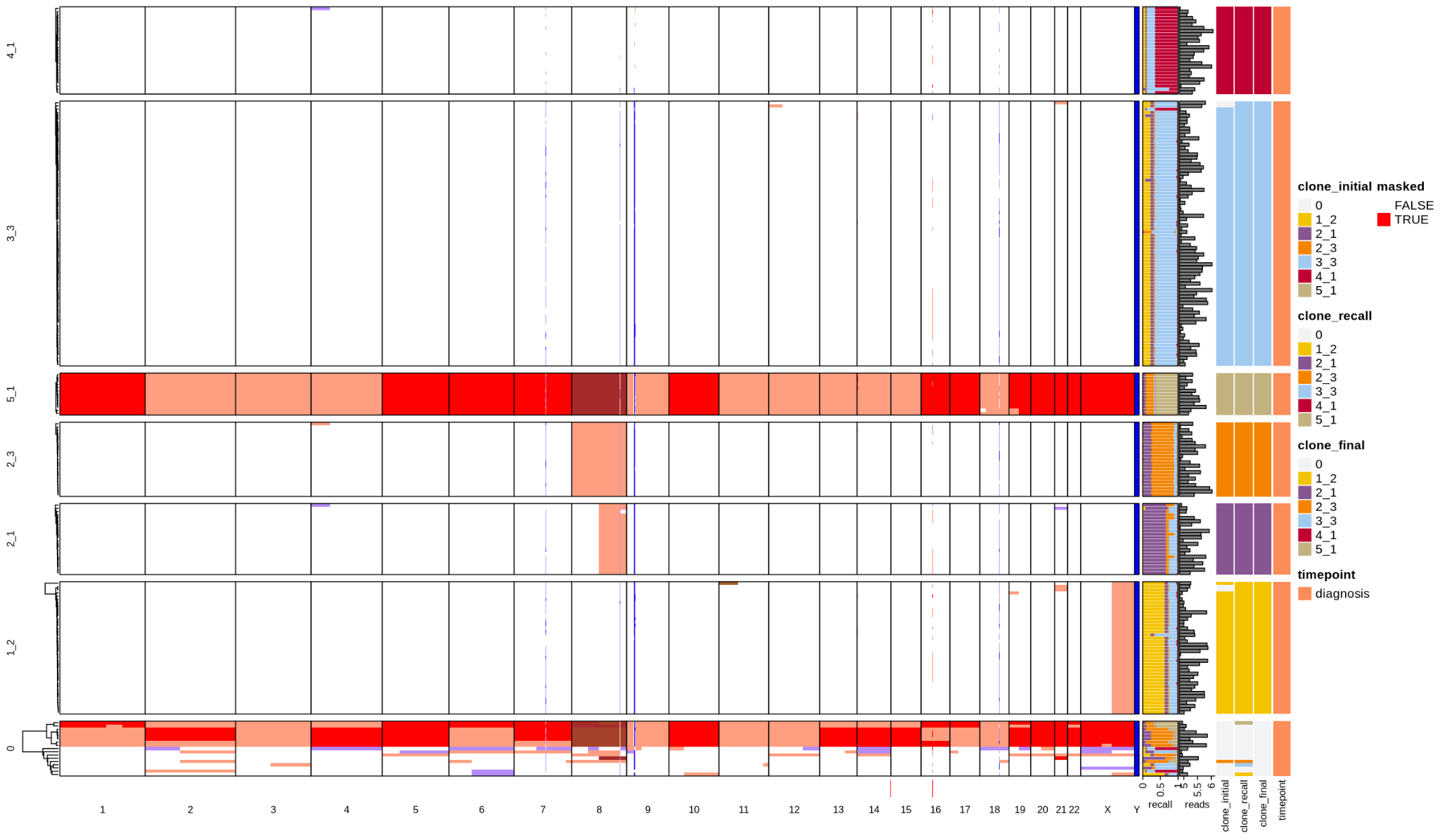

ALL8

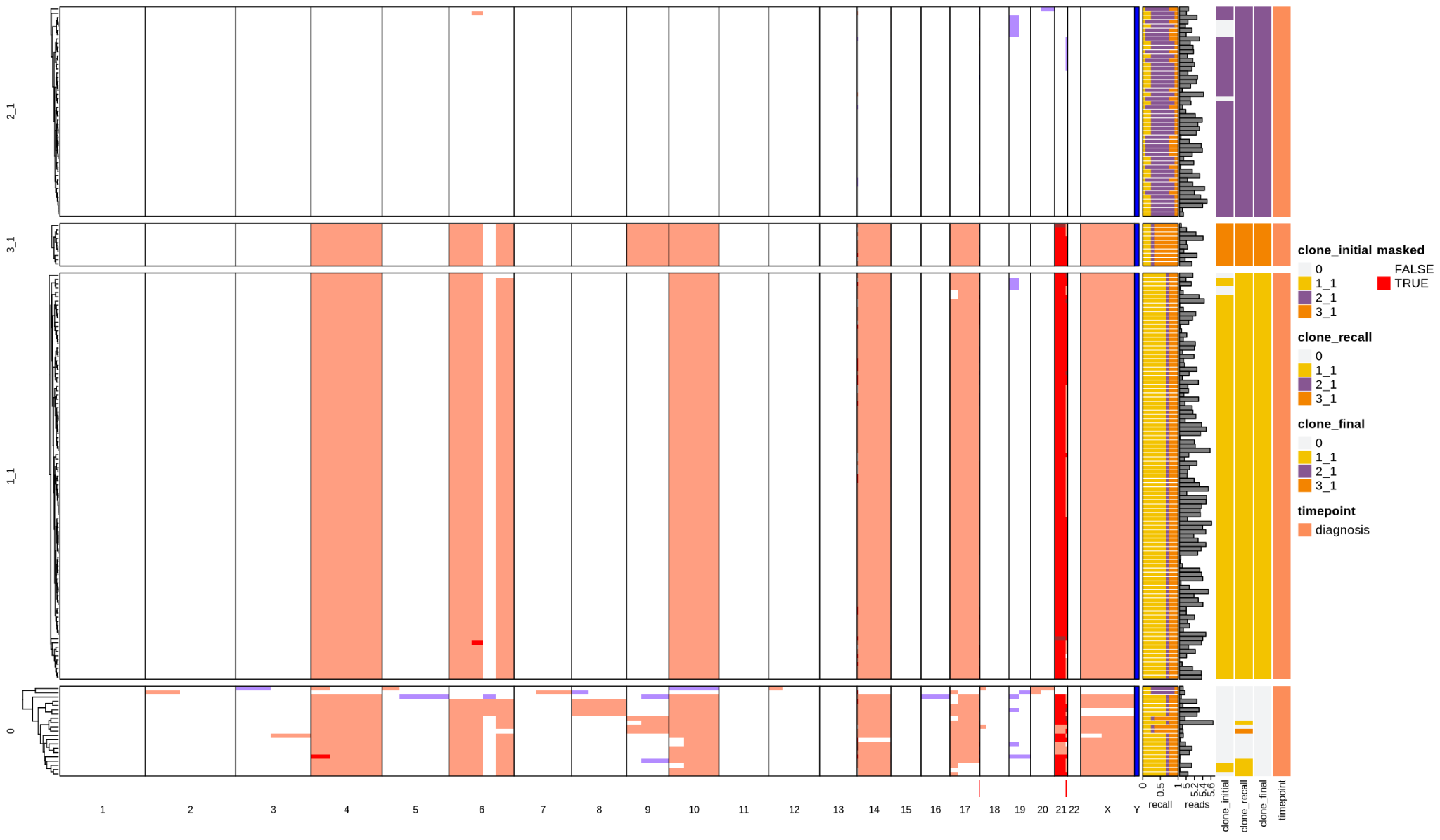

ALL35

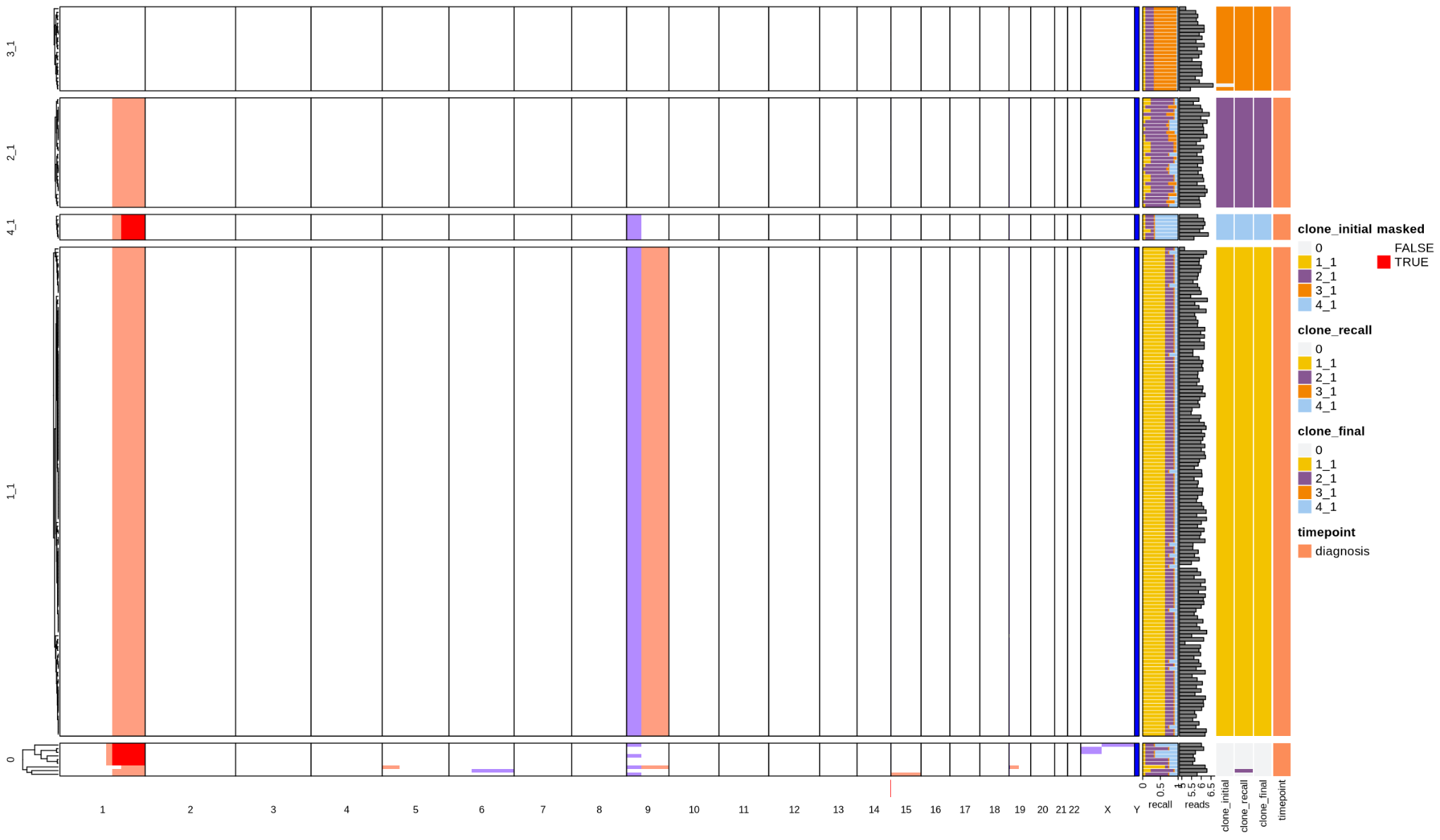

ALL40

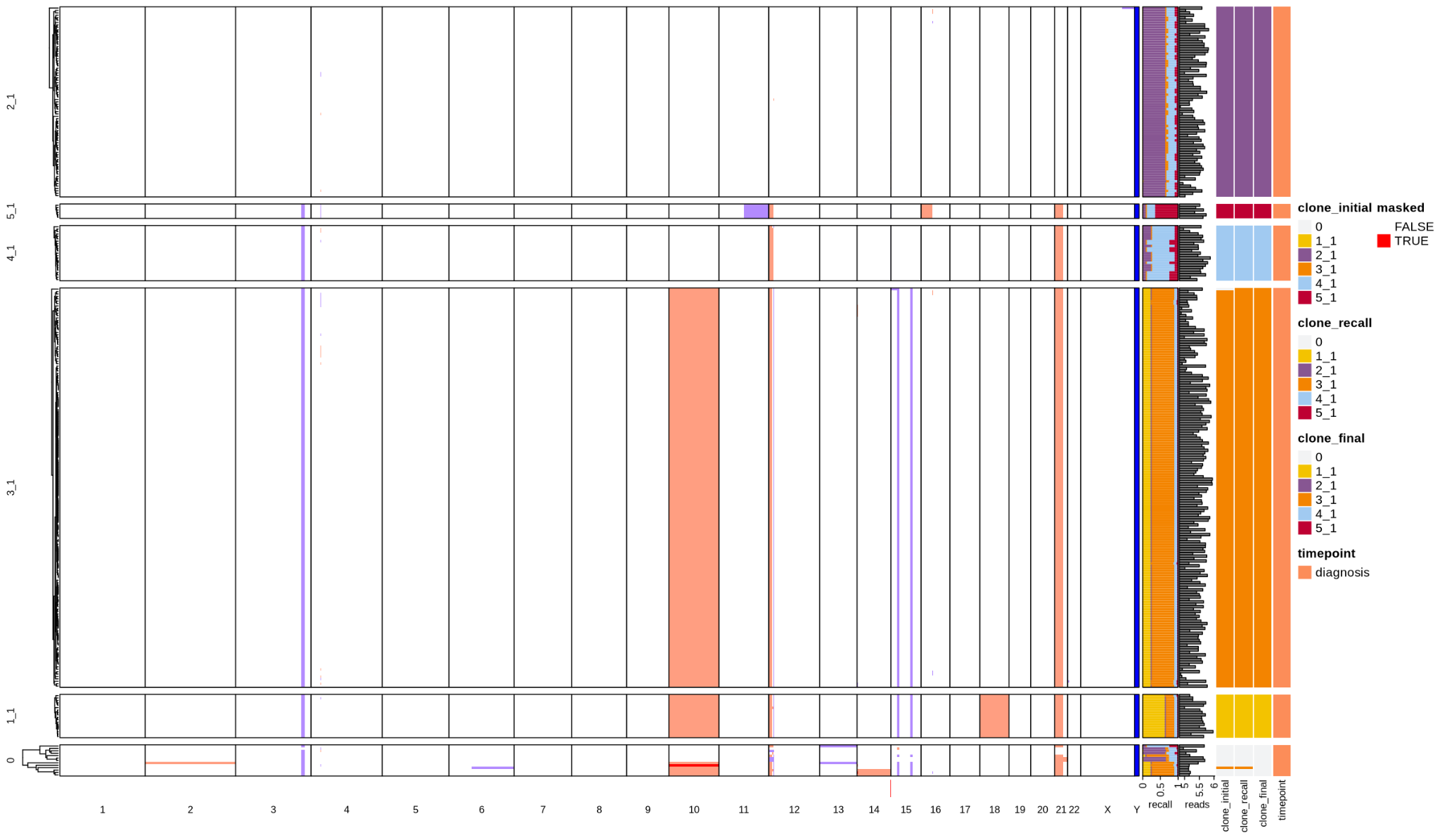

ALL42

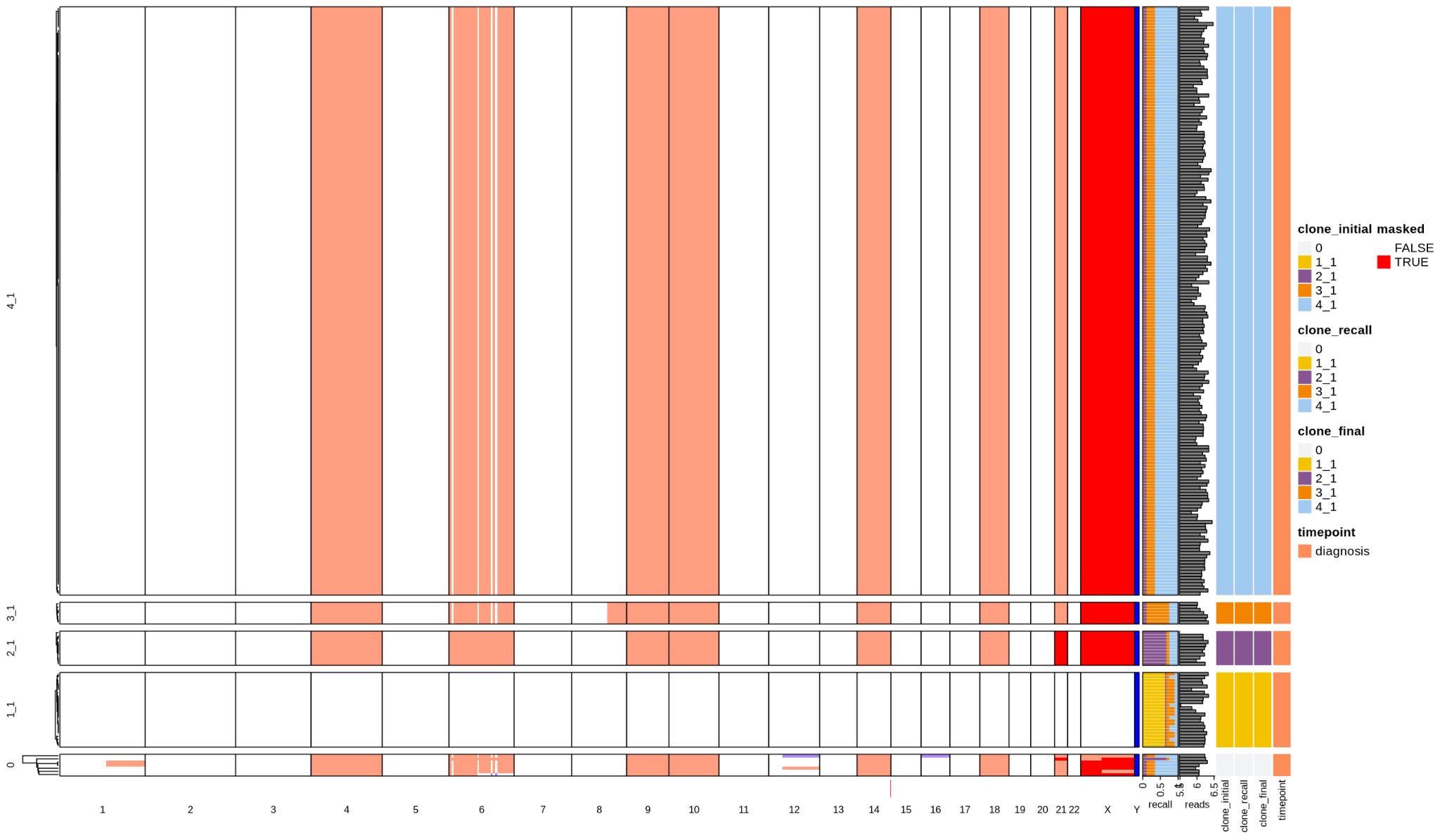

ALL47

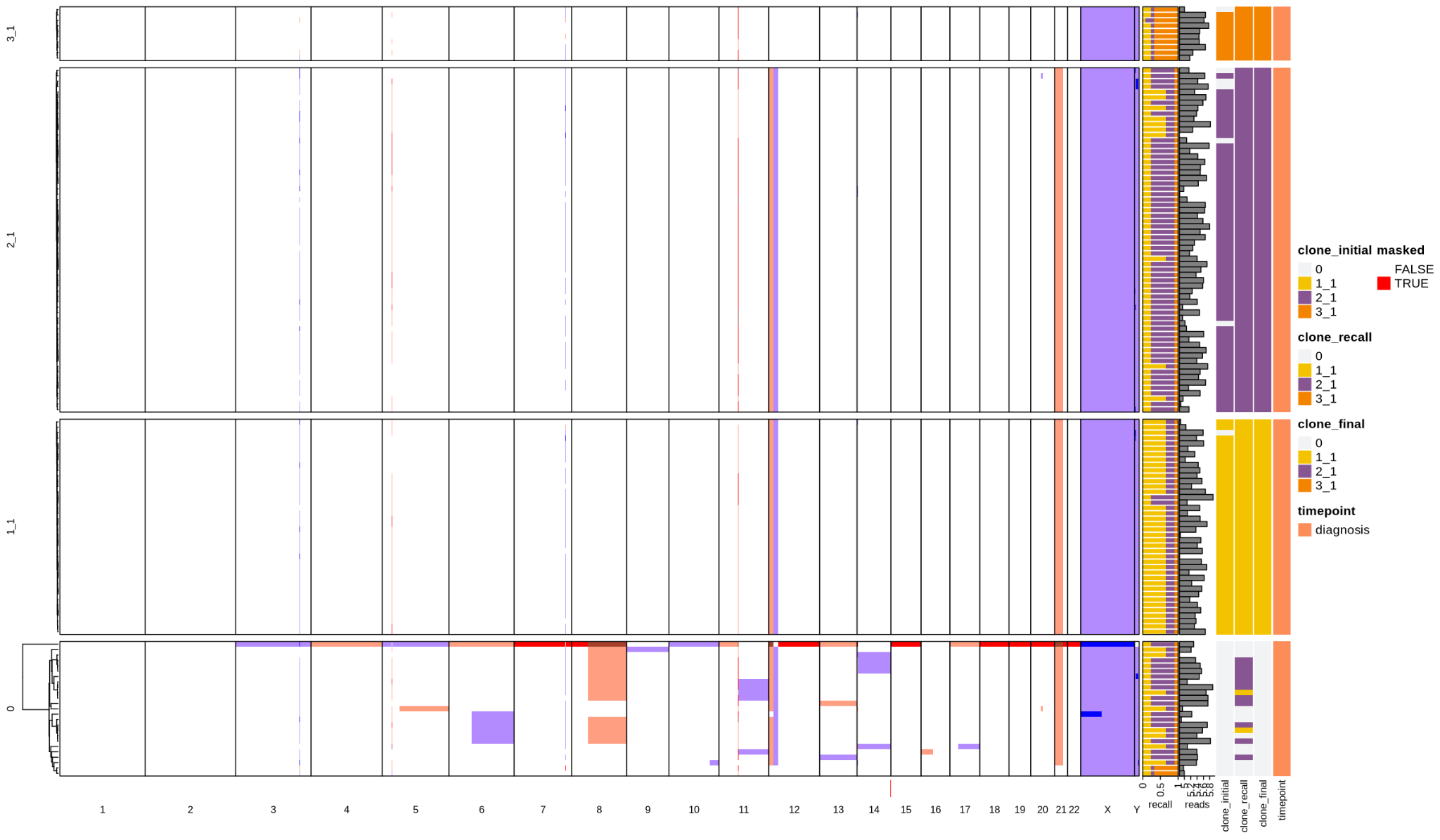

ALL64

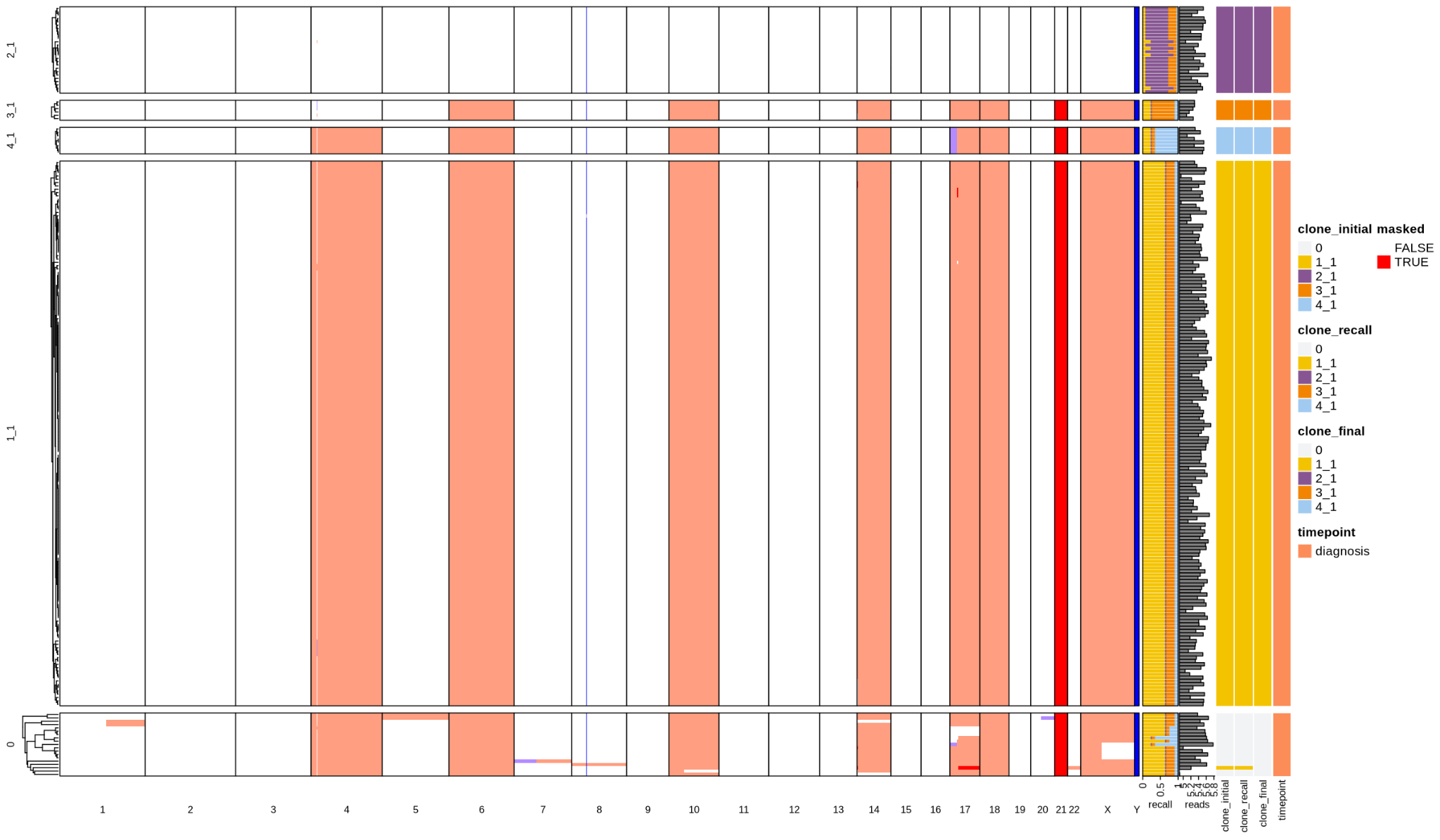

ALL66

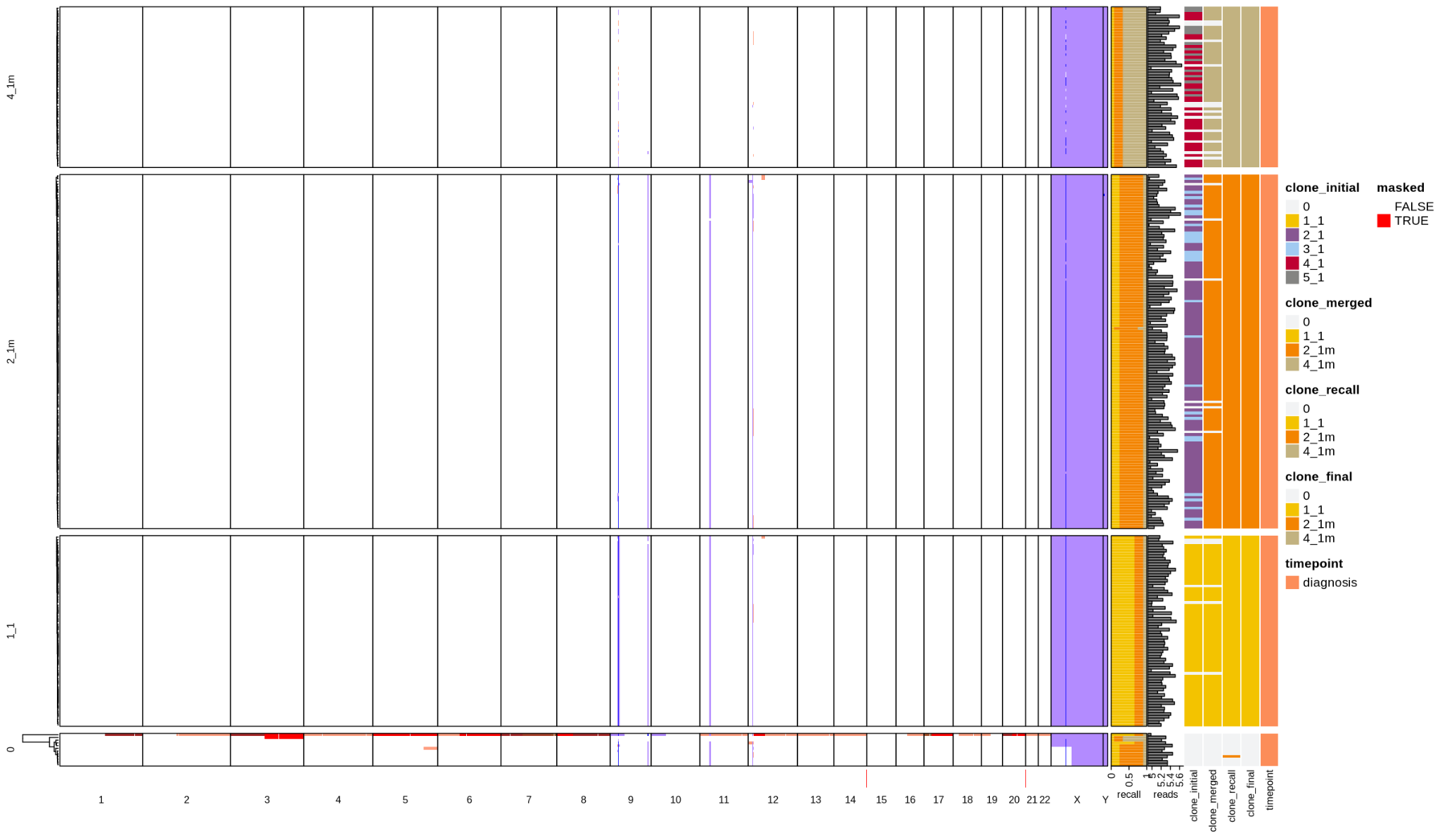

ALL67

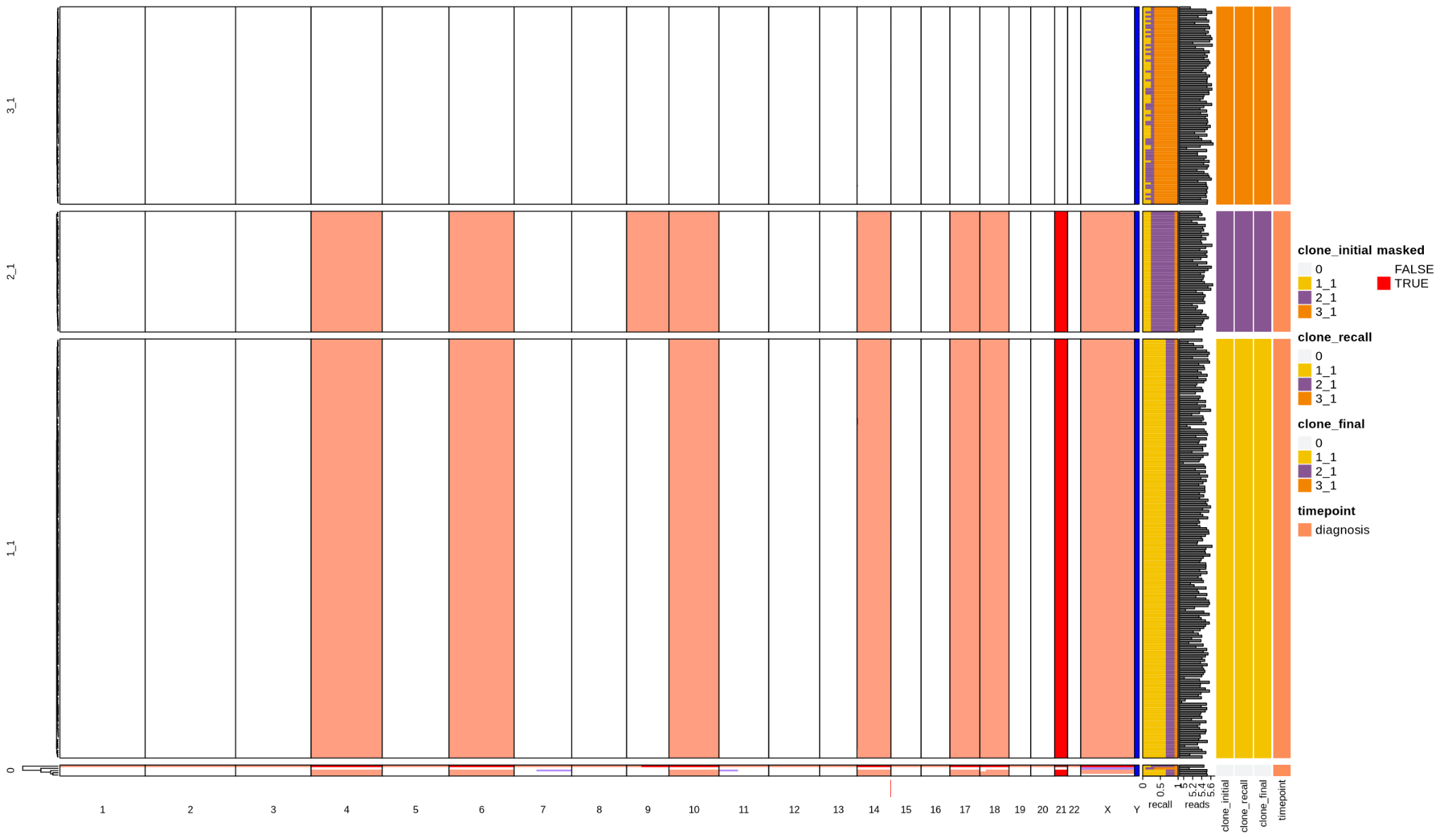

ALL68

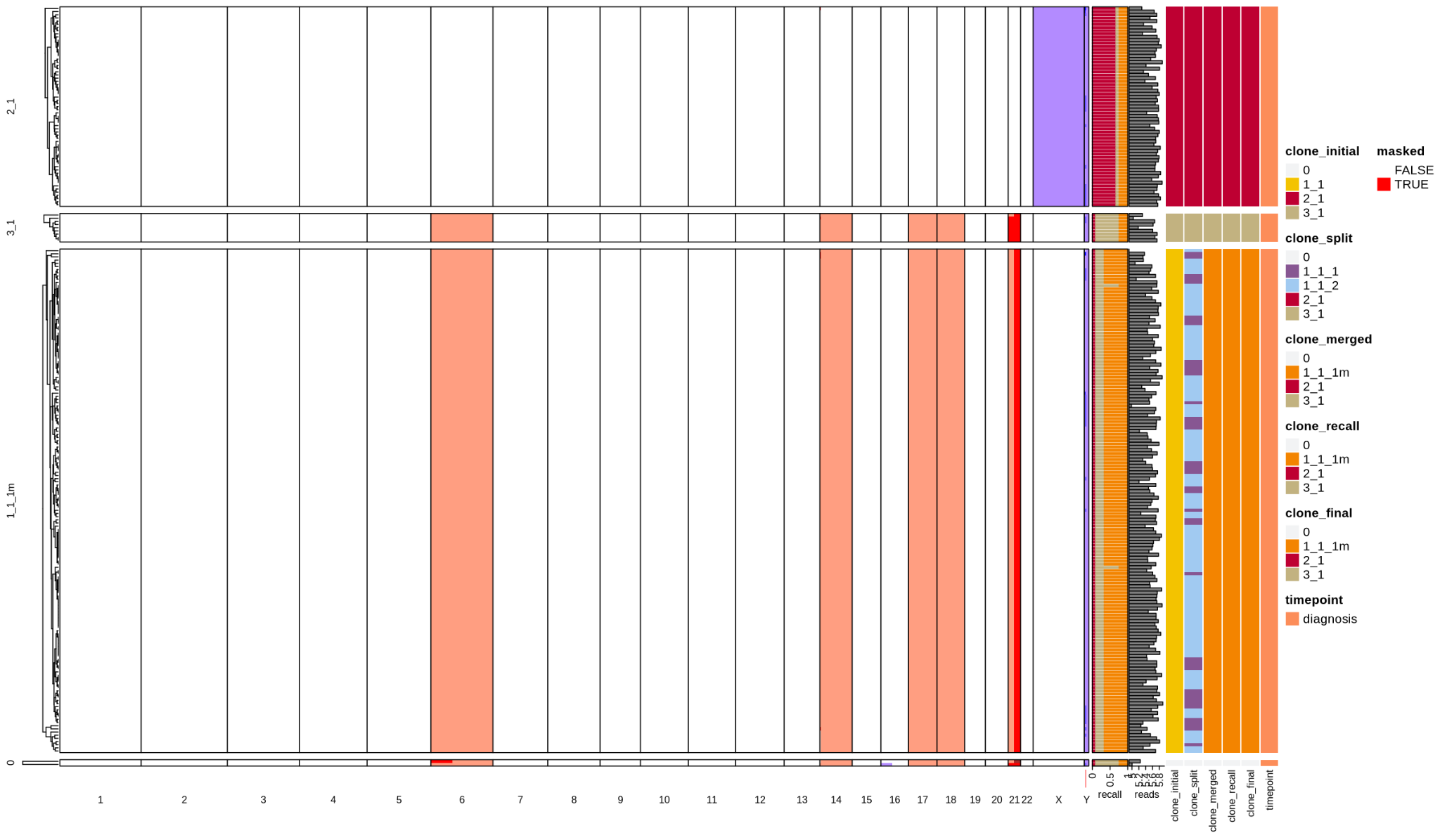

ALL71

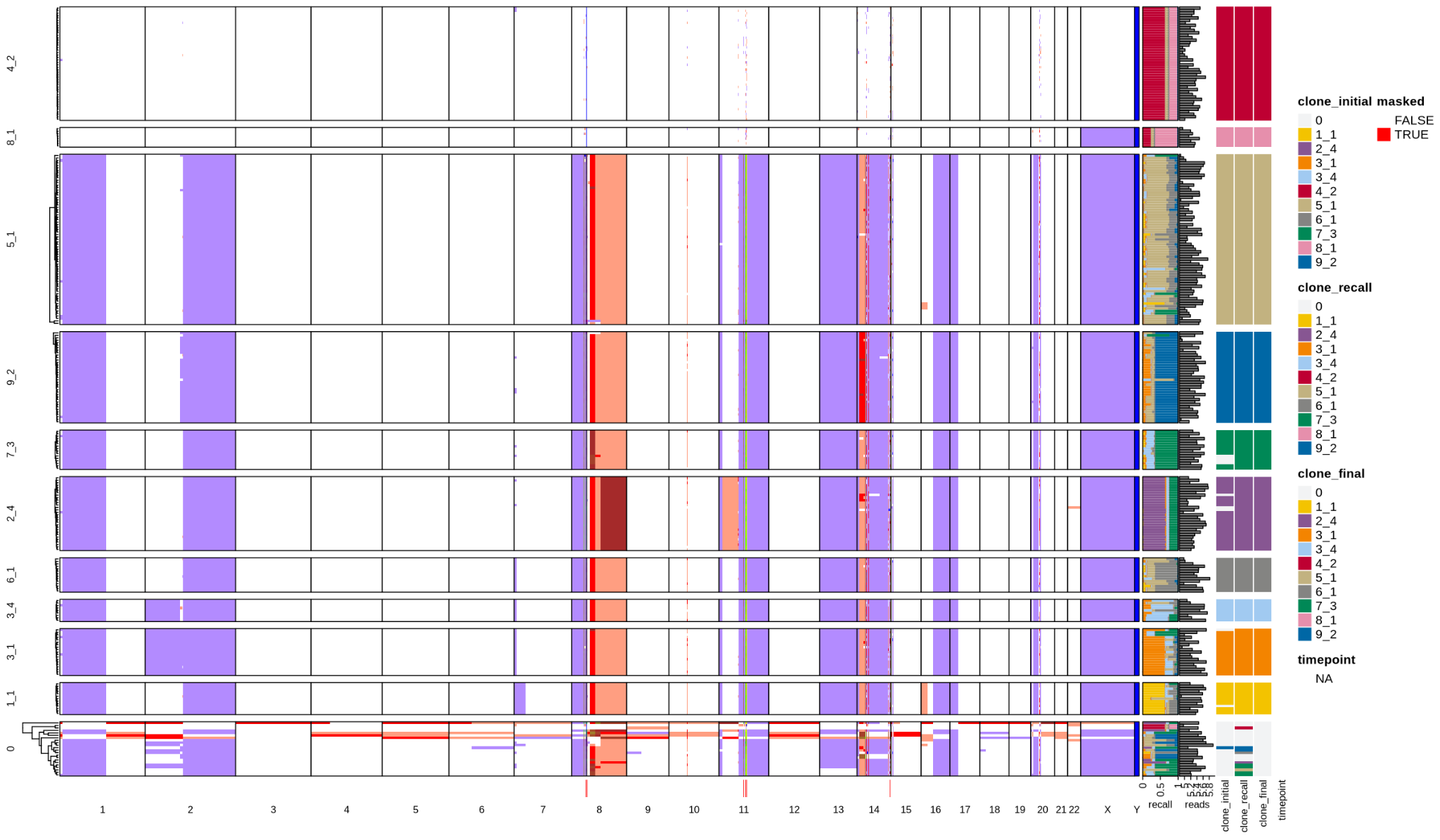

BC4

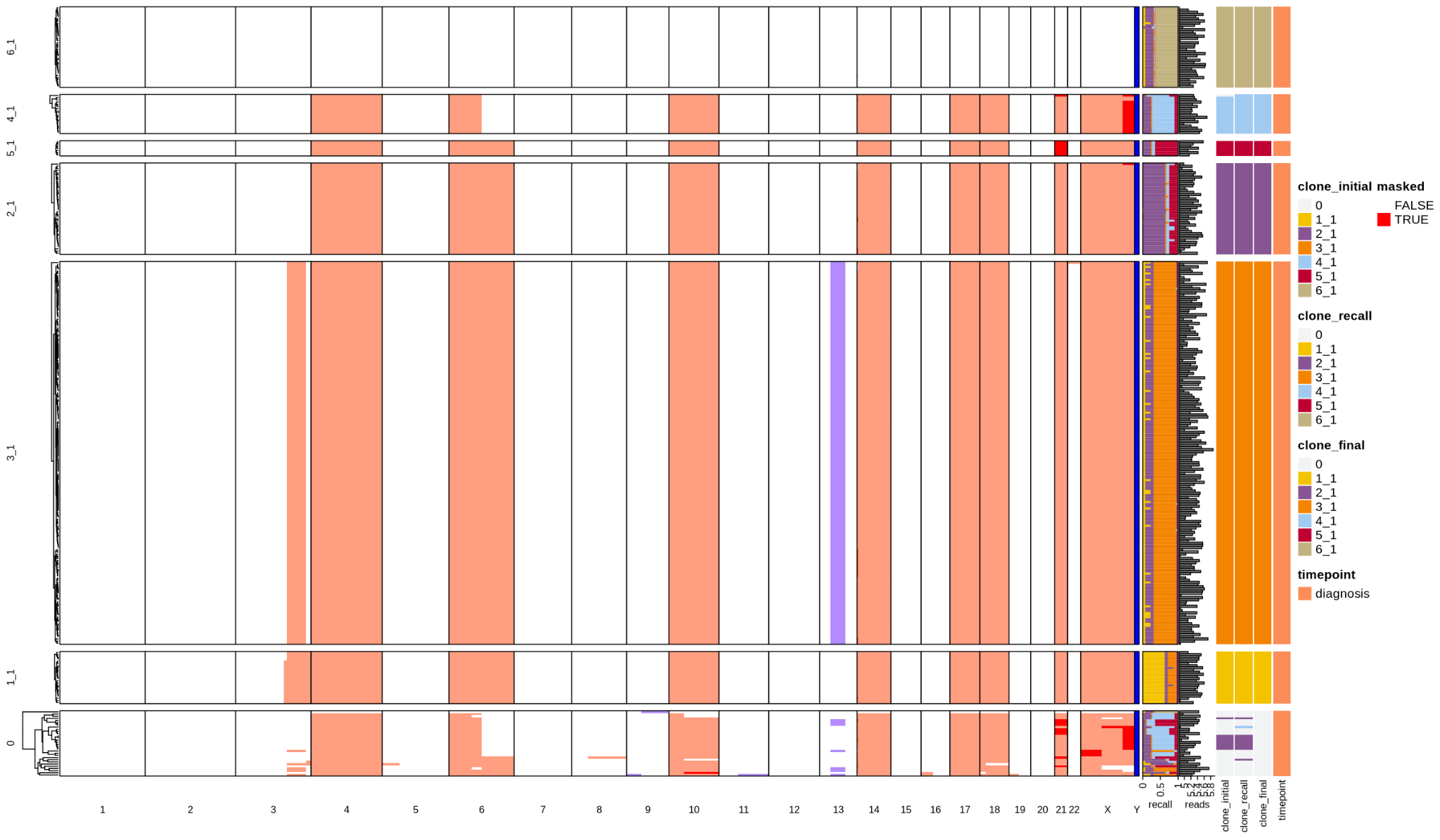

MEL5

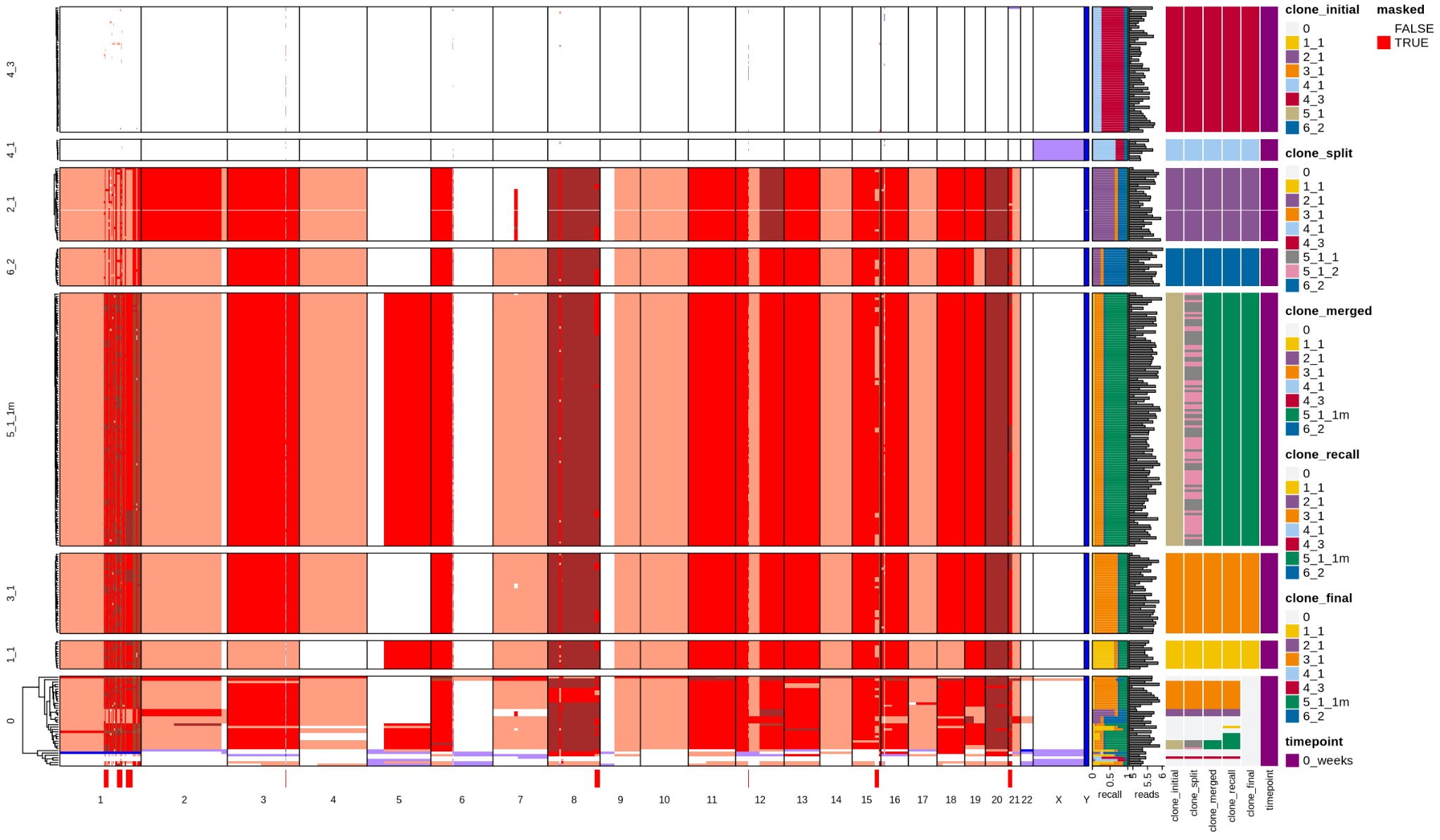

MEL6

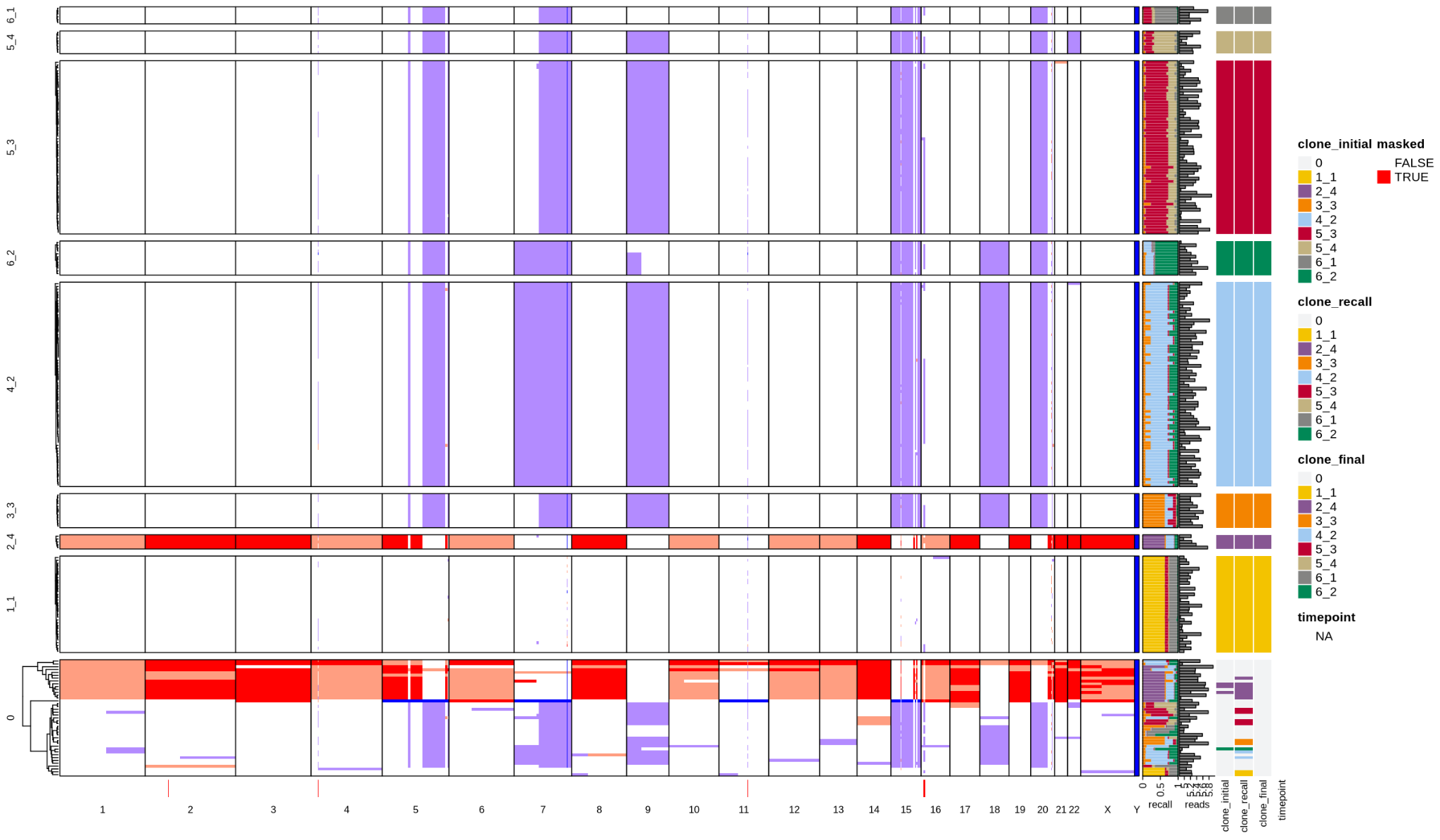

SRC20

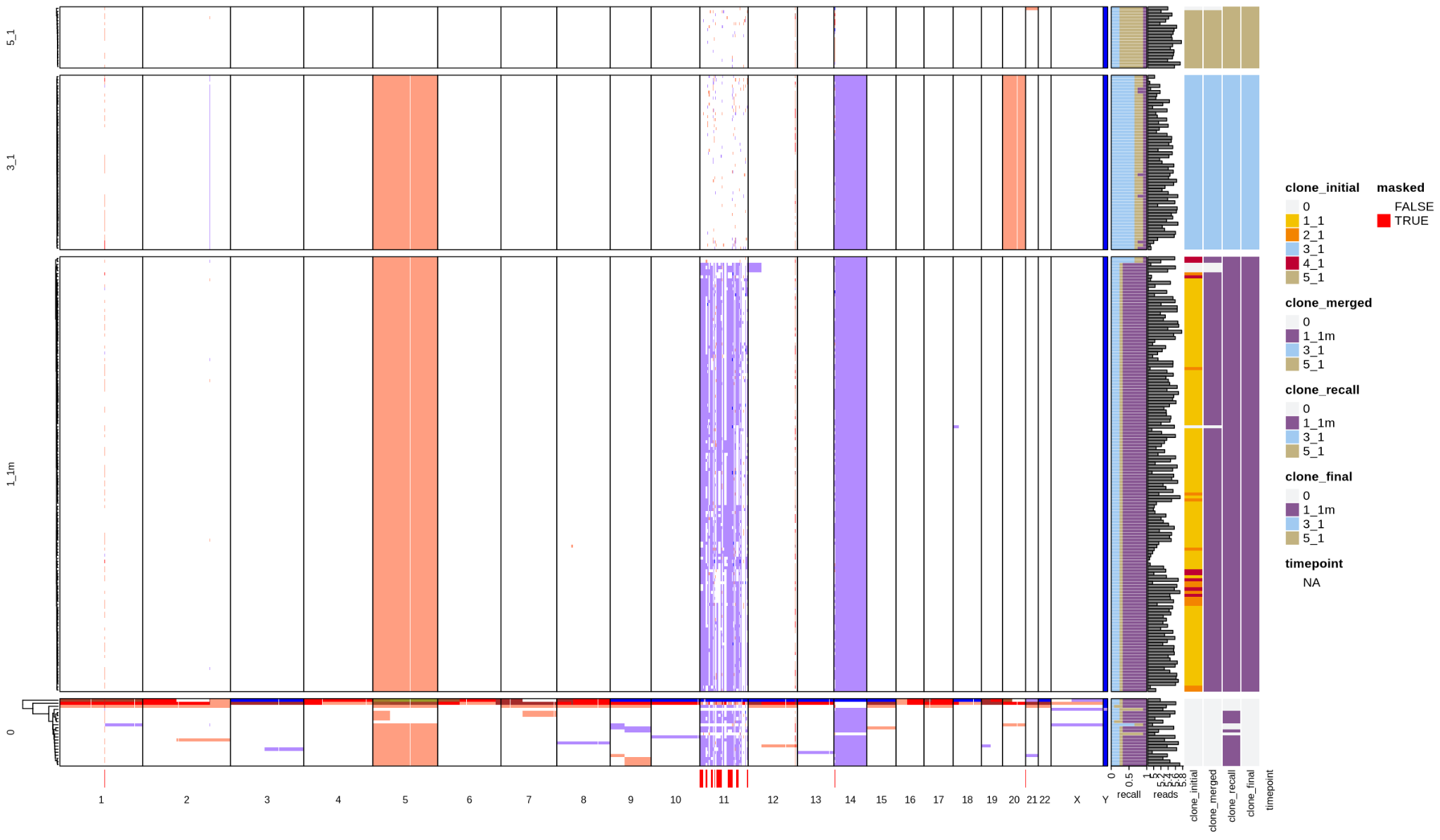

SRC21

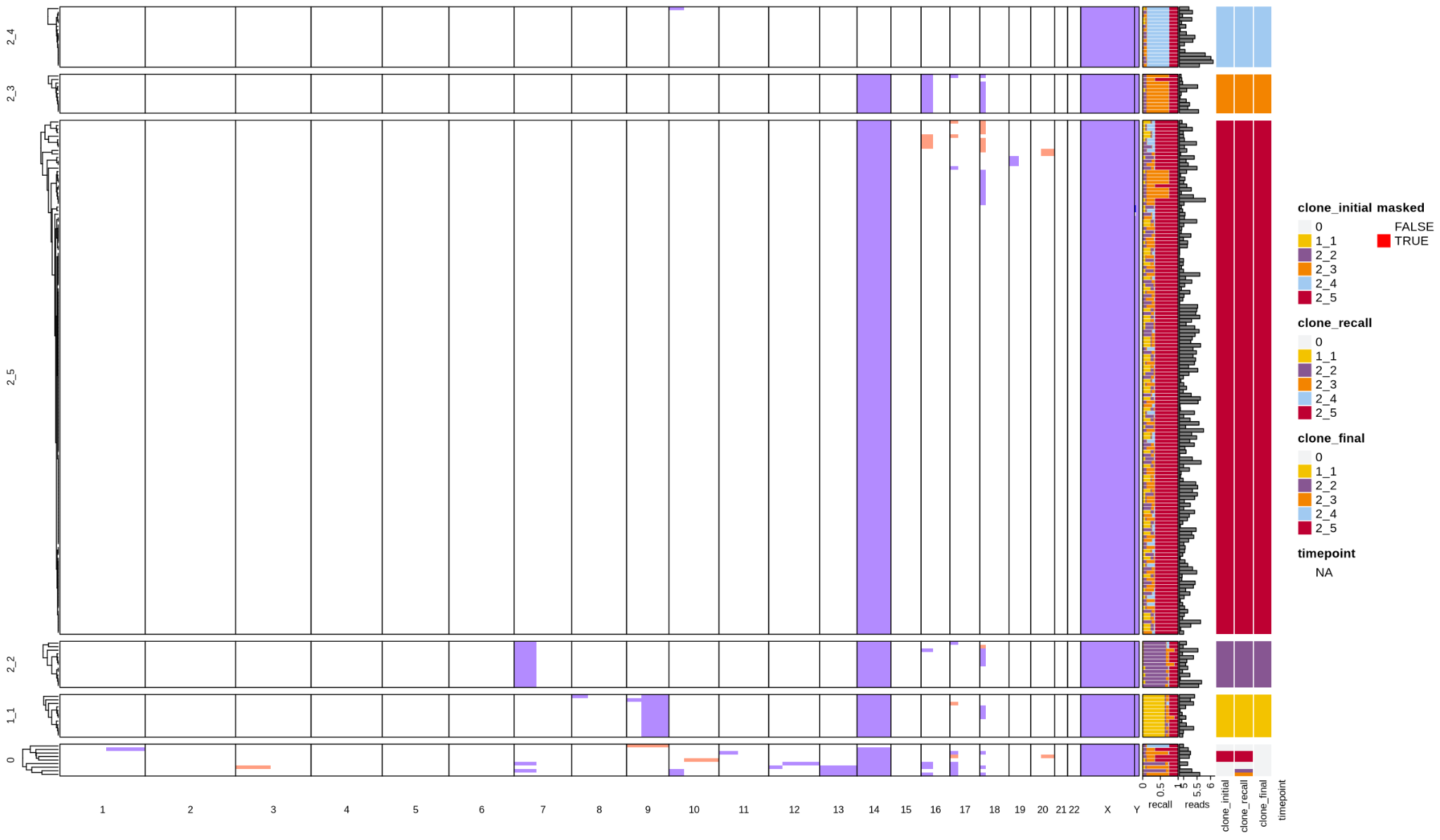

SRC23

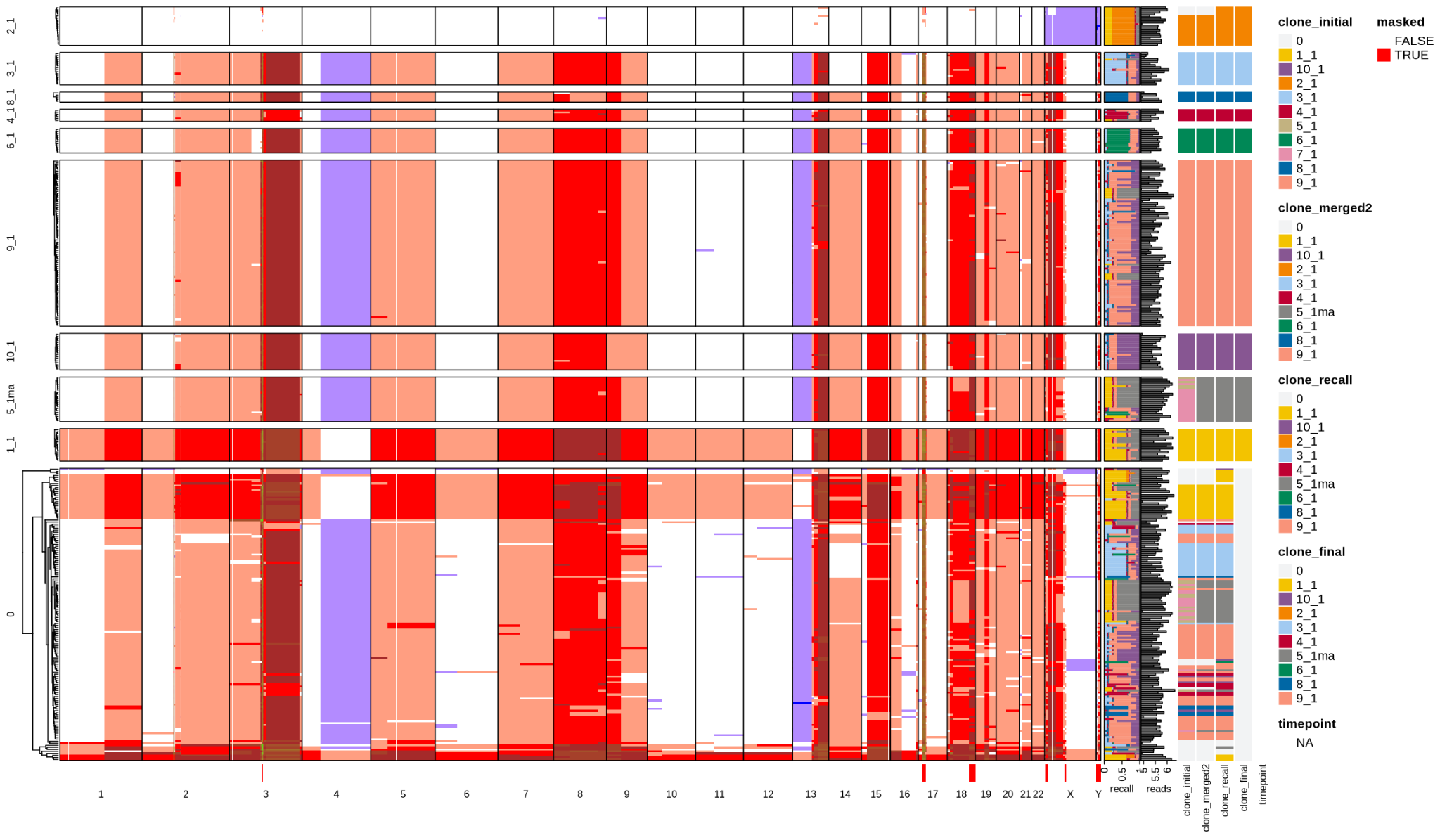

SRC27

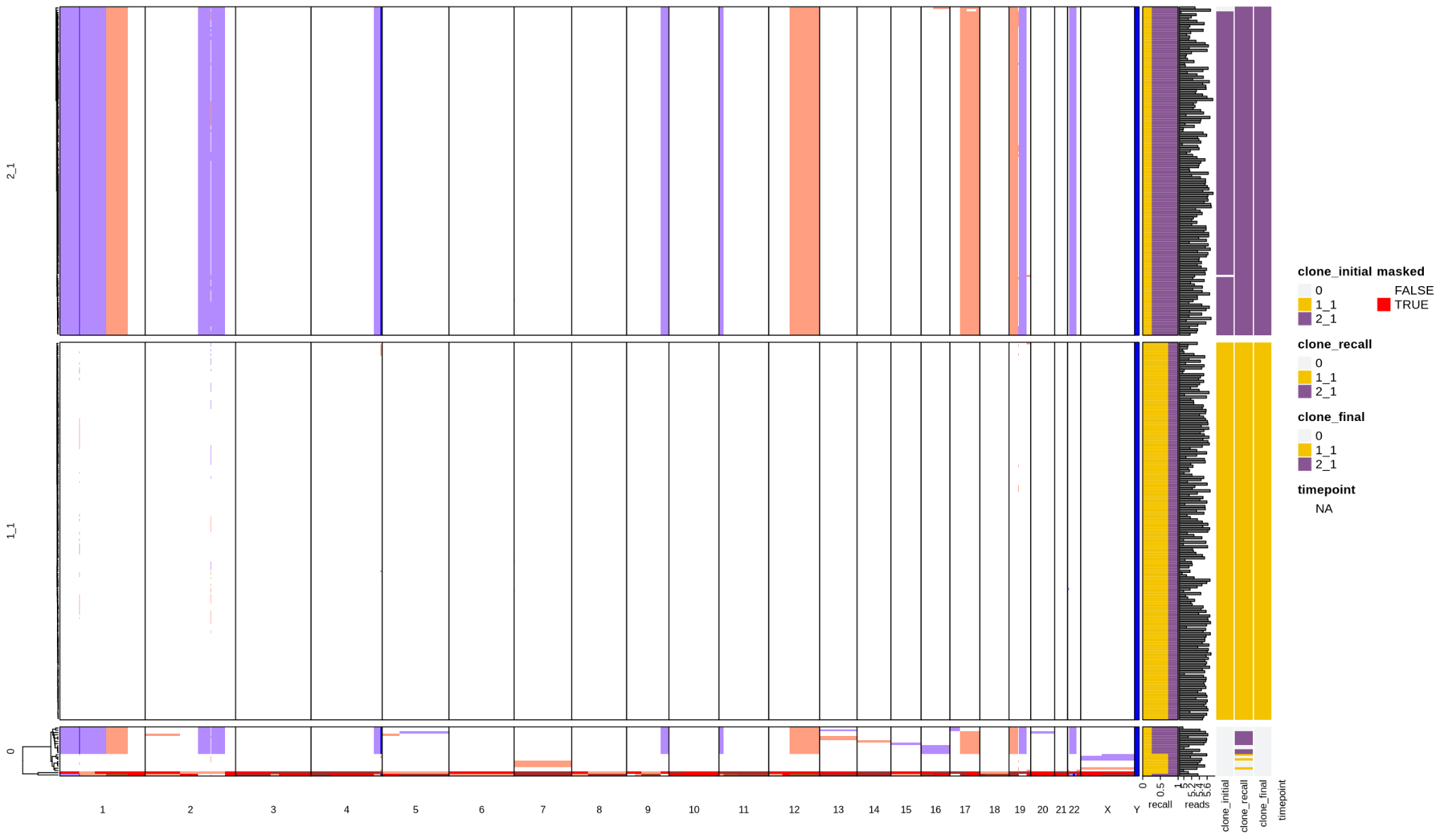

SRC35

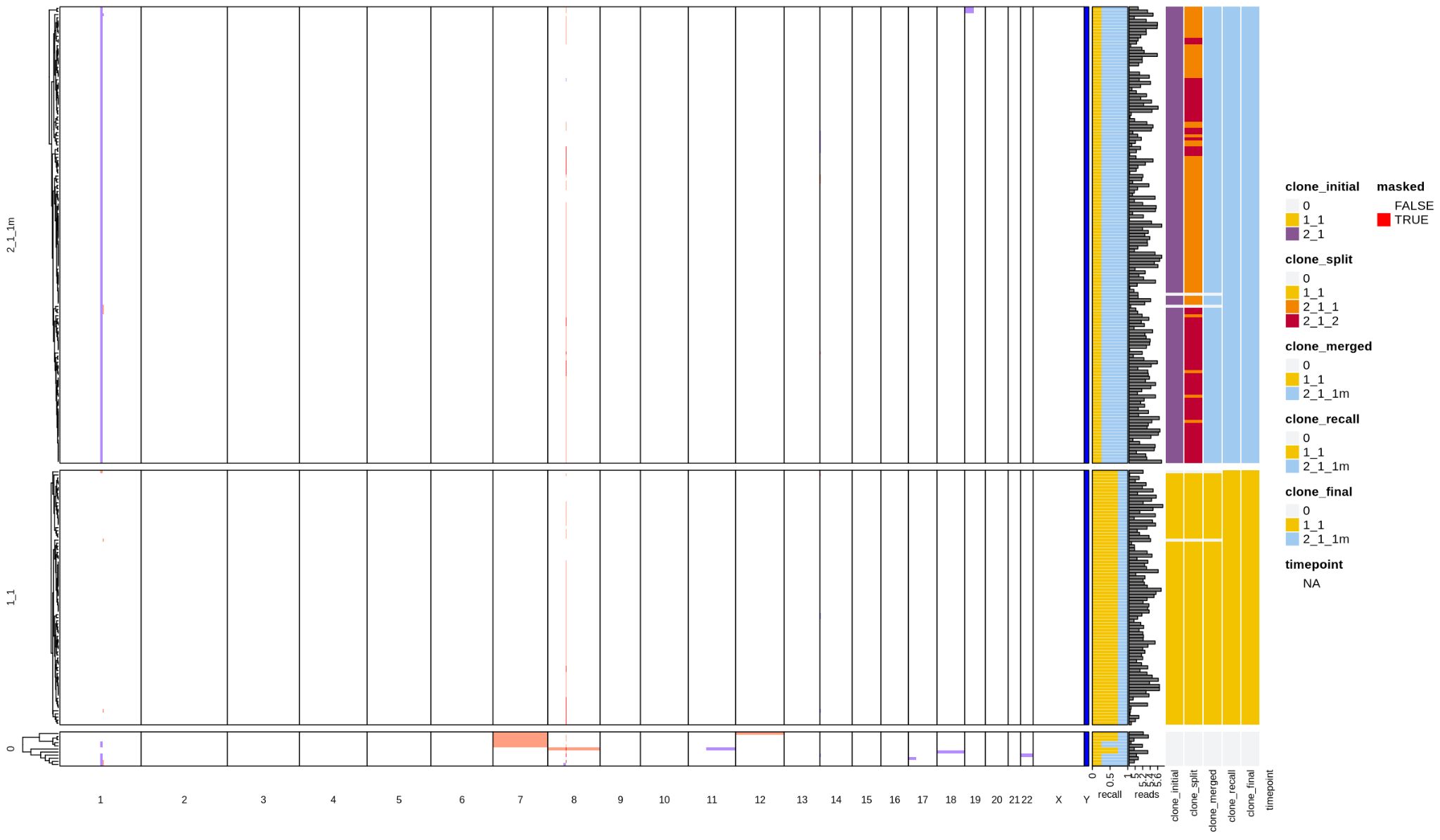

AML2

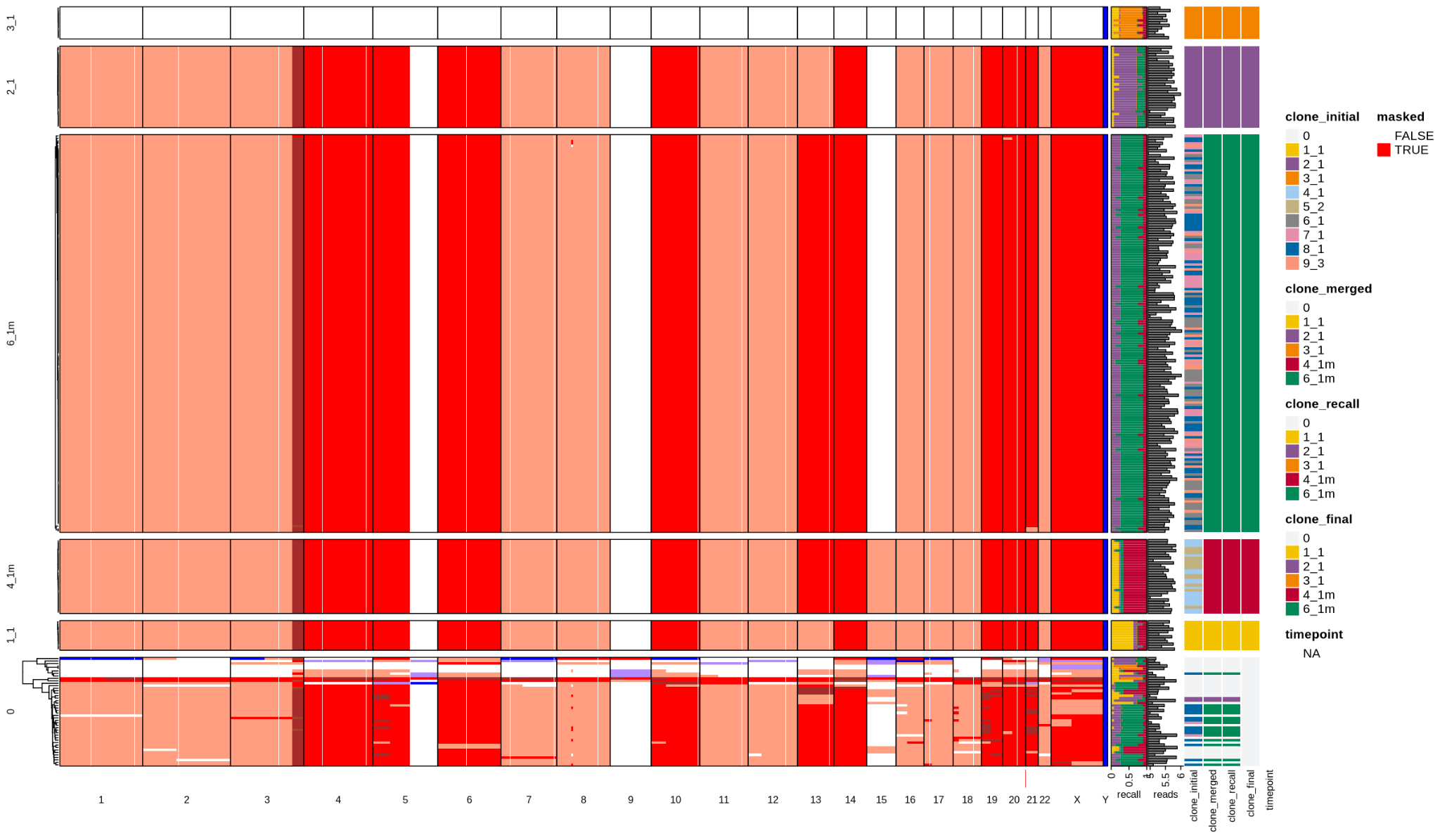

AML4

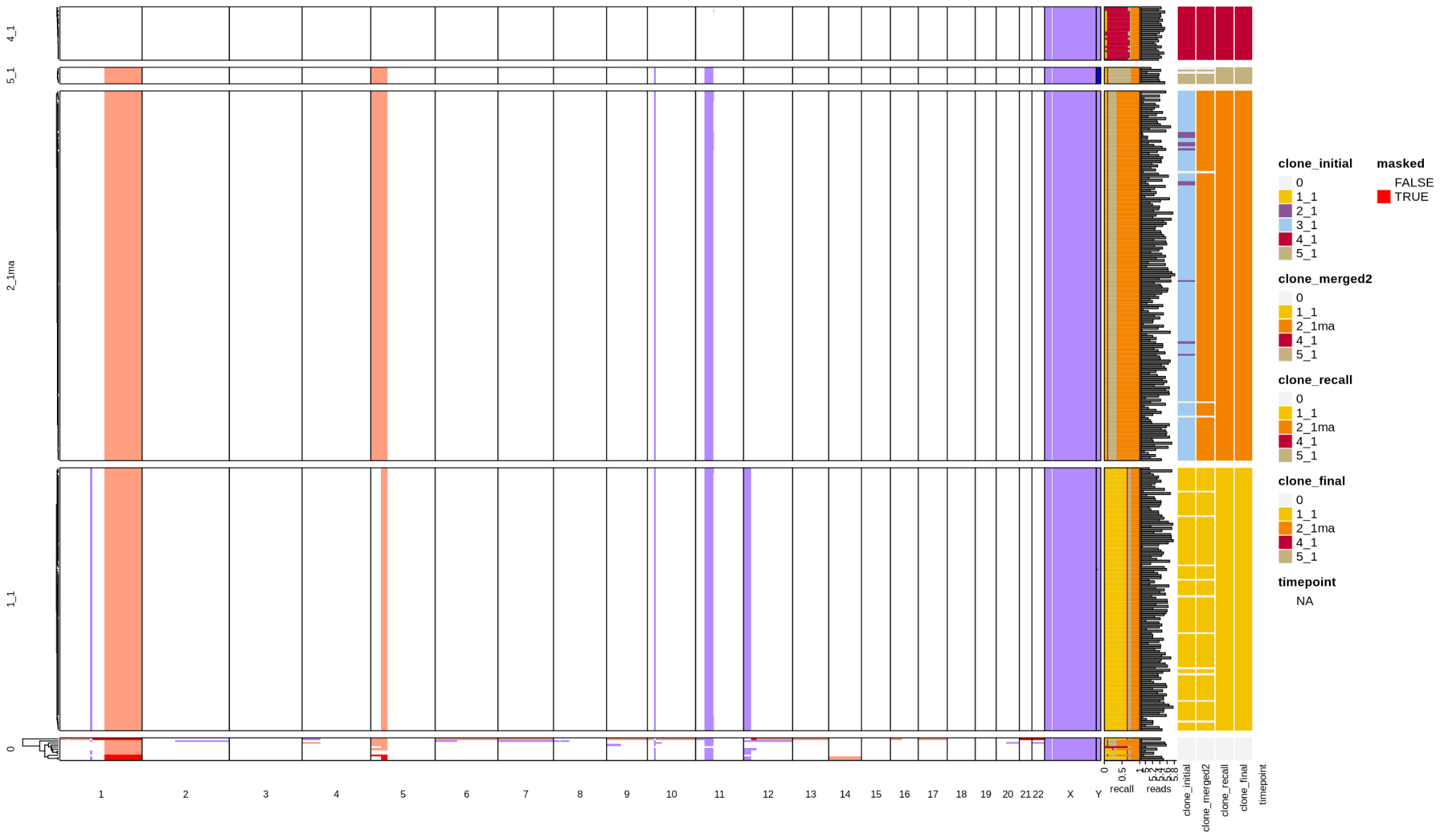

AML5

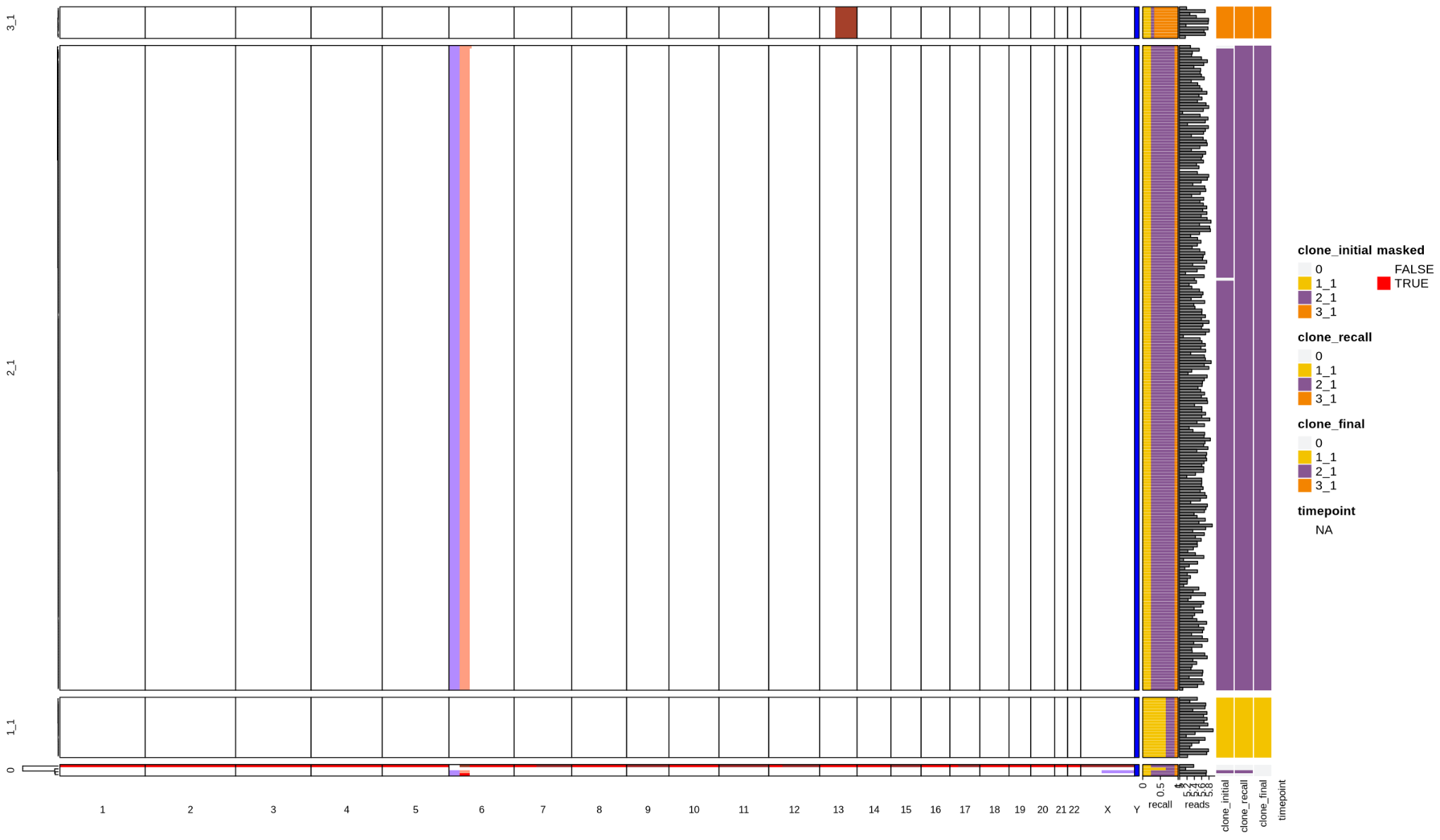

AML6
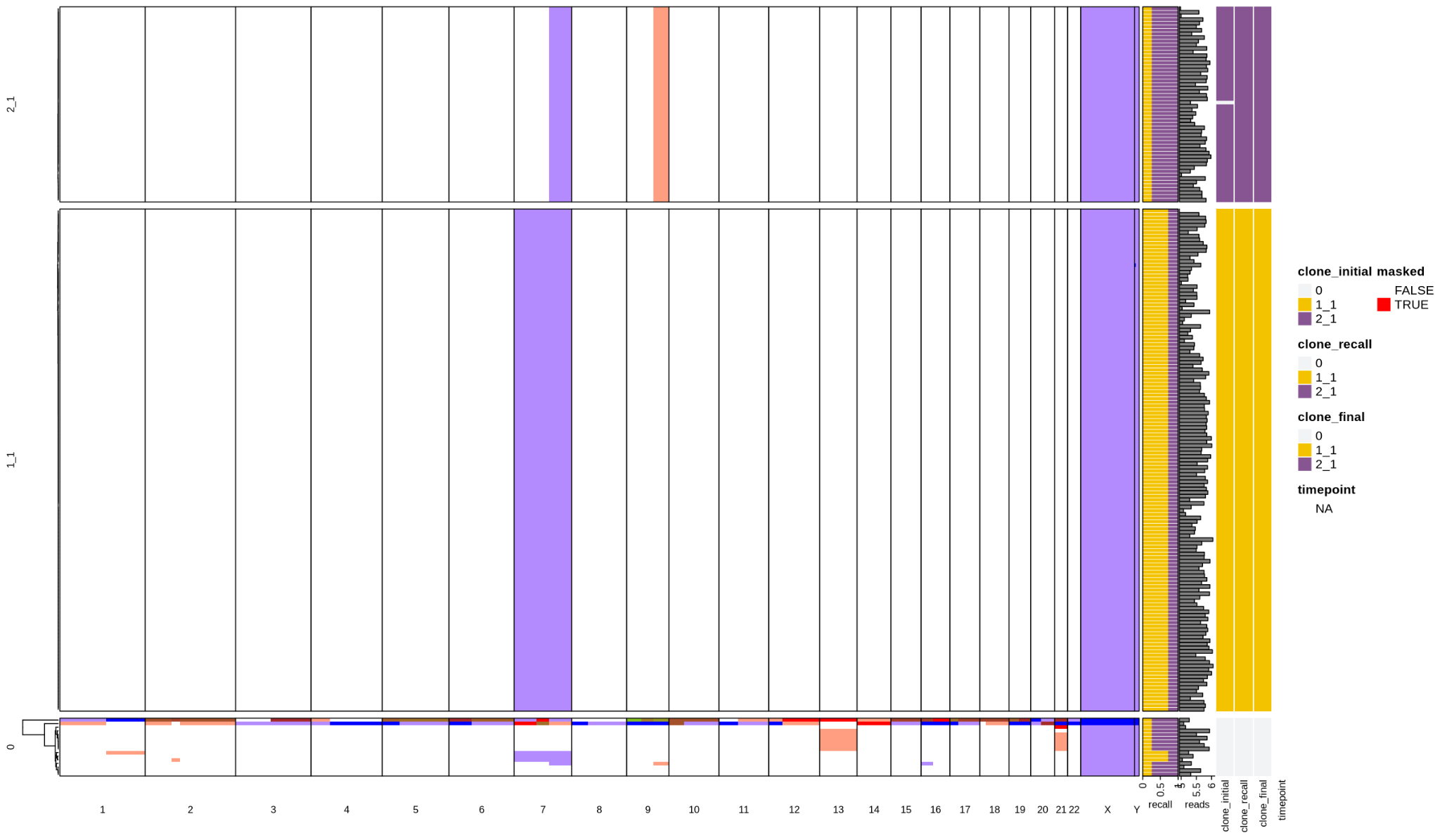

AML9

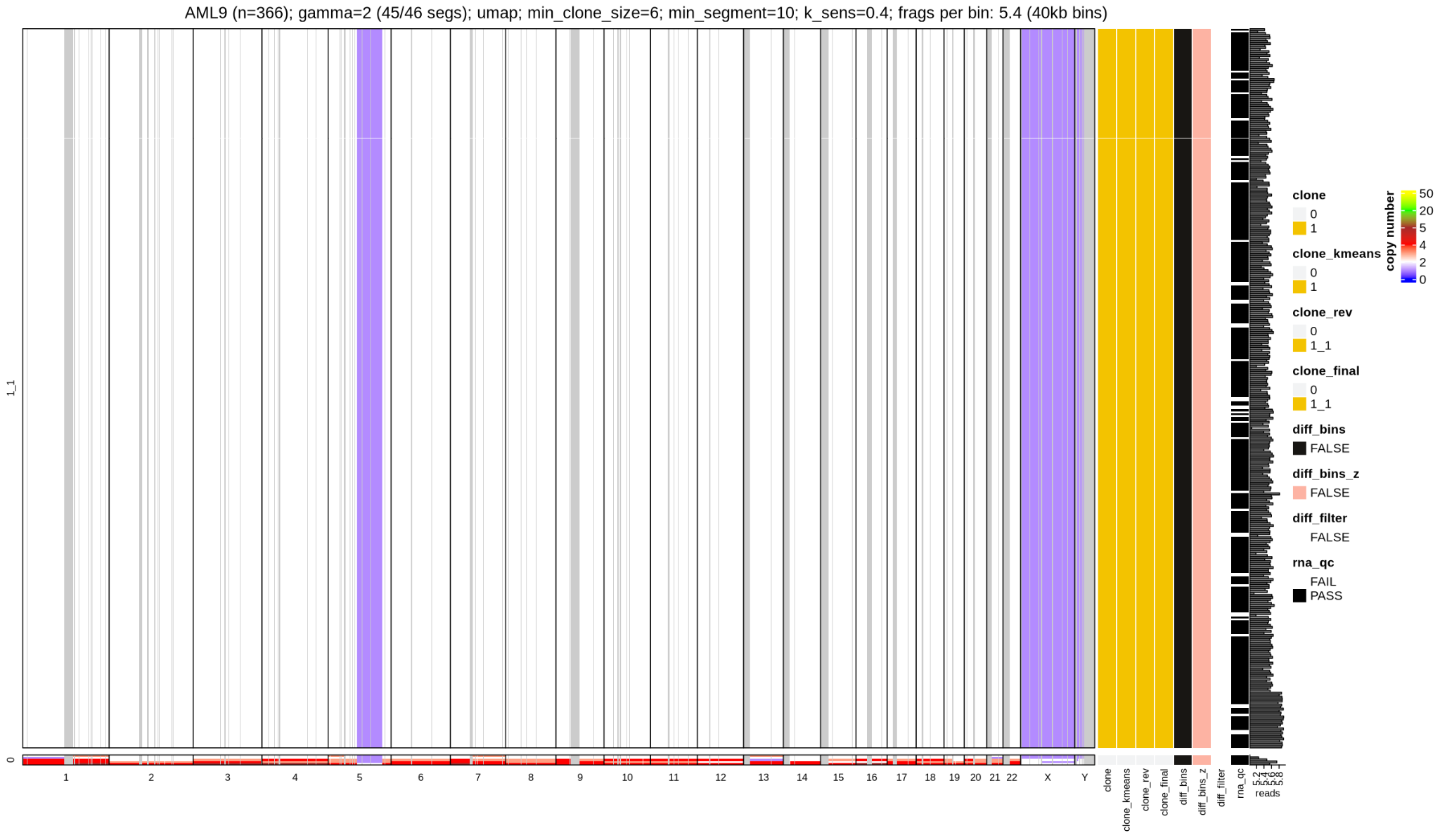

OC1

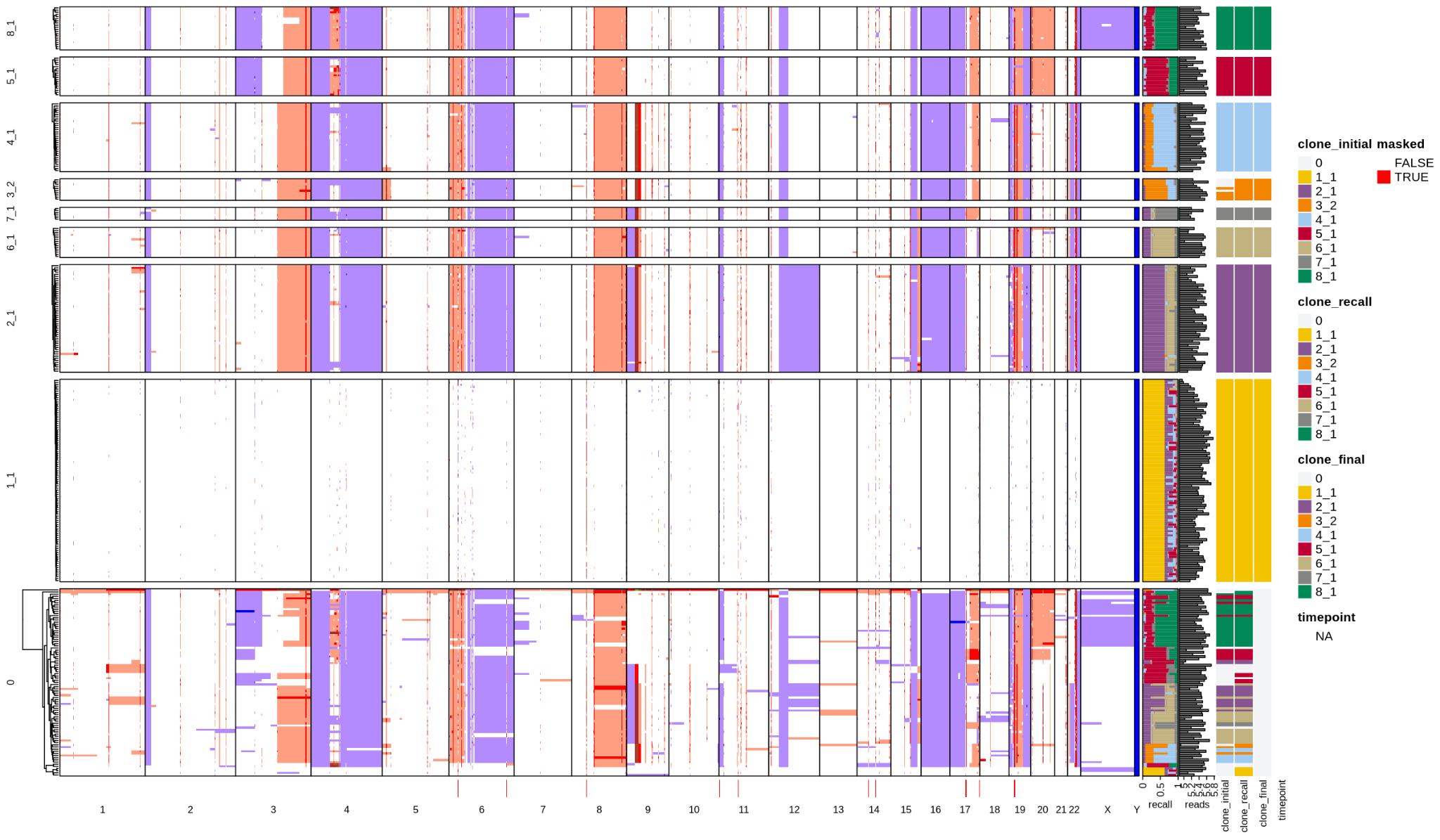

OC2

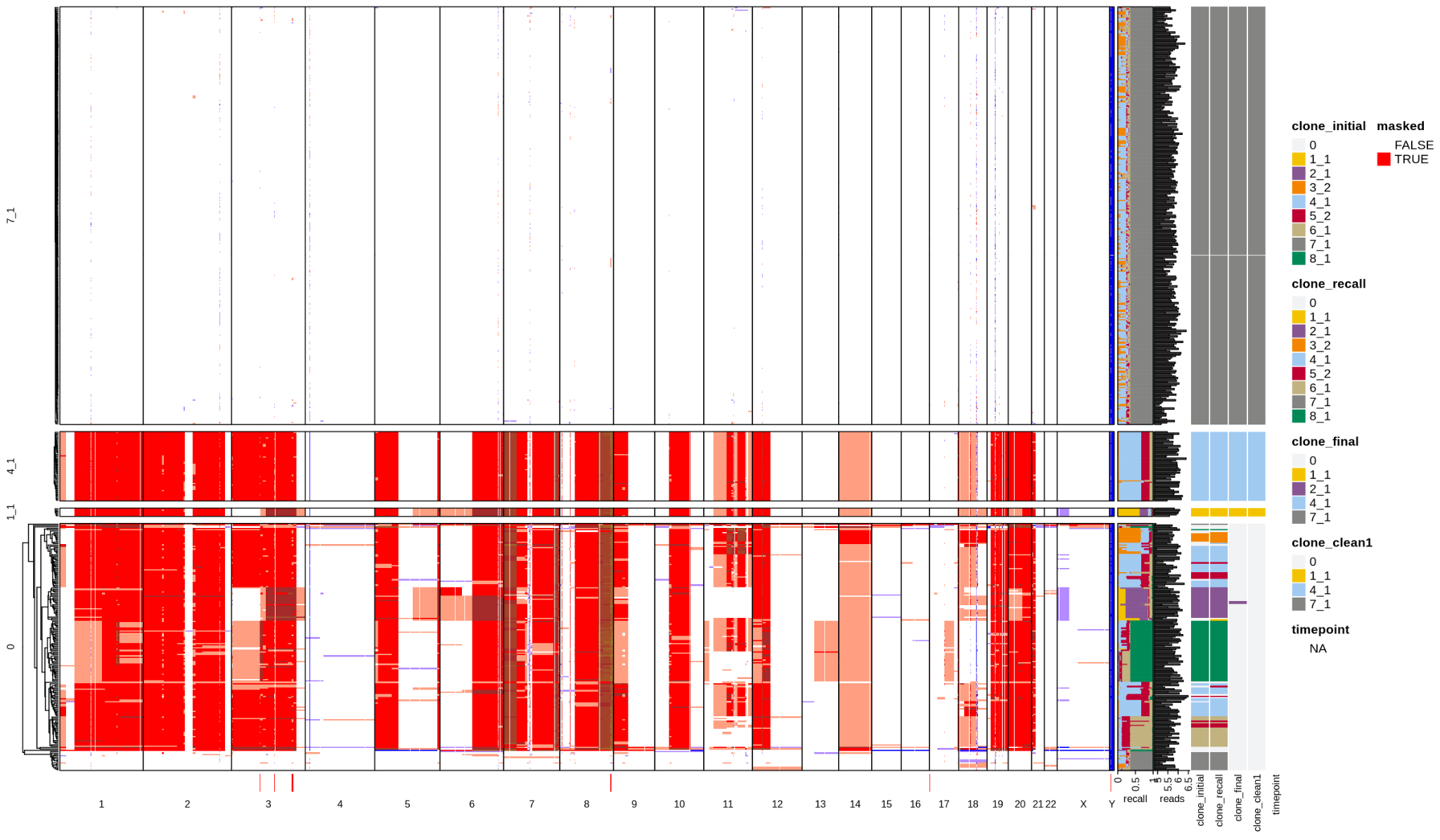

OC4

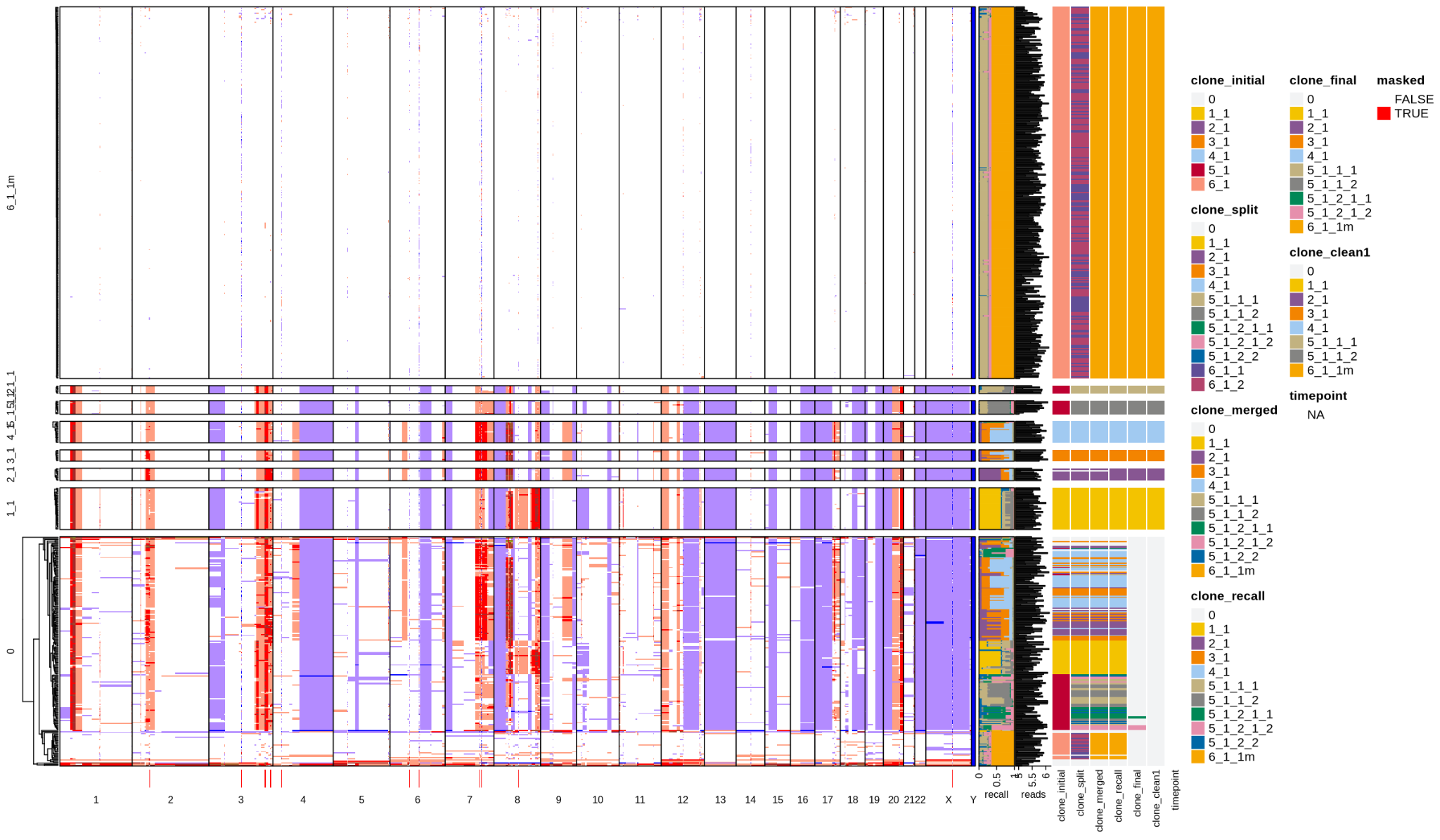

OC5

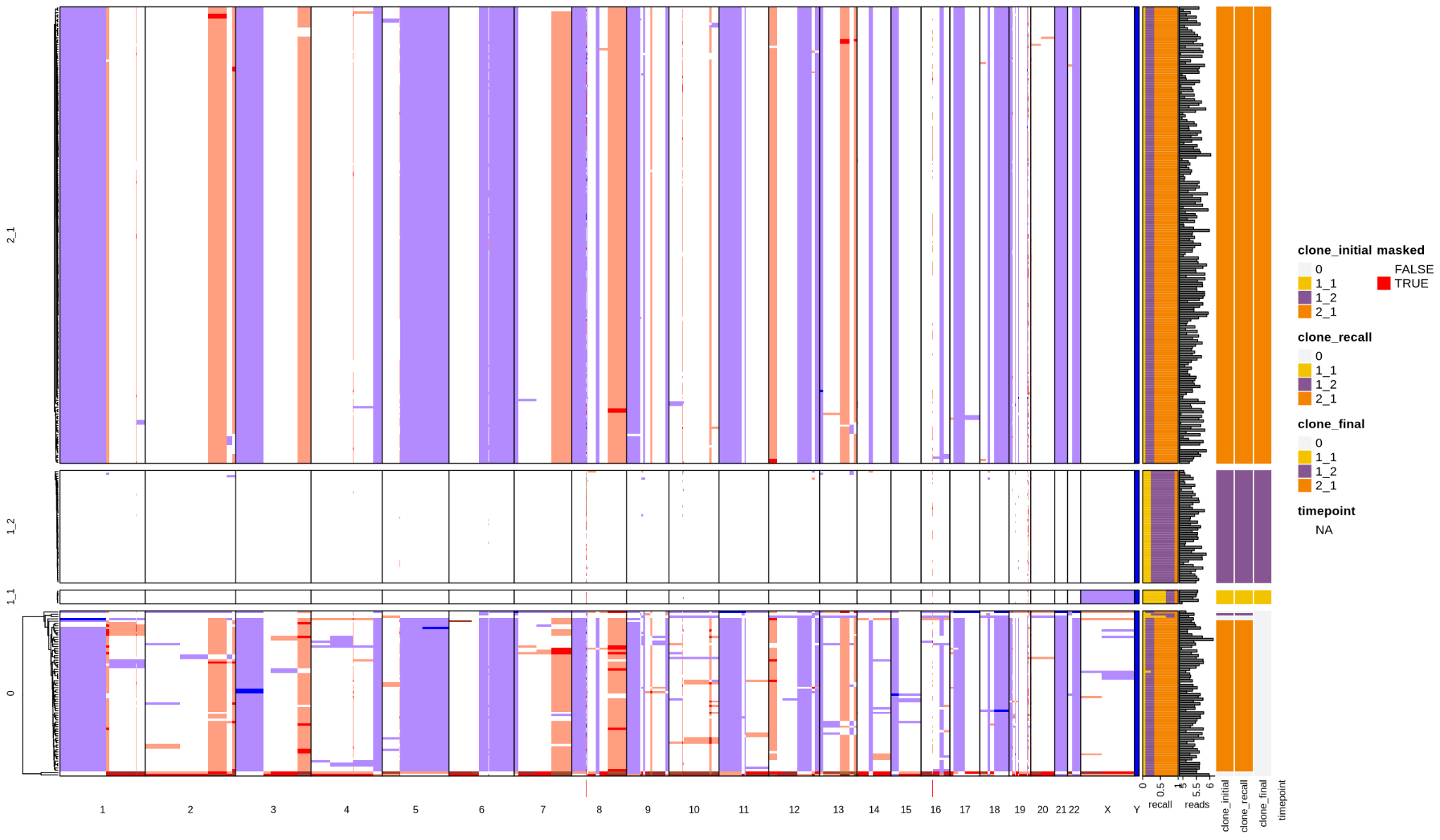

OC9

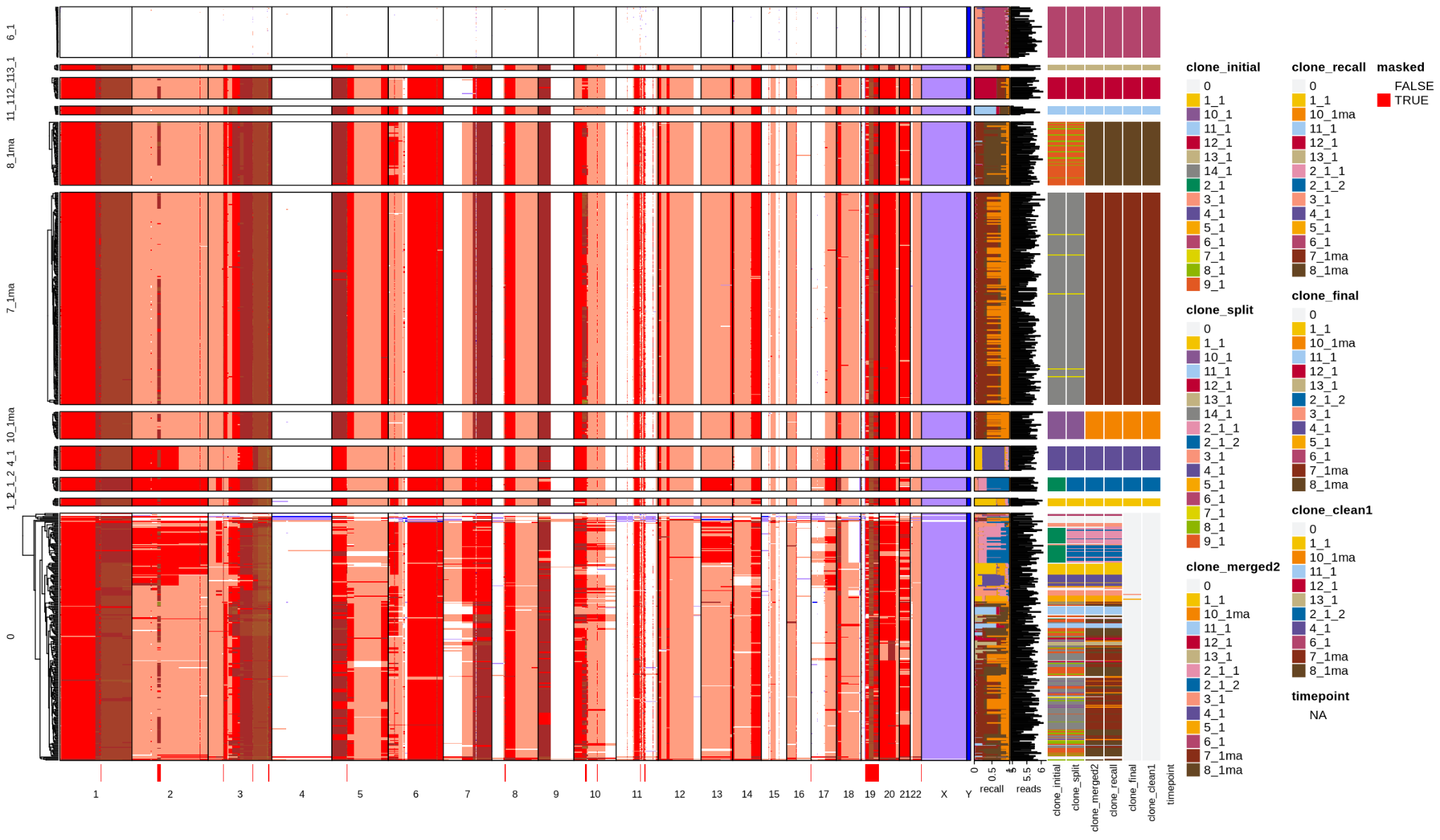

OC7

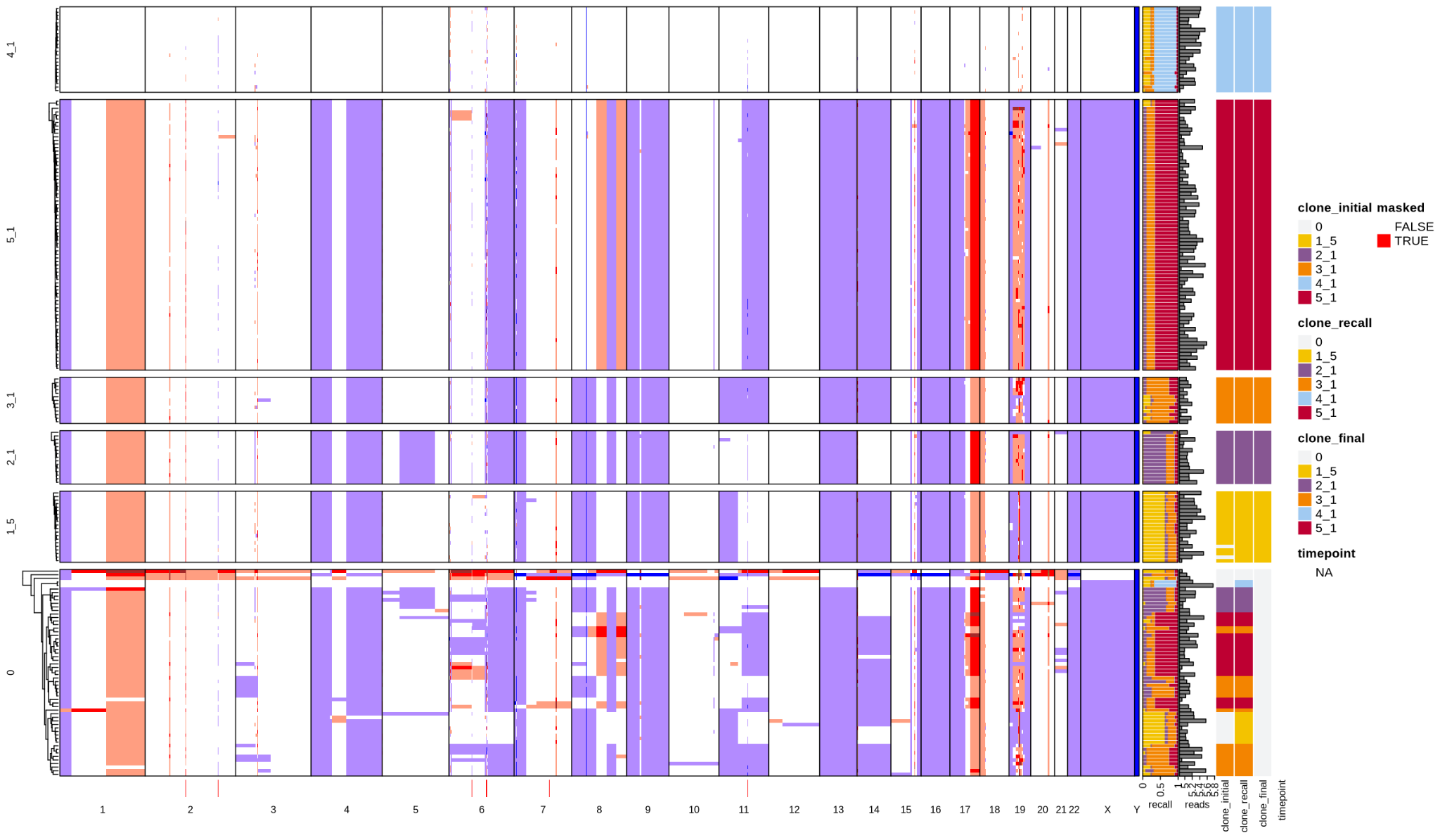

BC11

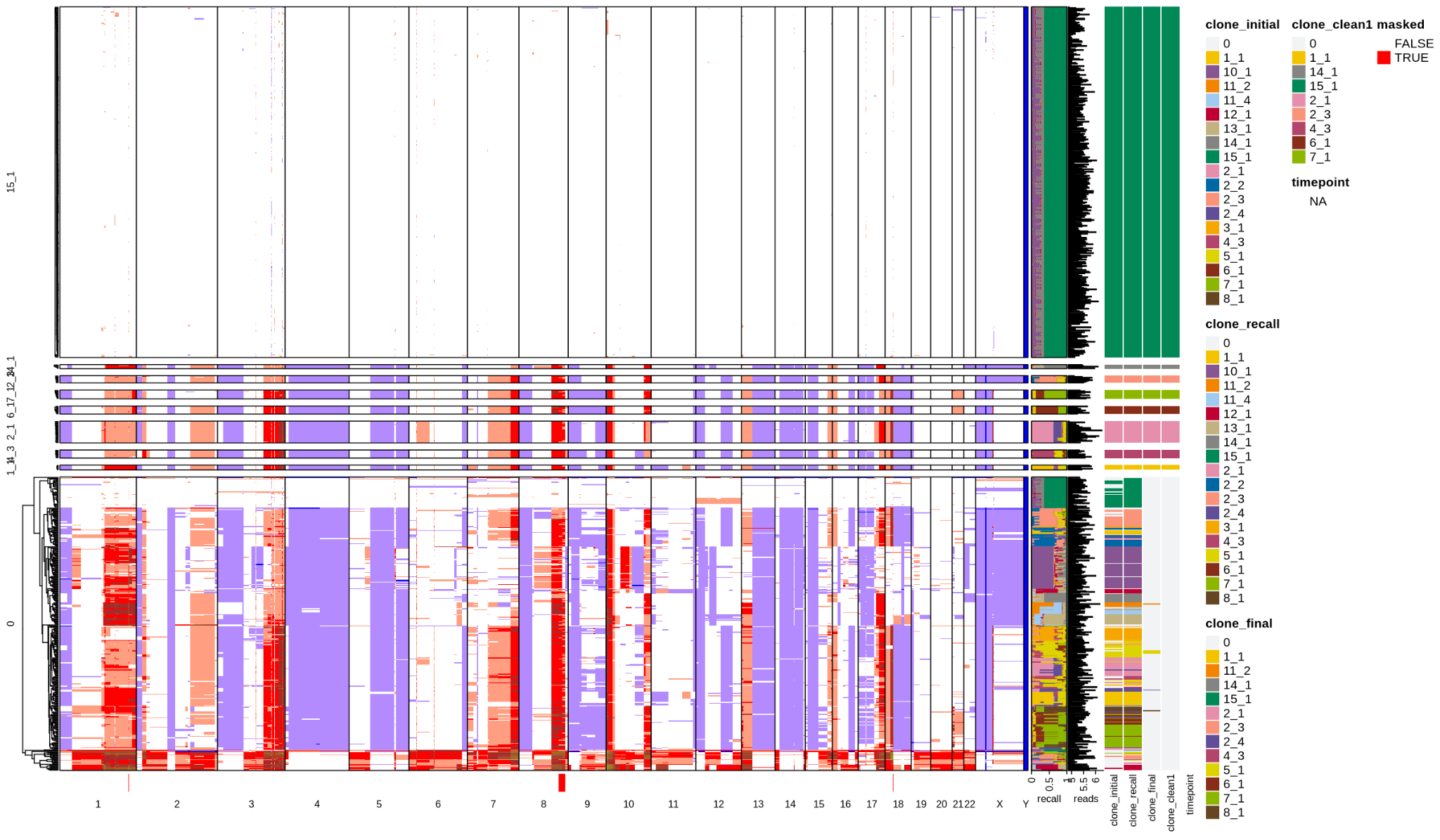

BC13

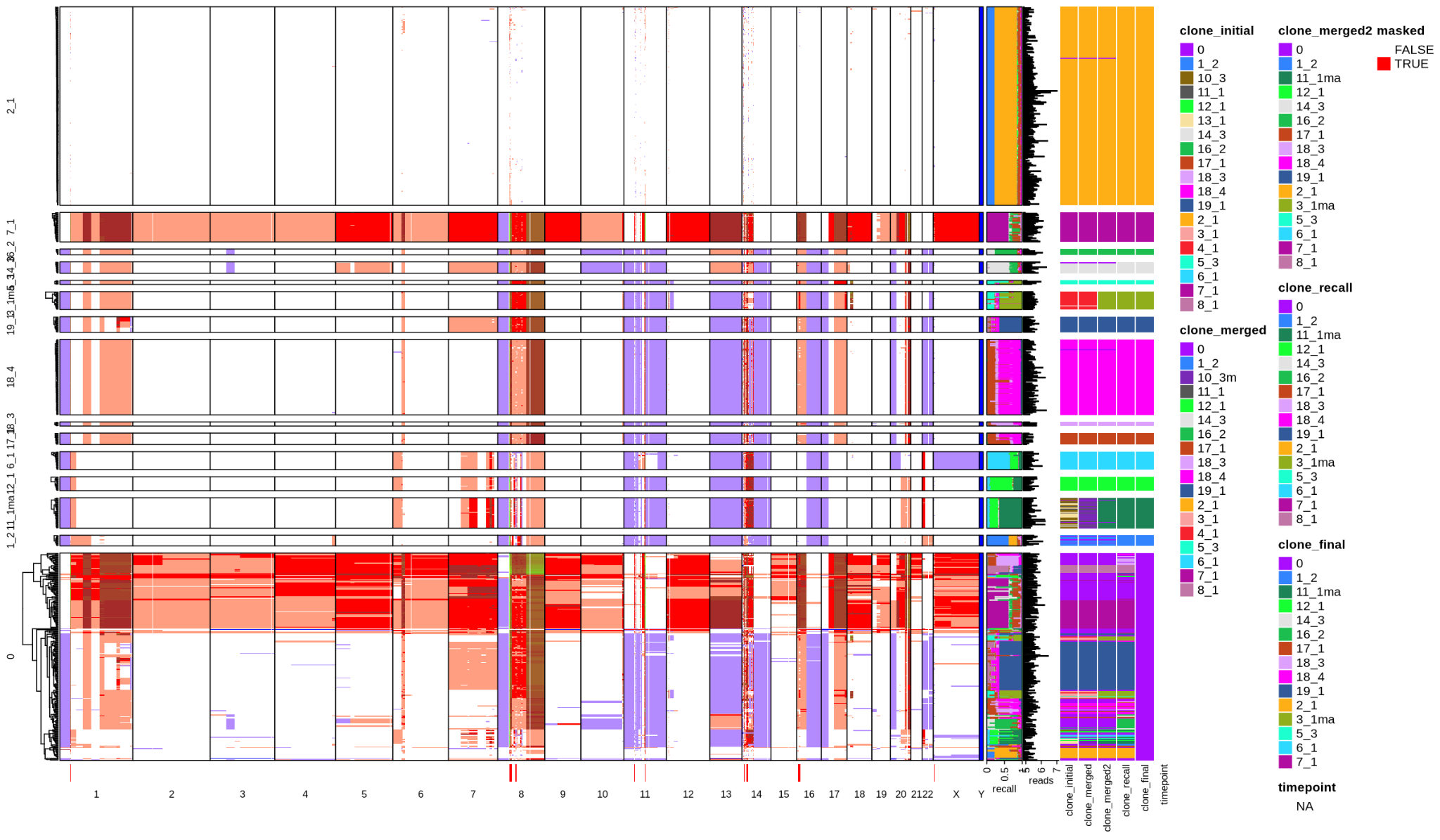

BC16

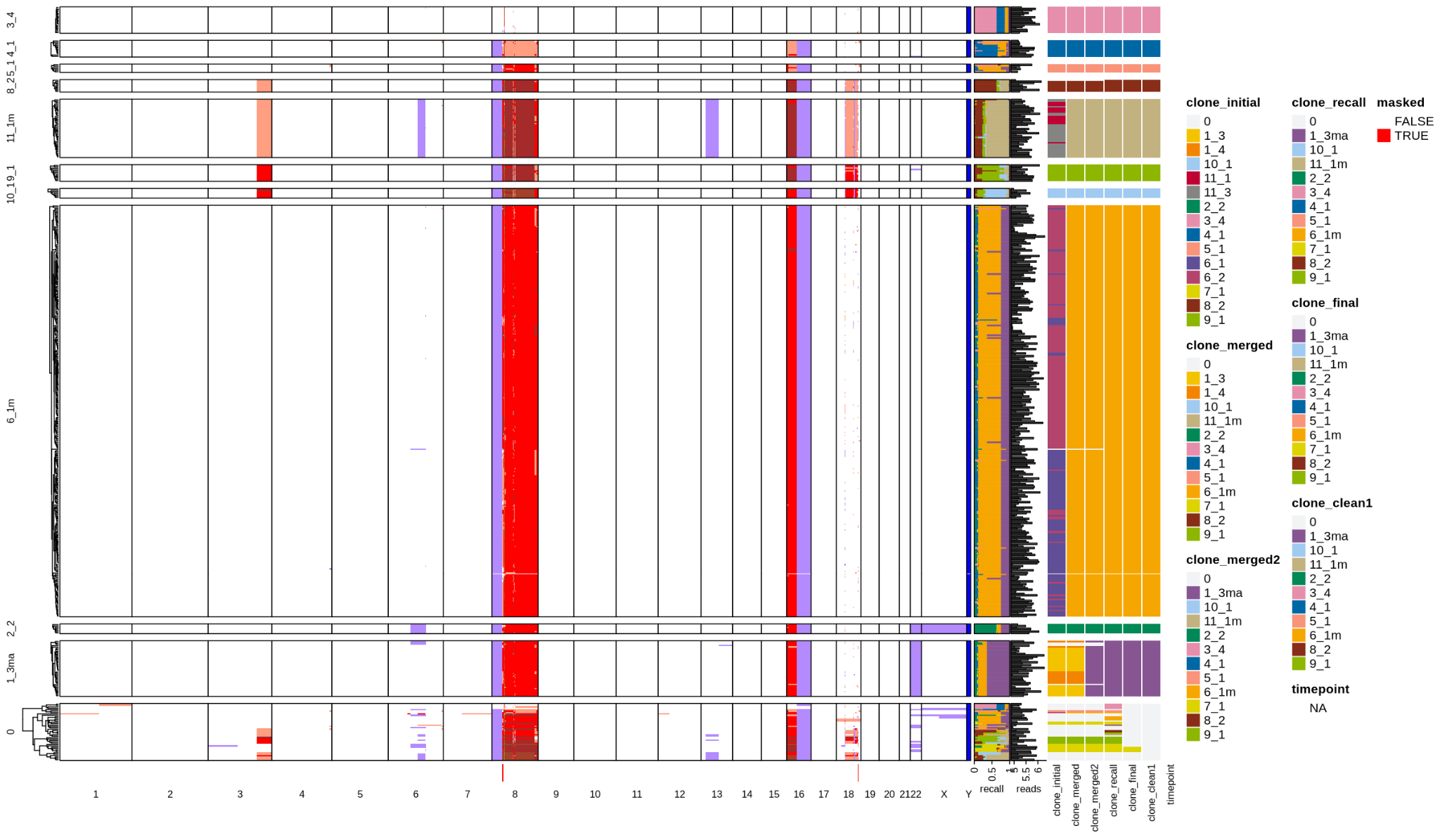

BC19

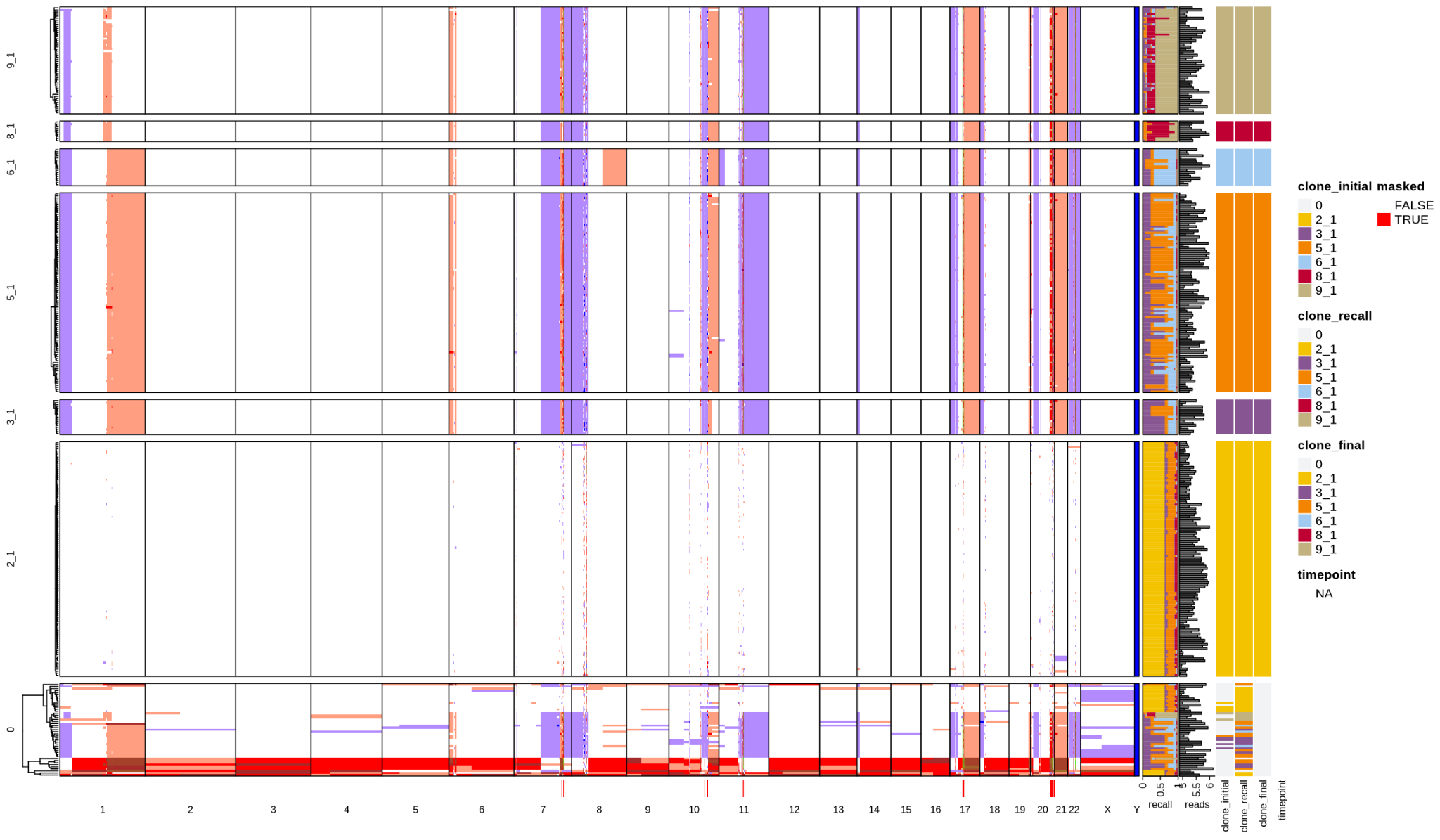

BC20

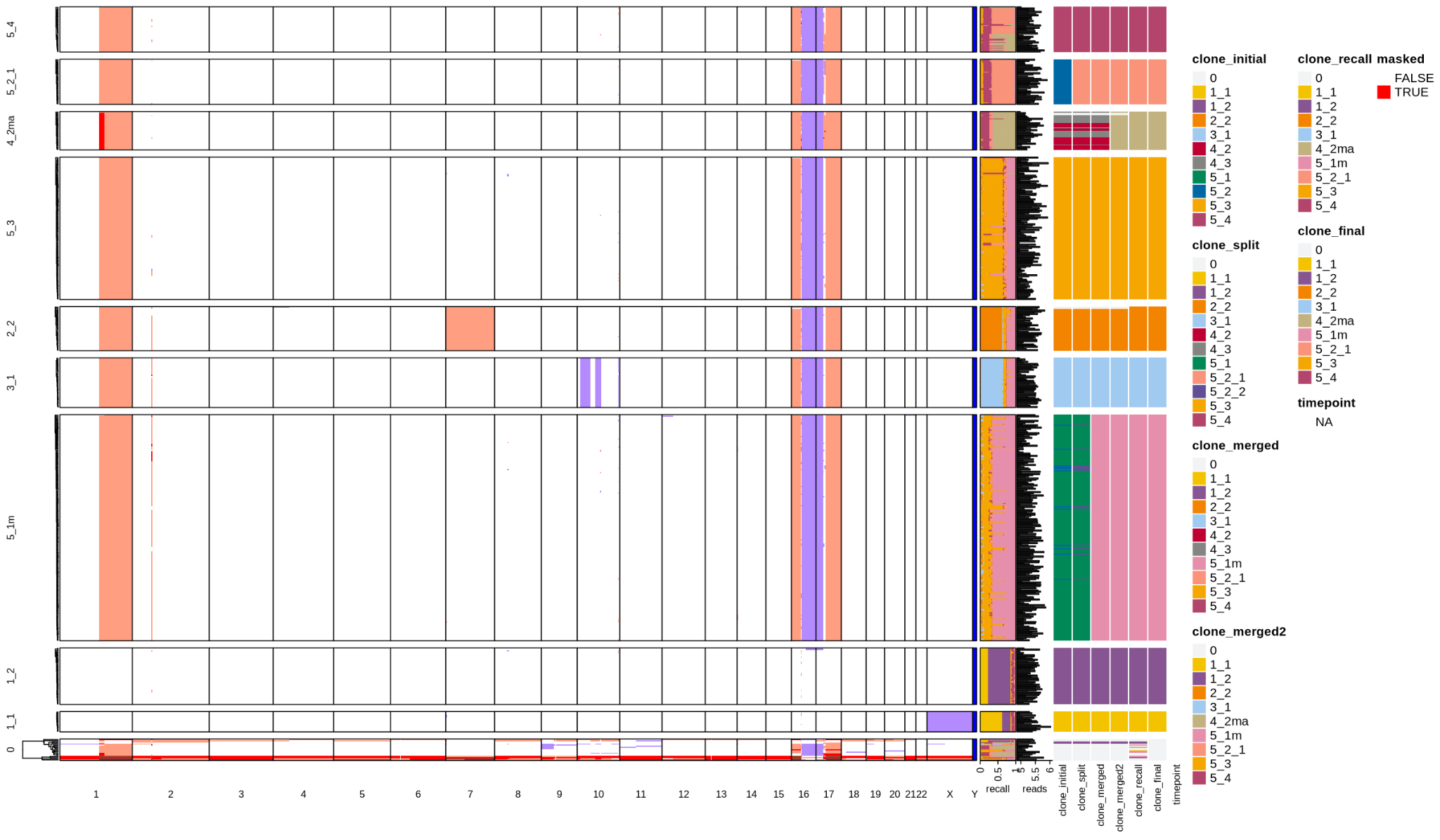

BC26

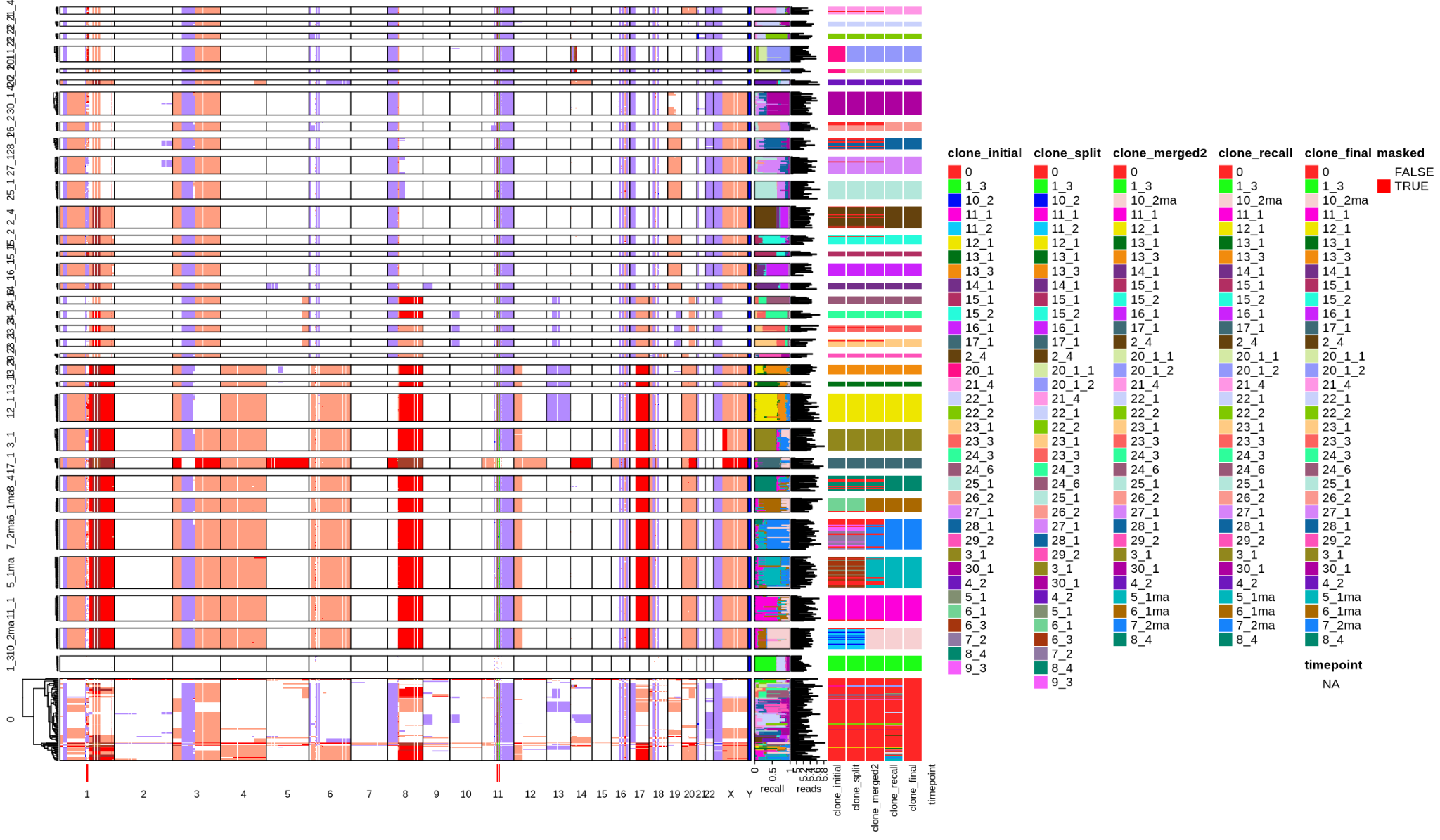

BC28

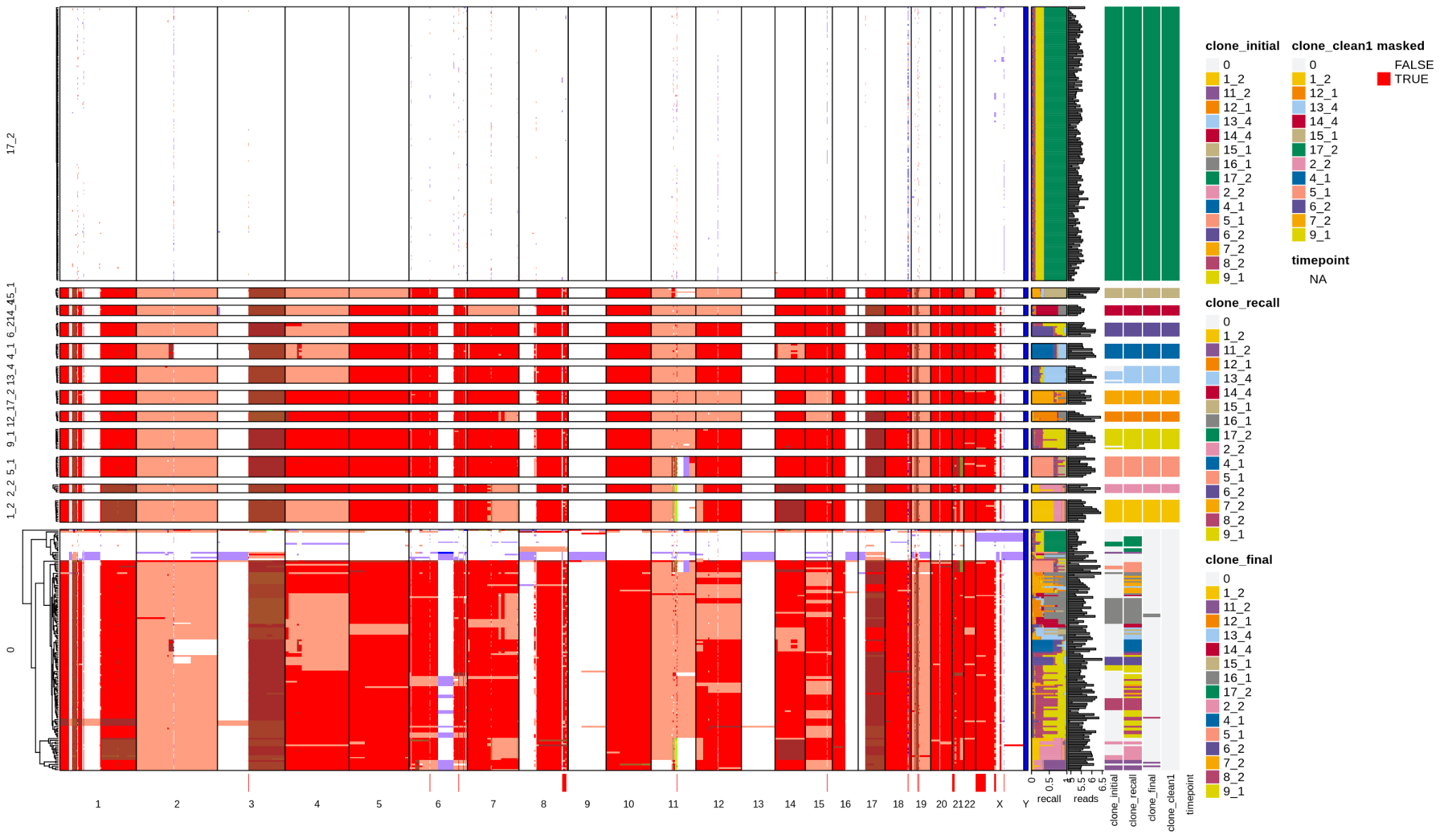

BC30

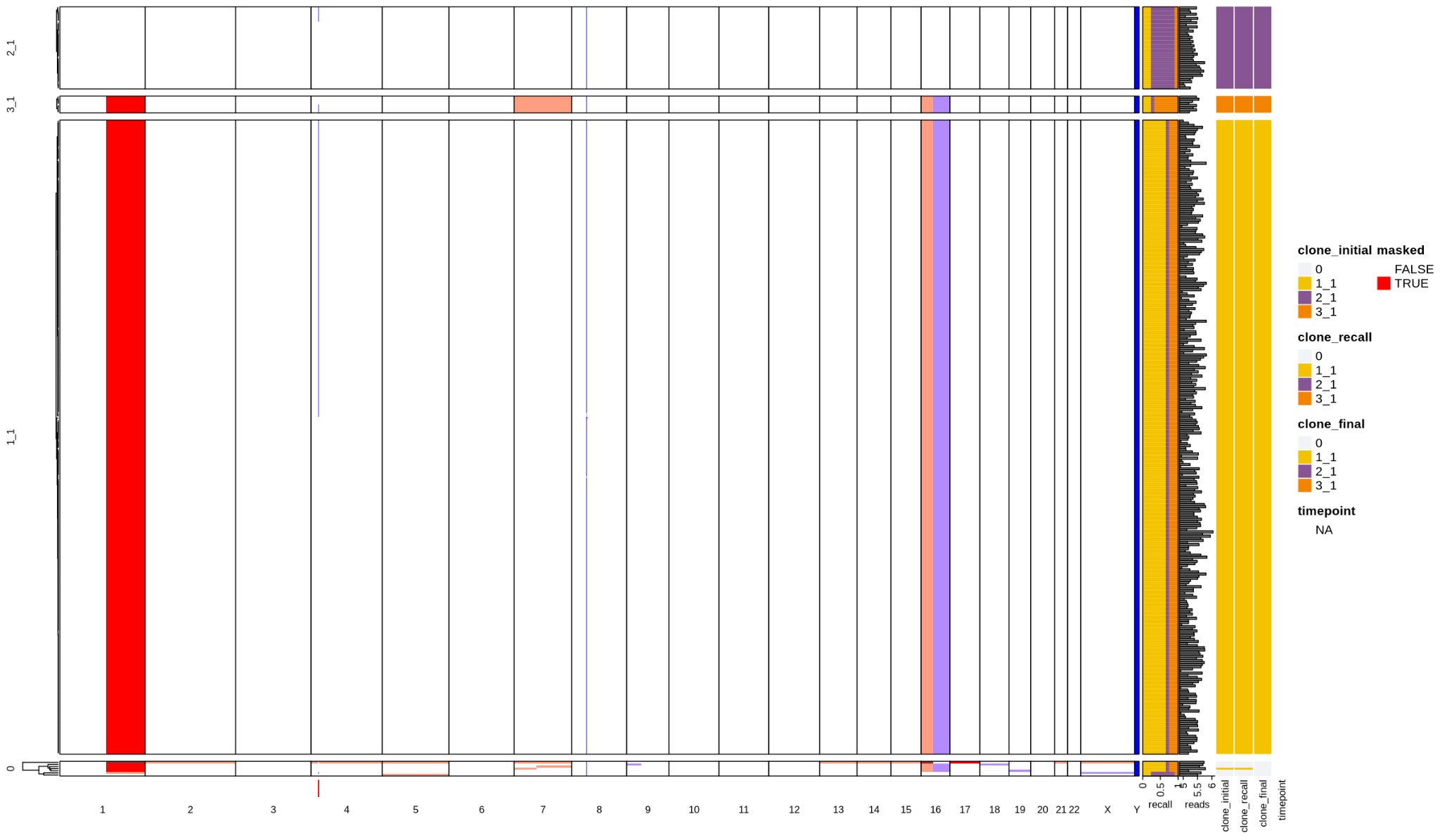

BC38

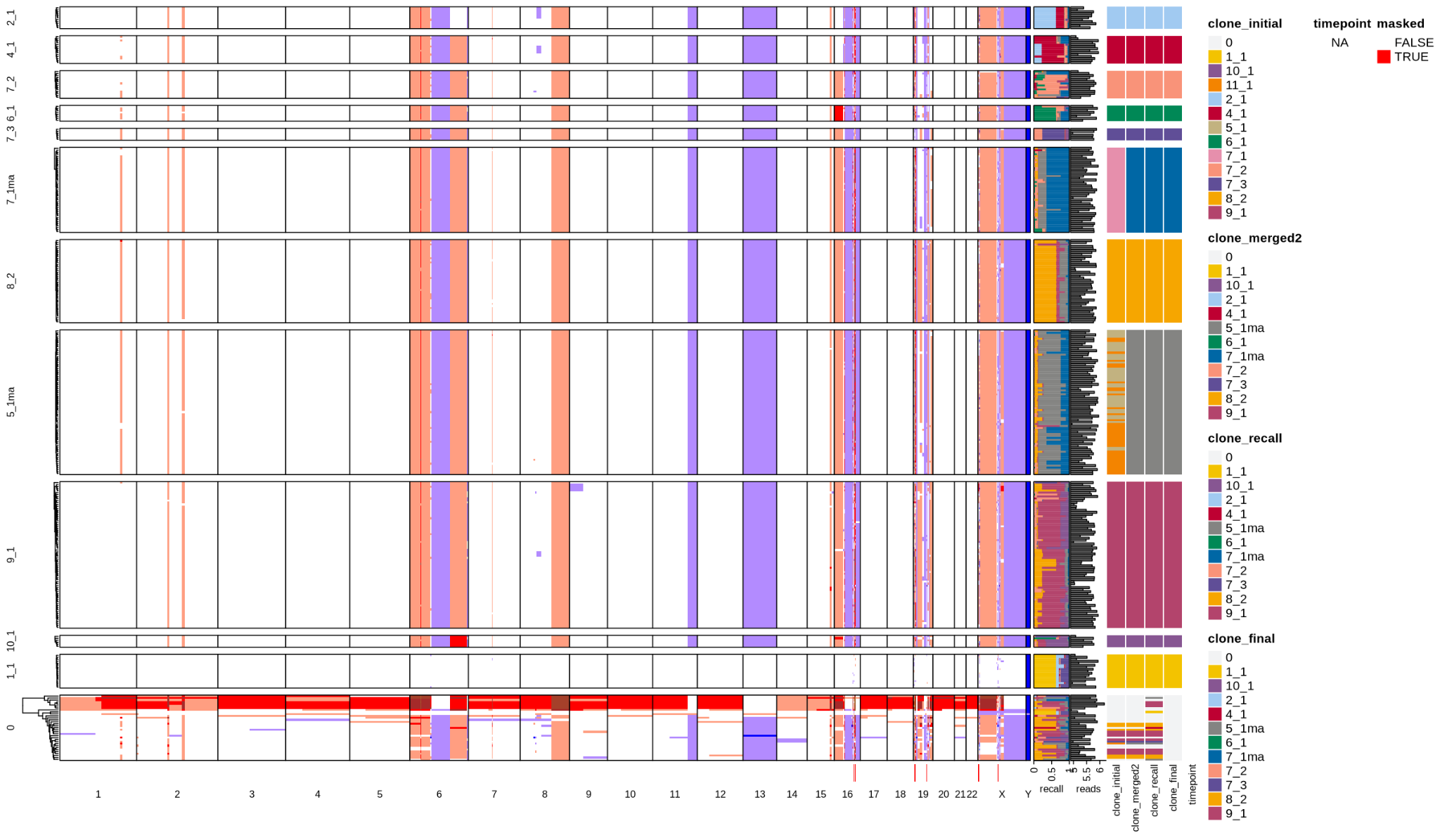

MEL1

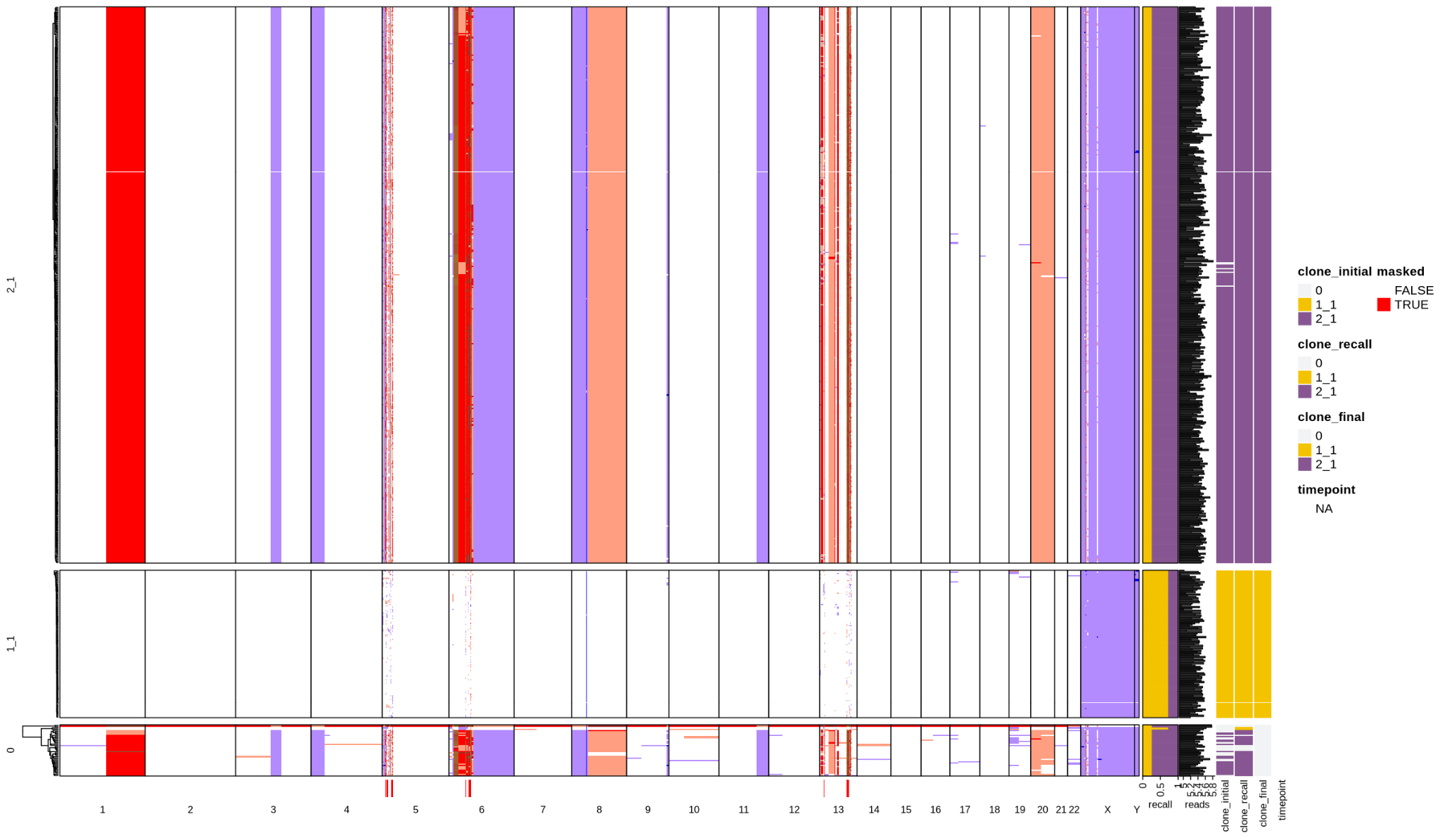

MEL2

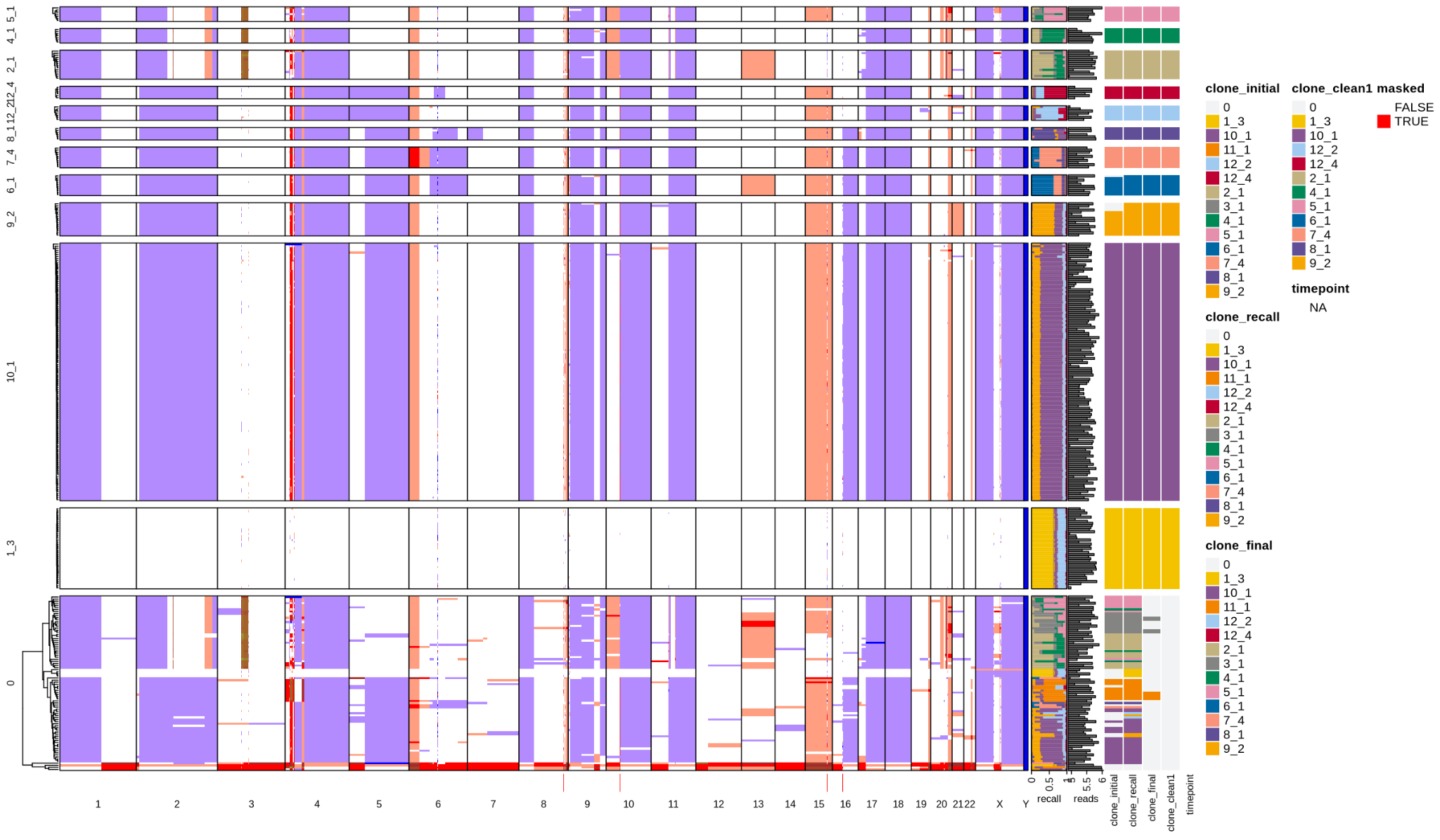

MEL3

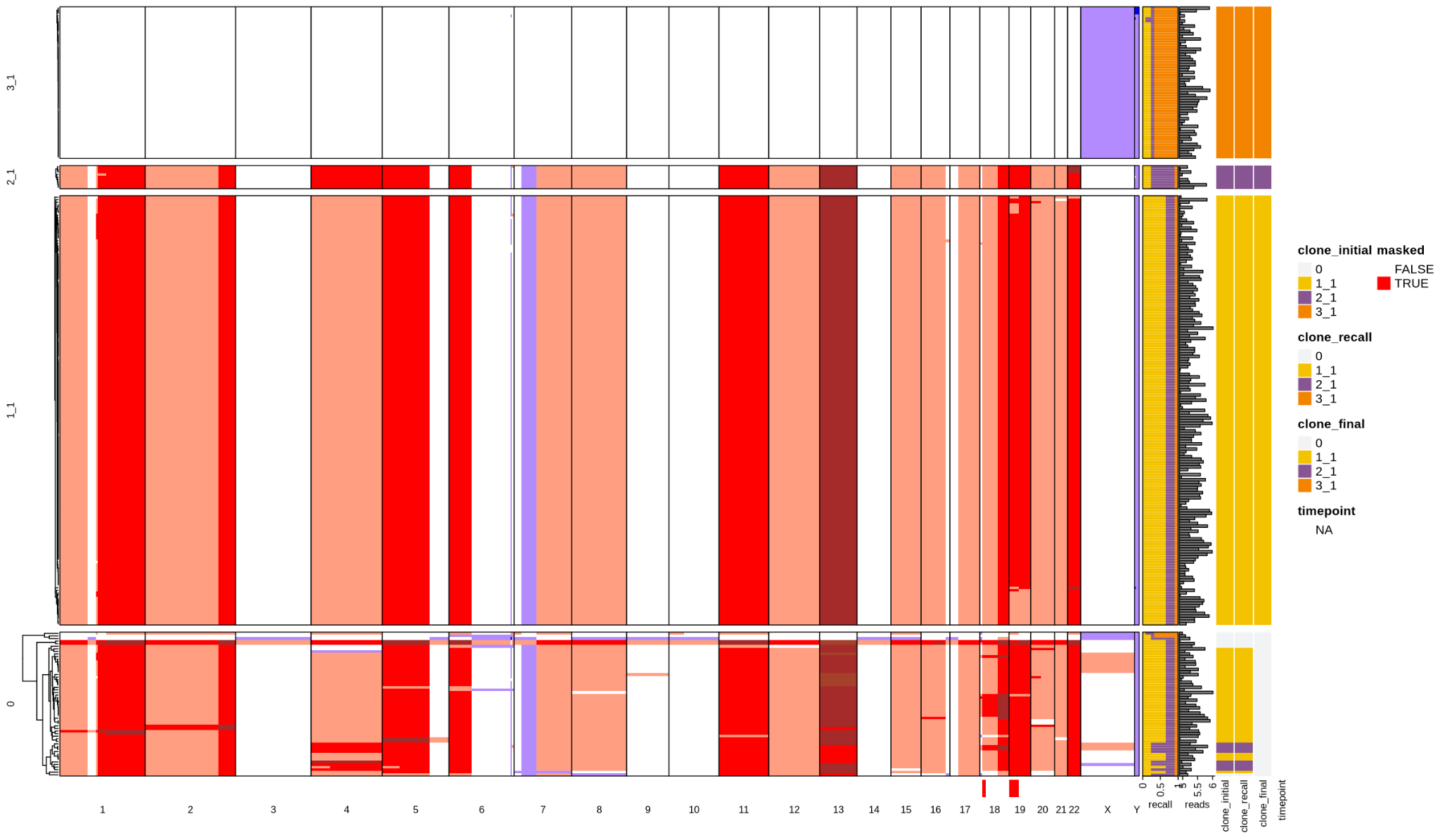

MEL4

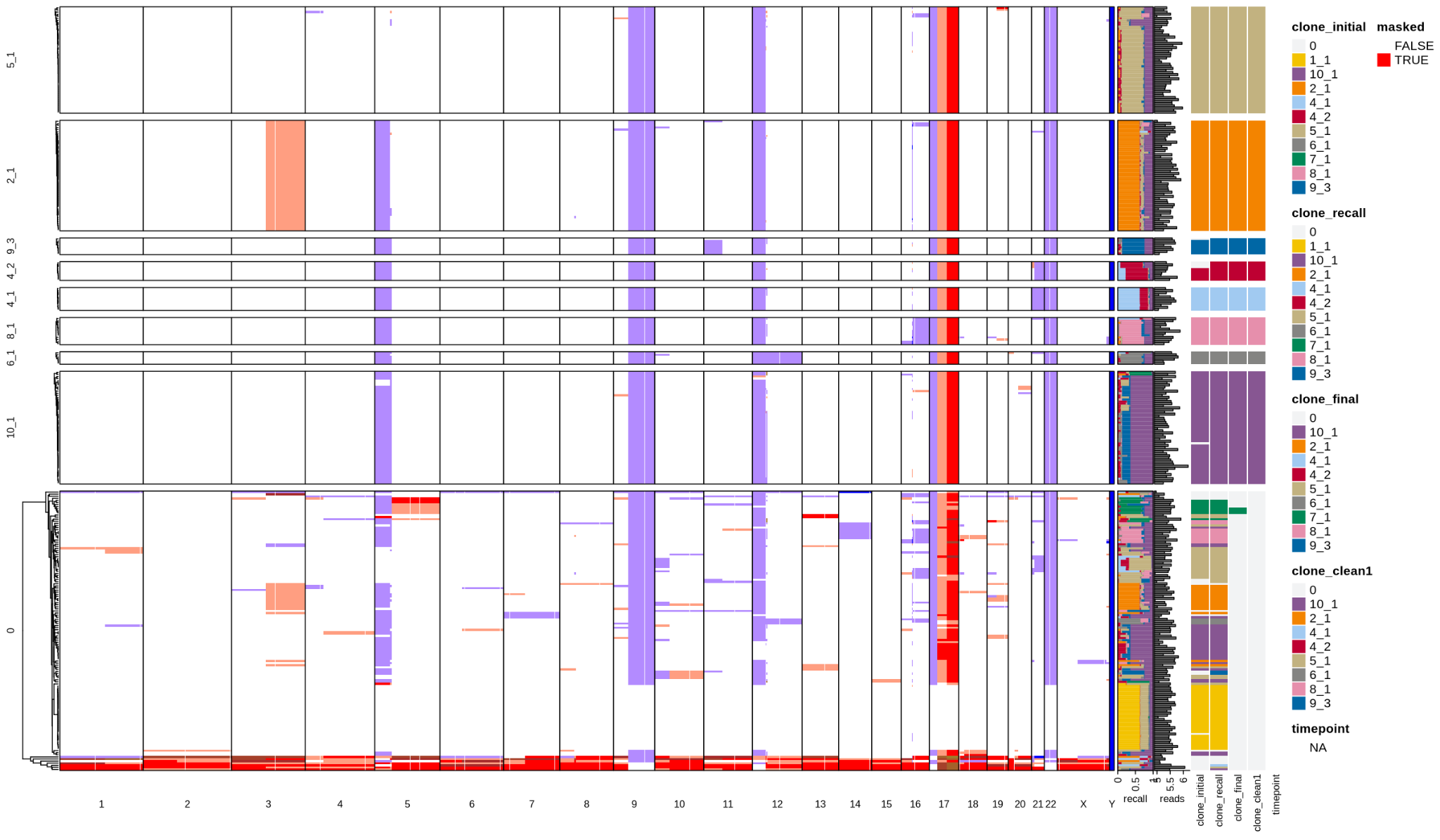

**TRANSIENT**

SRC26

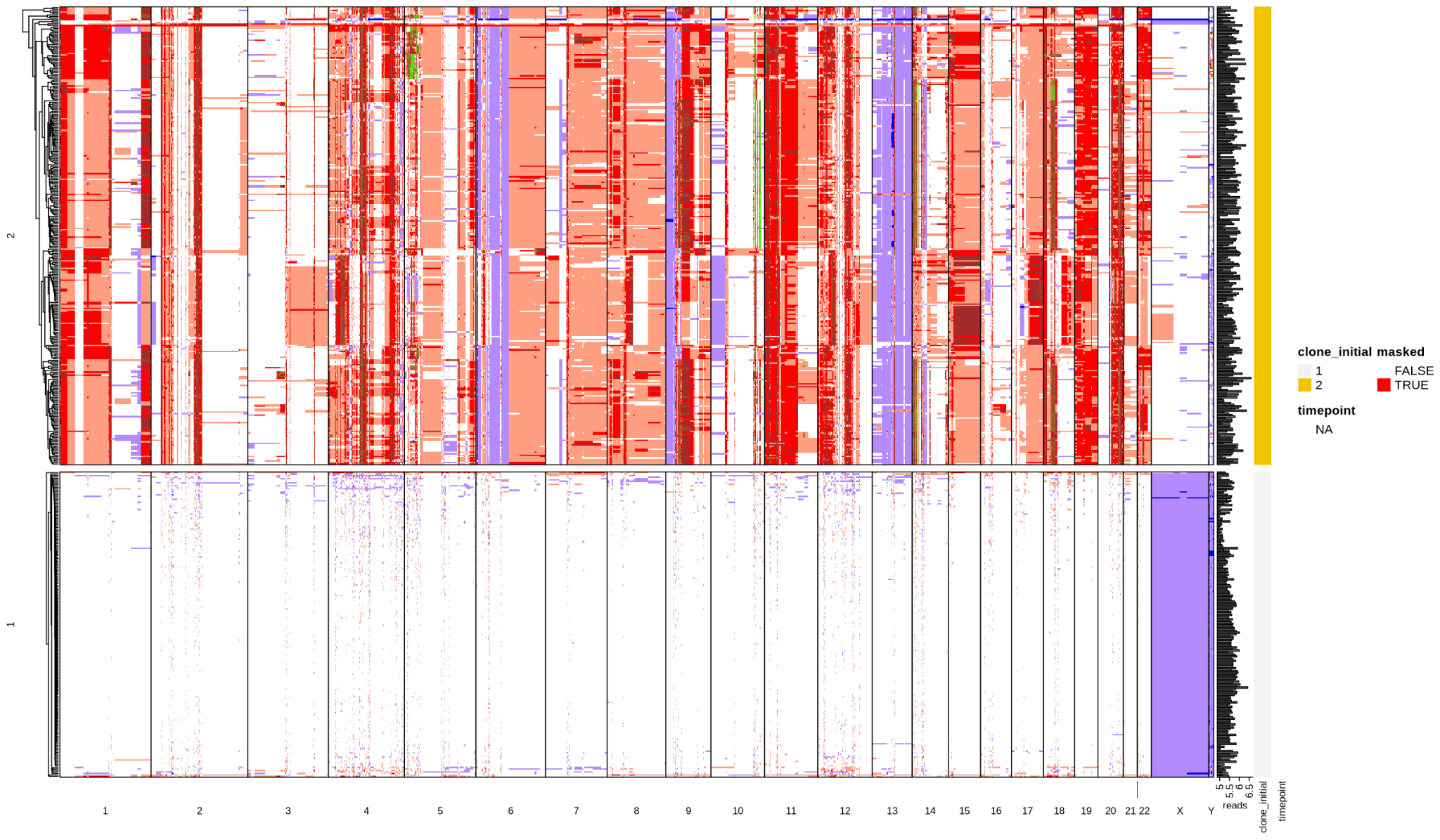

SRC39

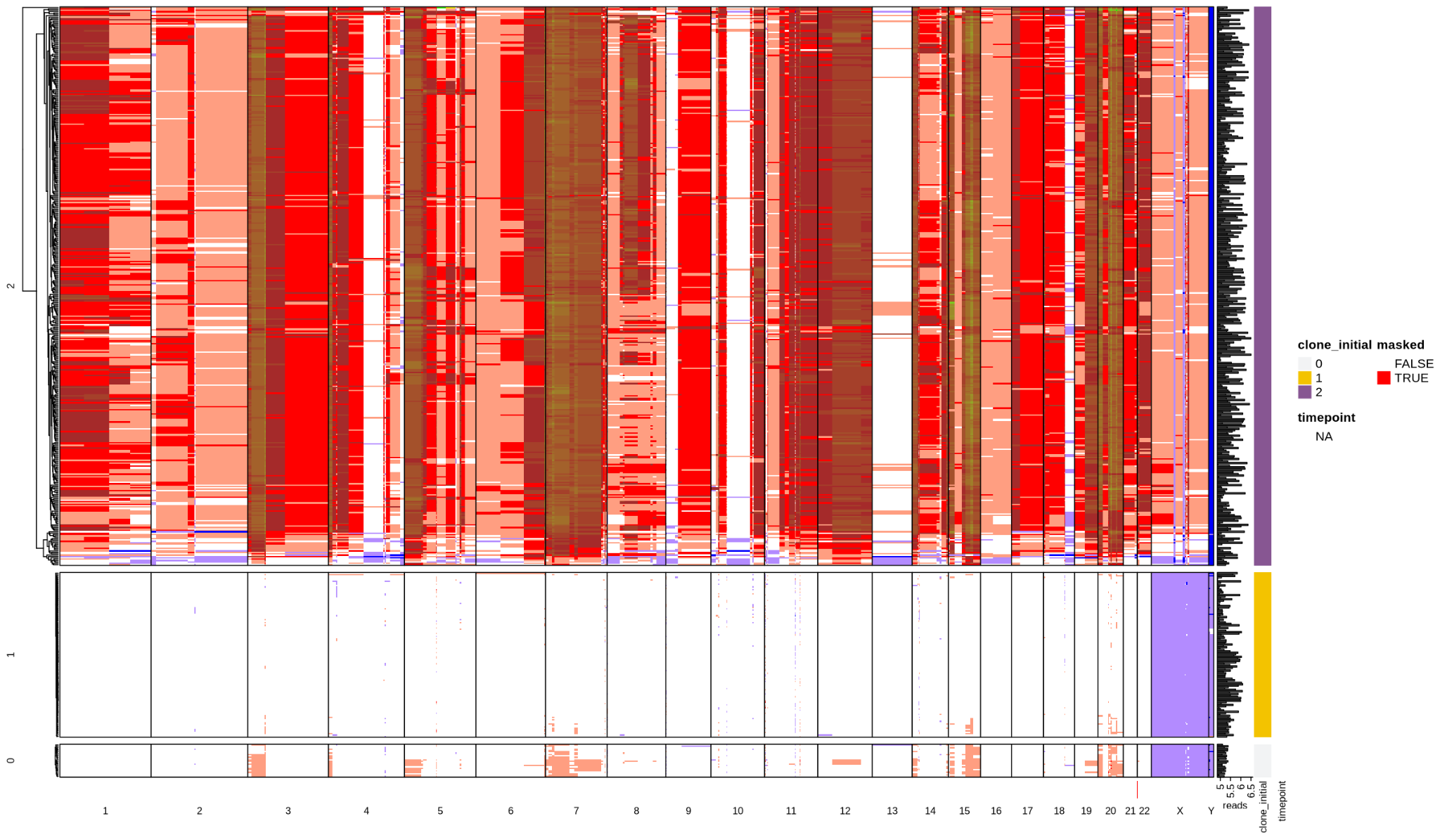

SRC32

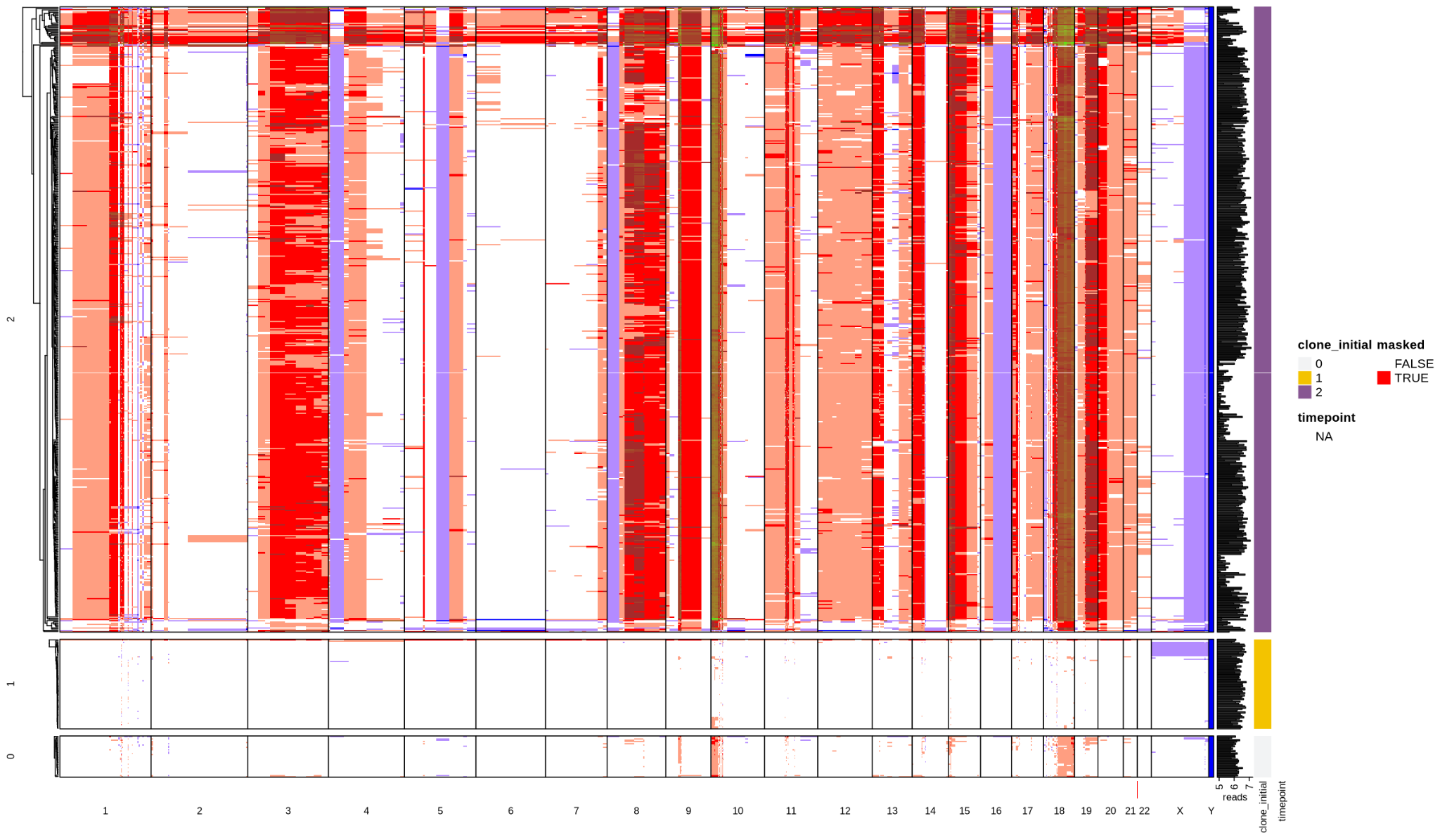

SRC29

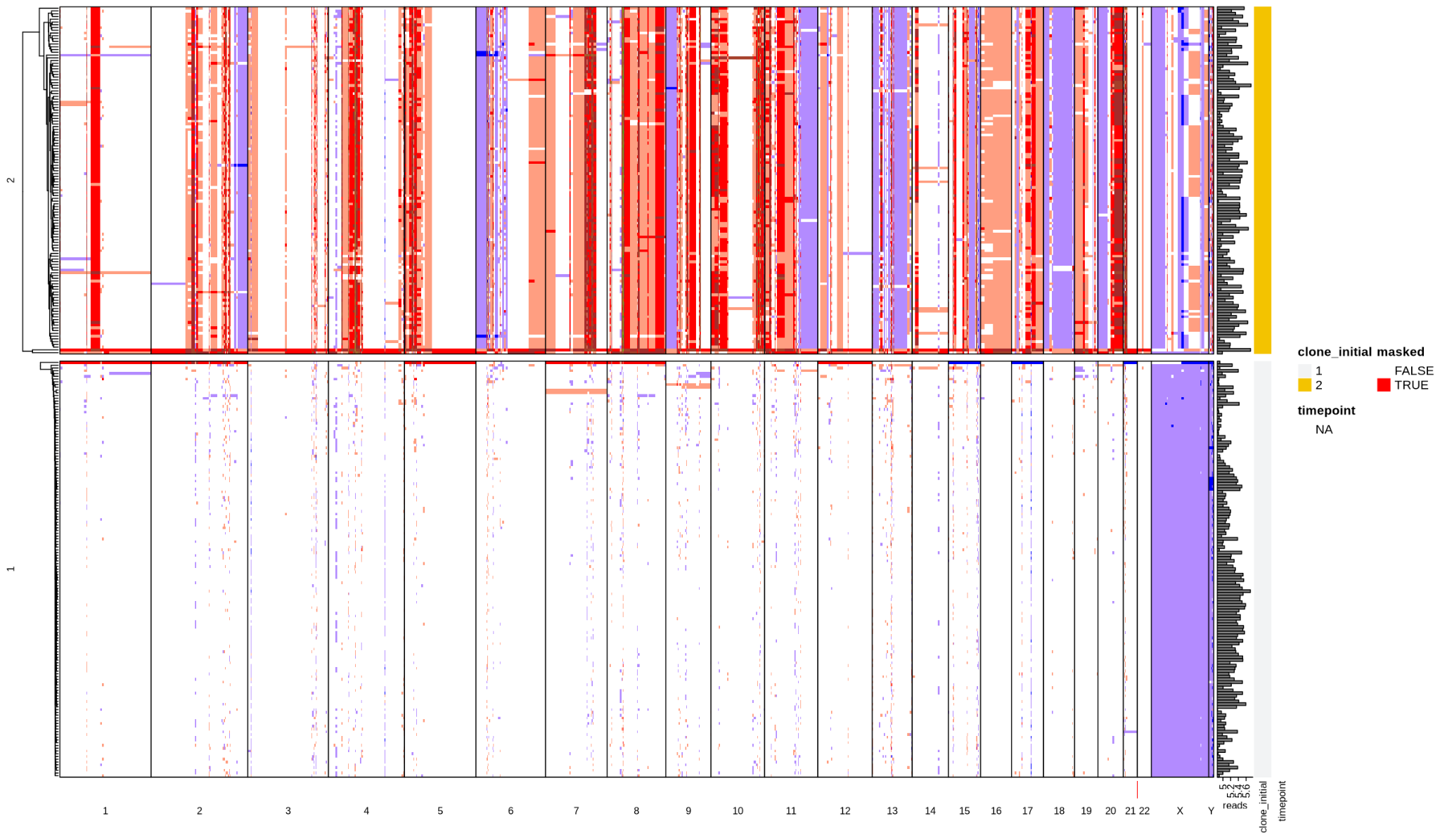

SRC24

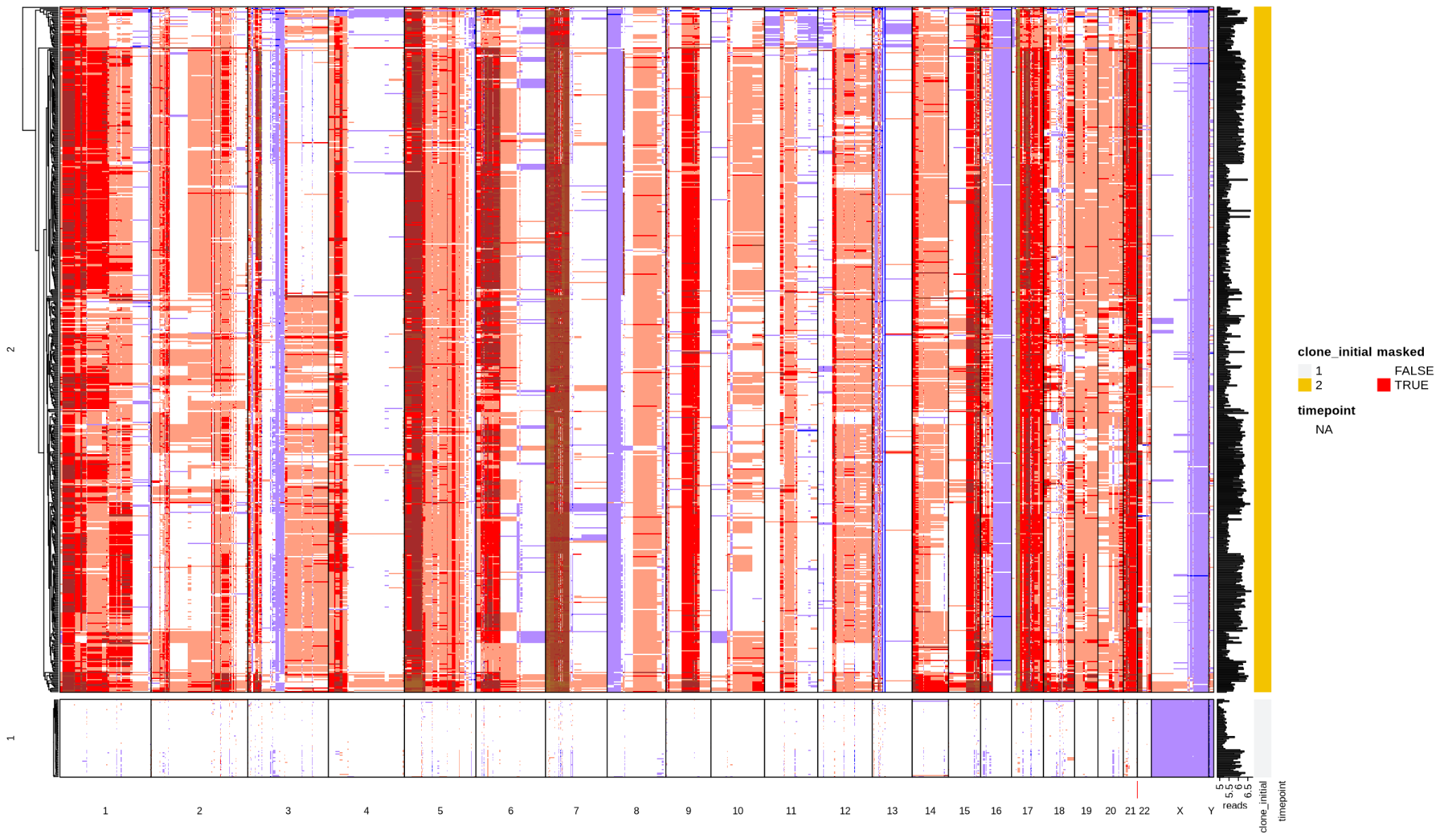

SRC16

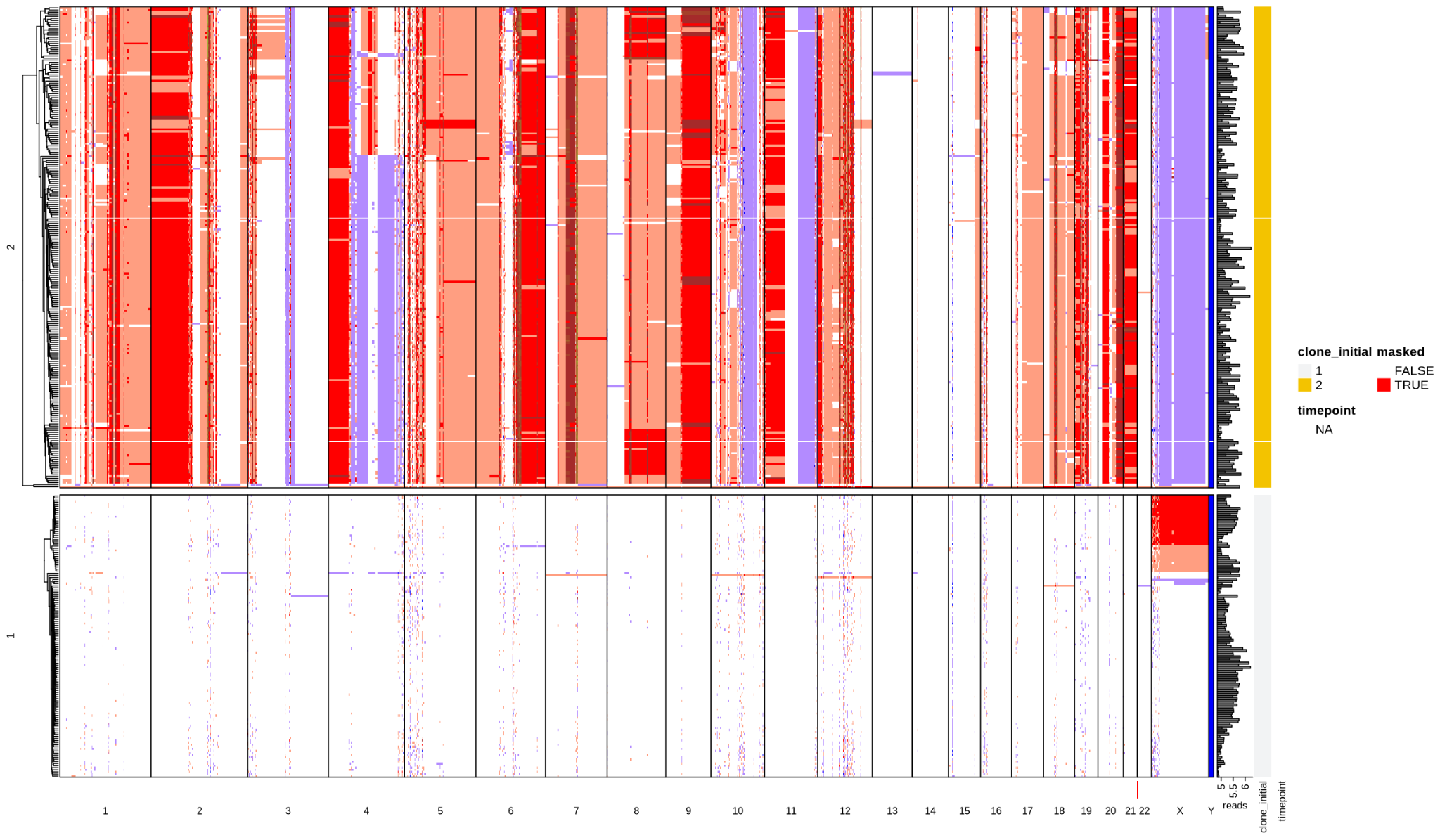

OC3

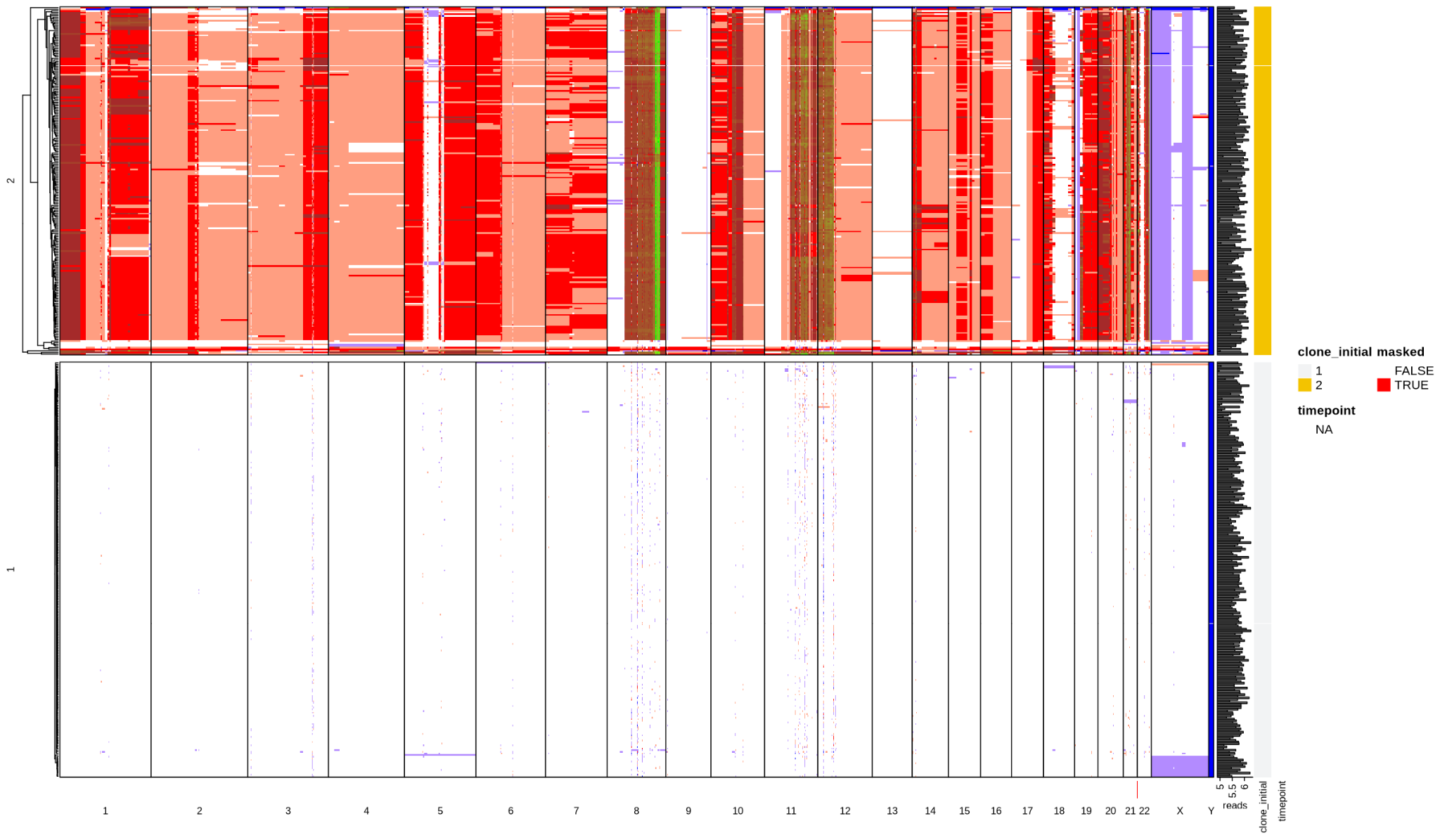

OC6

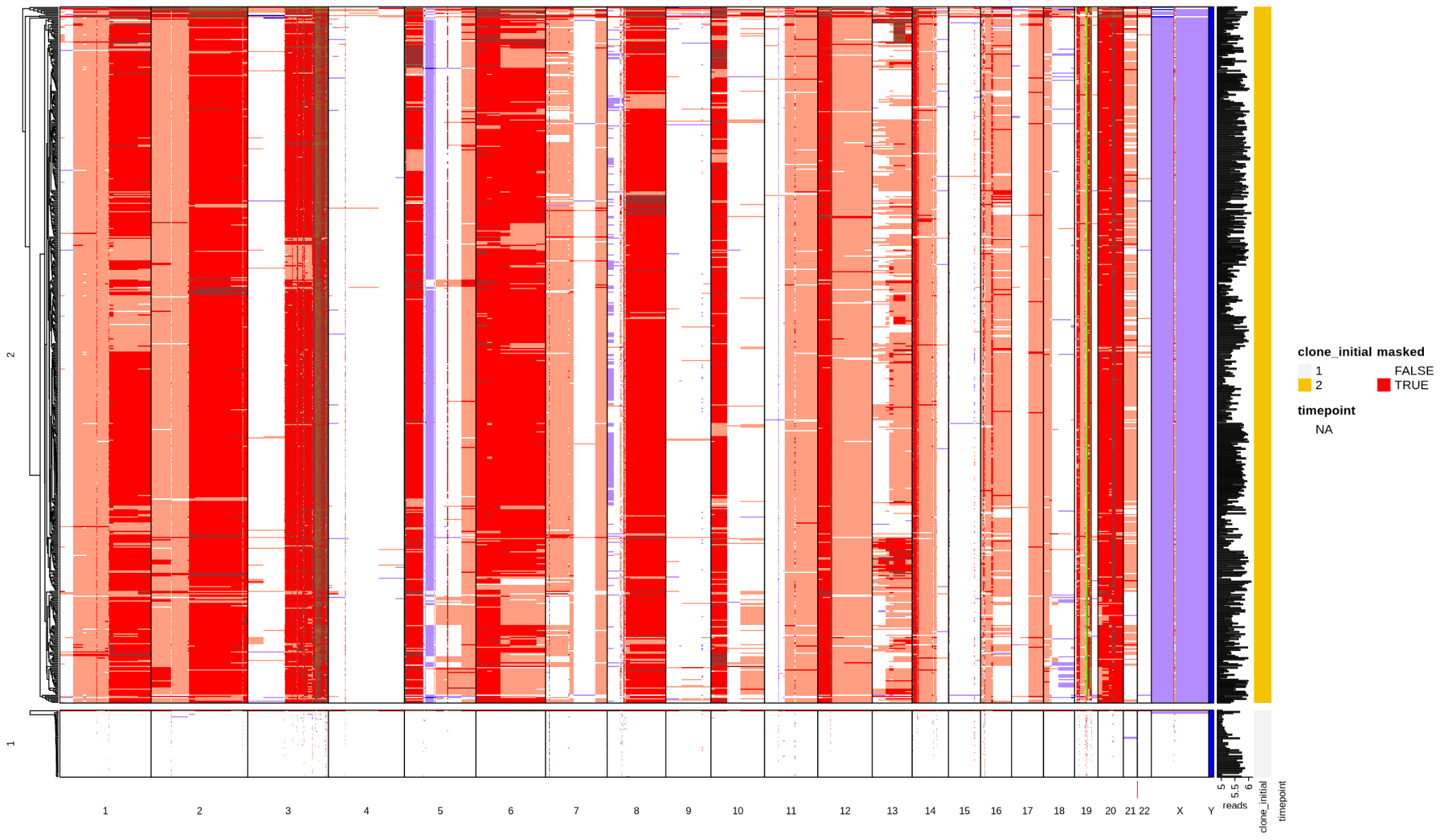

OC8

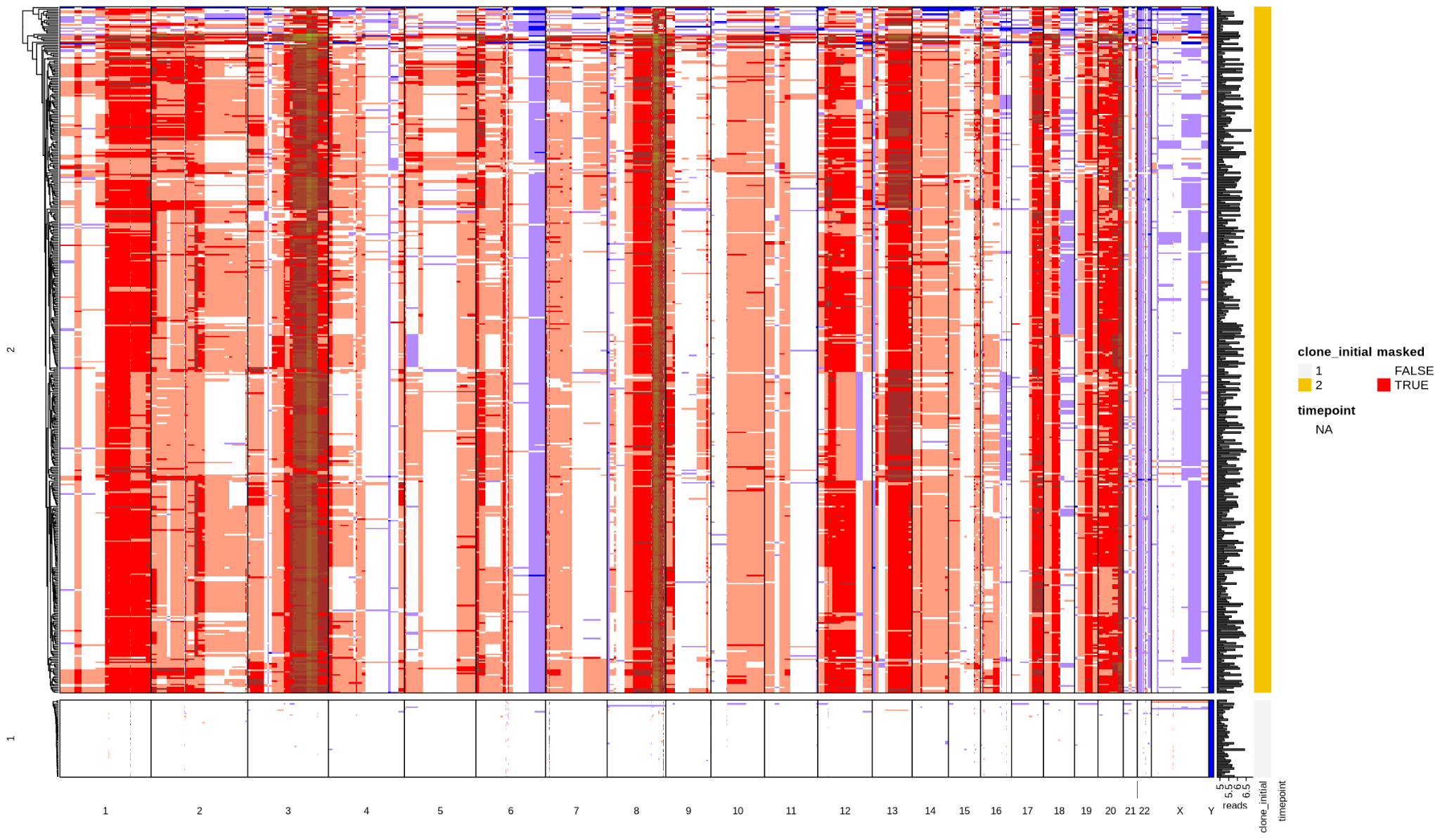

BC8

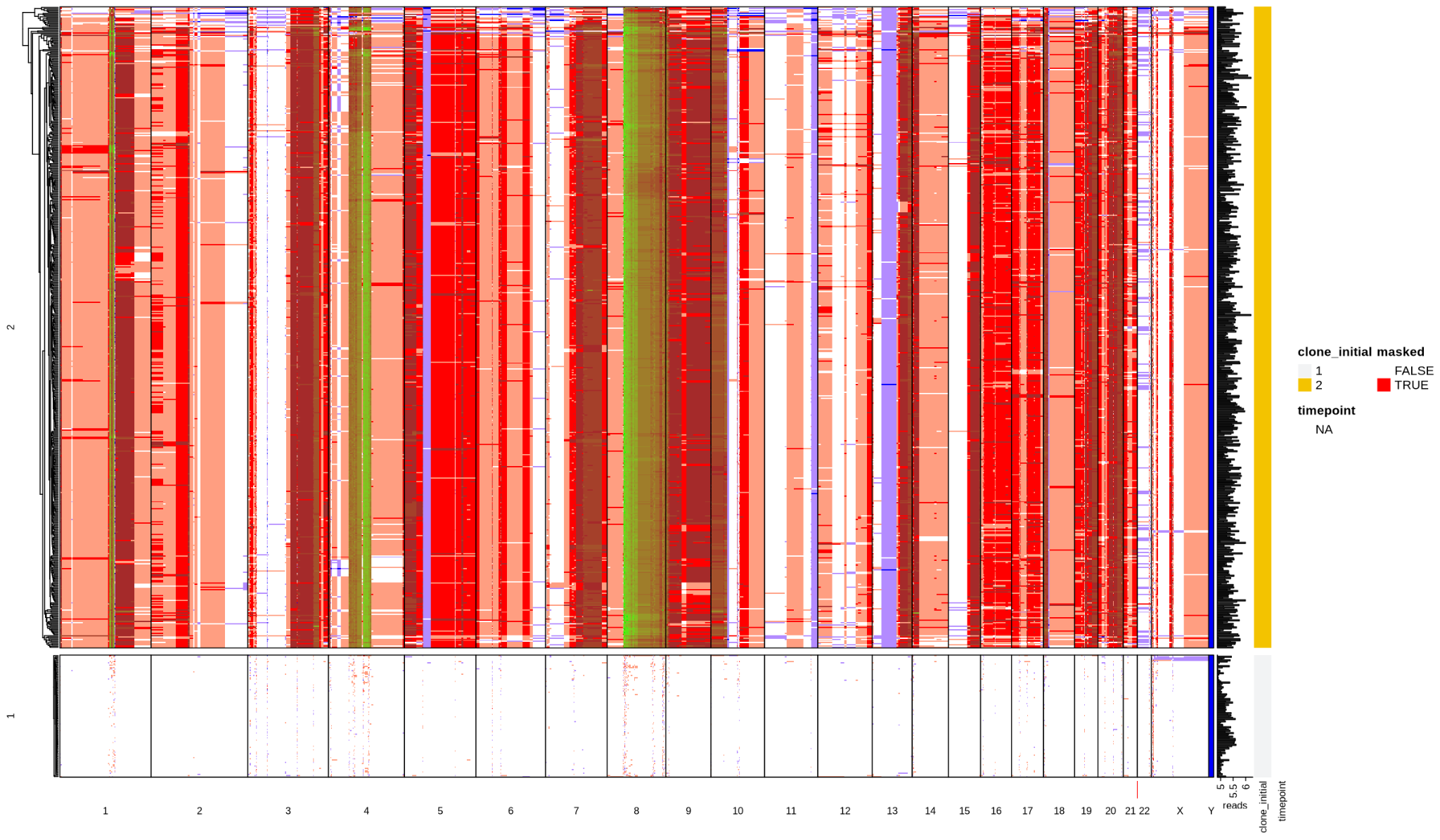
