## Supplementary figures for "Principles of subclonal gene dosage across human cancer"

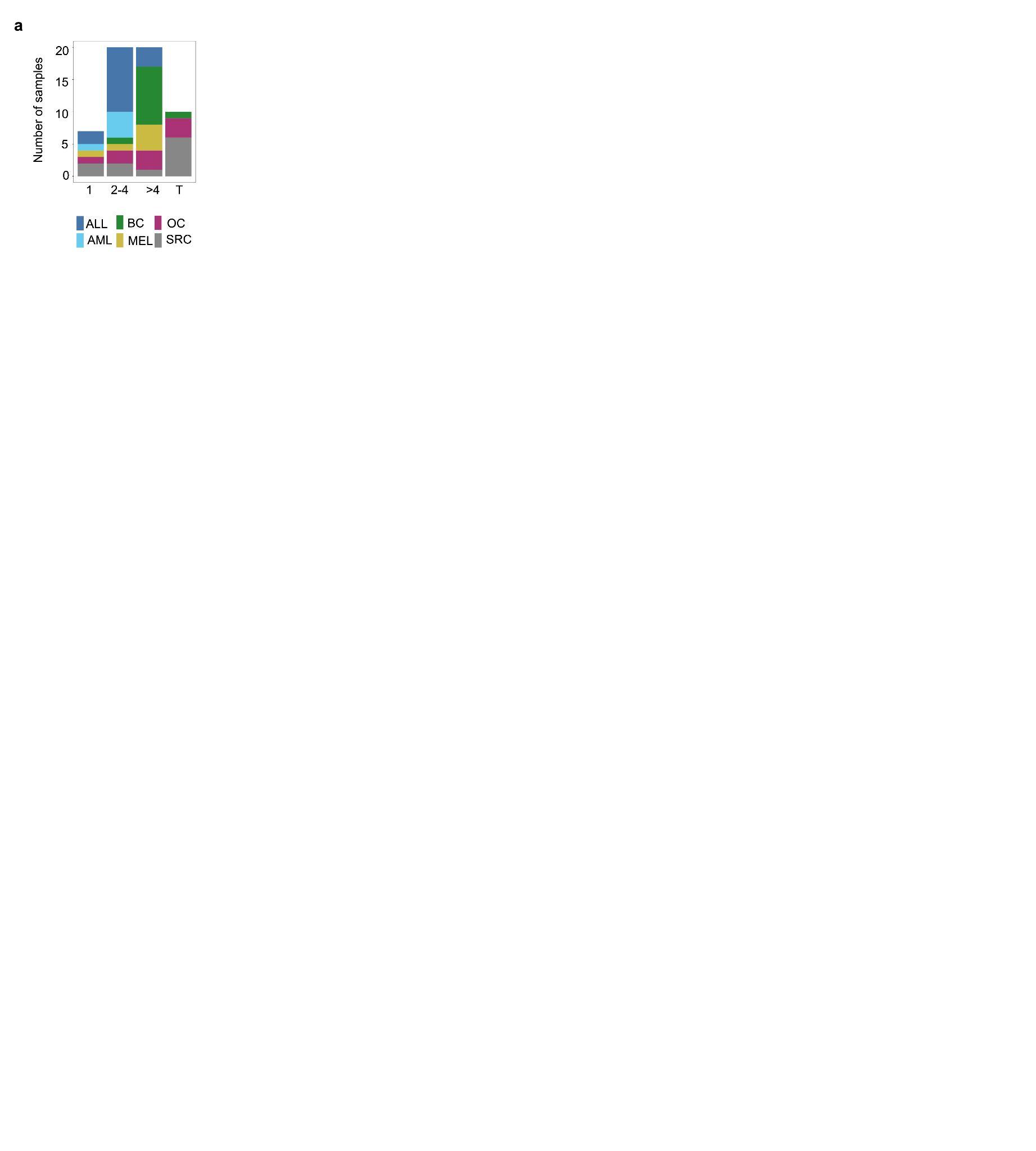

**Supplementary figure 1.Subclonal composition across the dataset. a:** X axis shows number of subclones across patients, colored by cancer type. T indicates *transient* samples.

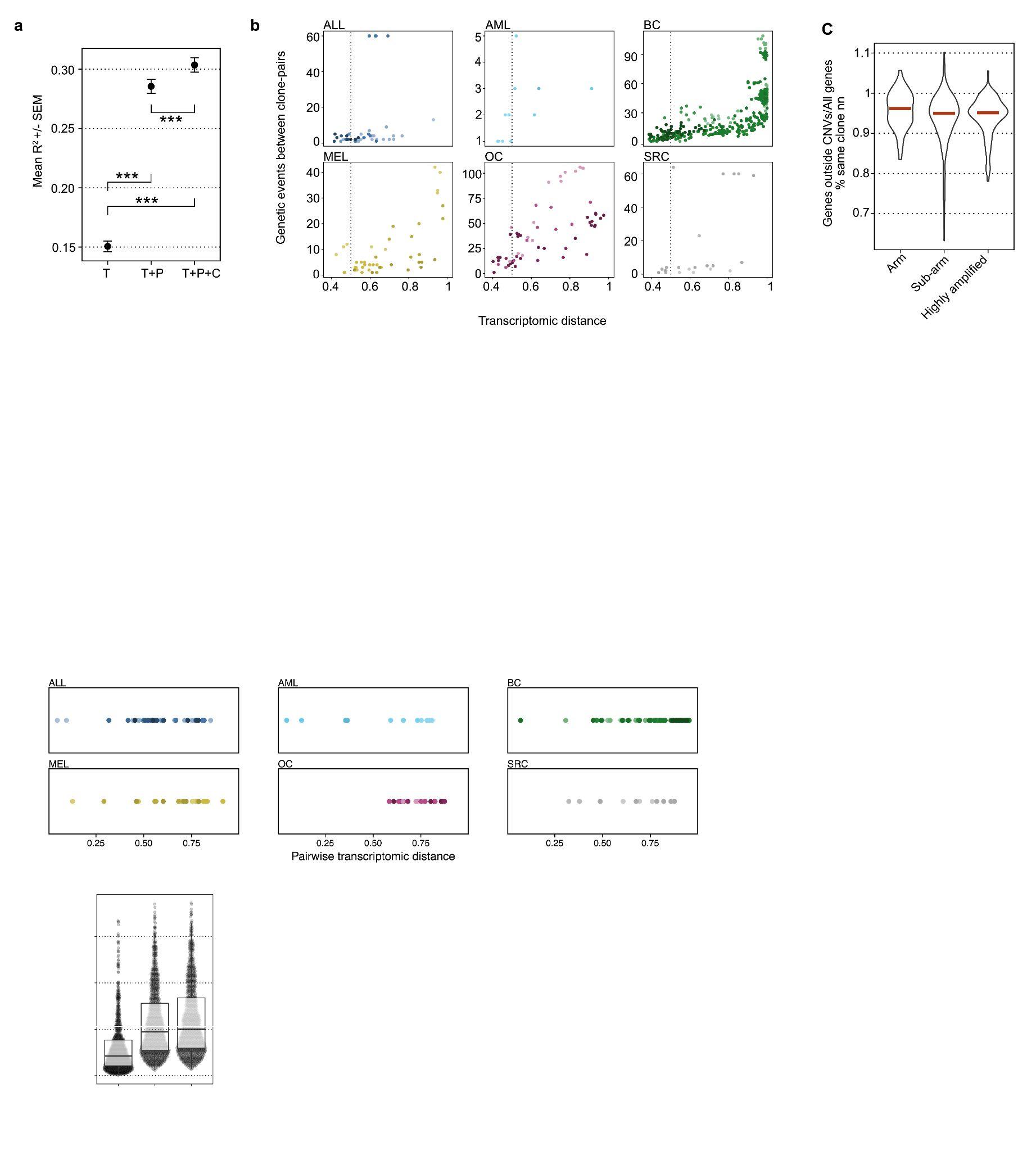

**Supplementary figure 2. Genetic constraints on phenotype. a**: Mean R^2^ from a linear model given T (tumor type), P (patient) and C (clone) of all tumor cells from patients that have subclones Error bars indicate standard deviation of the mean (SEM). Wilcoxon rank-sum test was used for p value statistics. All p values were <2.2e-16. Tumor type explains most of the variance, while subclonality increases R2 by approximately 5% of total, or 10% of the additional variance explained by the patient identity. **b**: Pairwise analysis of clone pairs, genetic events plotted against transcriptomic distance. Vertical dotted line indicates 0.5.

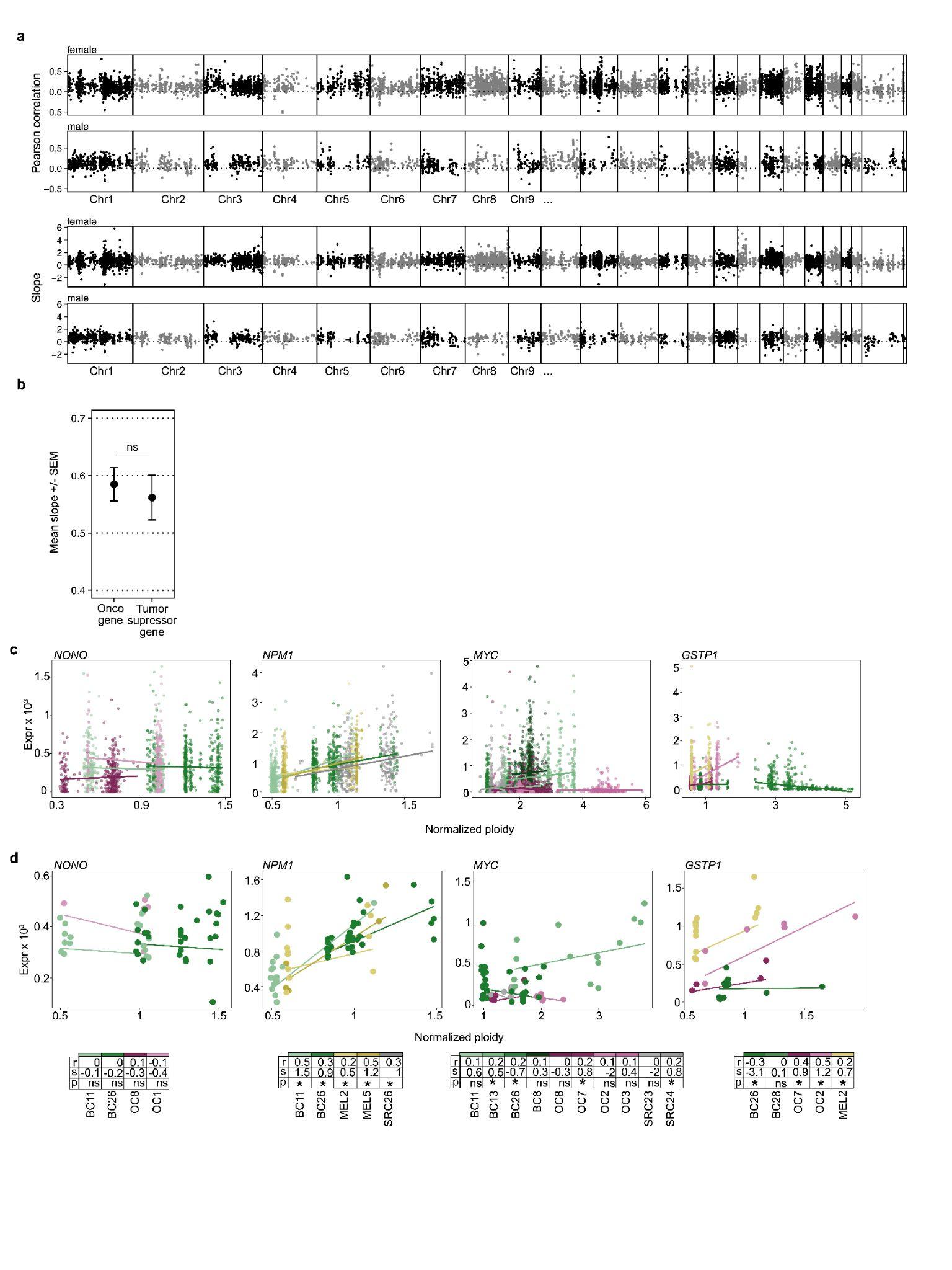

**Supplementary figure 3. Genome-wide dosage effects across cancer types. a**: Top: Pearson correlation across all chromosomes split by sex of the patient. Bottom: Normalised coefficient (slope) across all chromosomes split by sex of the patient **b**: Mean slope +/- standard error of the mean (SEM) of oncogenes and tumor suppressor genes. P values were calculated with a Wilcoxon rank-sum test, significance threshold was at 0.05. **c**: Dosage effects of individual genes. The slope of each line is from single-cell robust linear models and the data shown are single cells. Table values are Pearson correlation (r) and Pearson p value (p) and linear model slope (s) based on single-cell data. **d**: Same as in c except the dots shown are subclones.

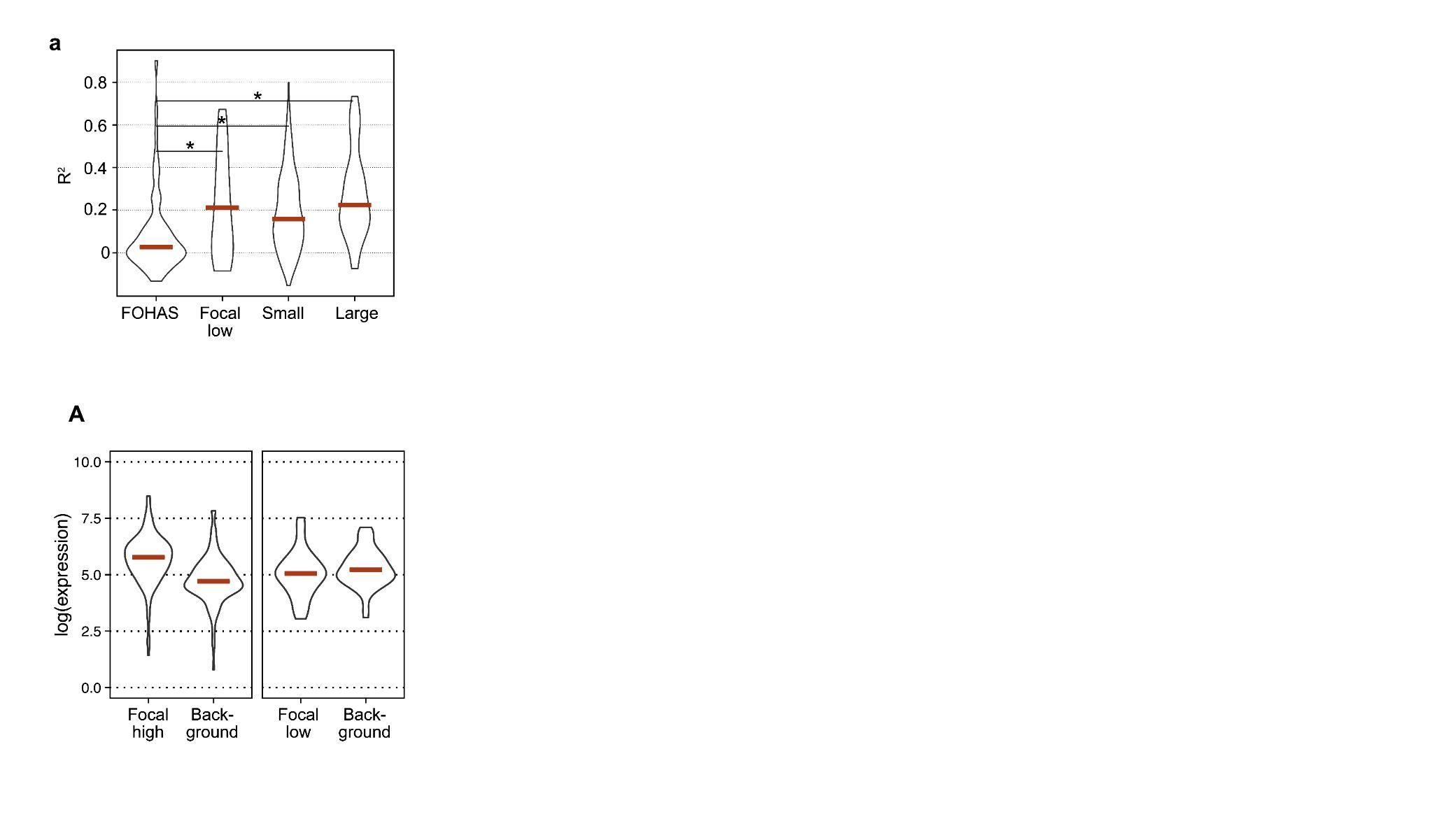

**Supplementary figure 4. Predictions of CN by expression based on CNV segment size and amplification. a**: Adjusted R squared of random forest predictions of CN by expression. Wilcoxon rank-sum test was used for p value statistics. Star indicates significance p<0.05.

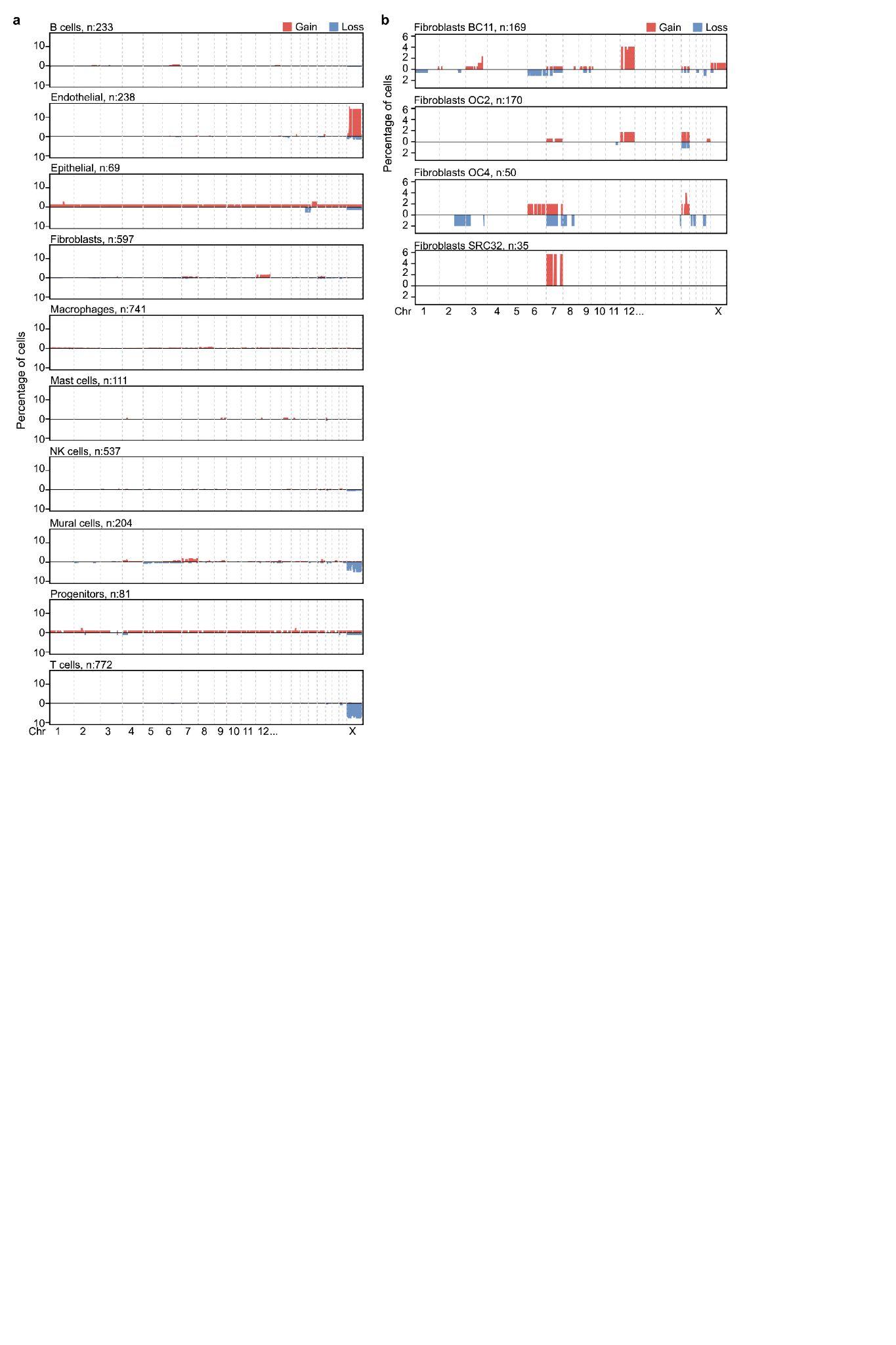

**Supplementary figure 5. Non-cancer cell CNVs. a**: All CNVs found in non-cancer cells, split by cell type, annotated by transcriptomic class. **b**: All fibroblasts with amplifications on chr 7, chr 12 or chr 18, split by patient.

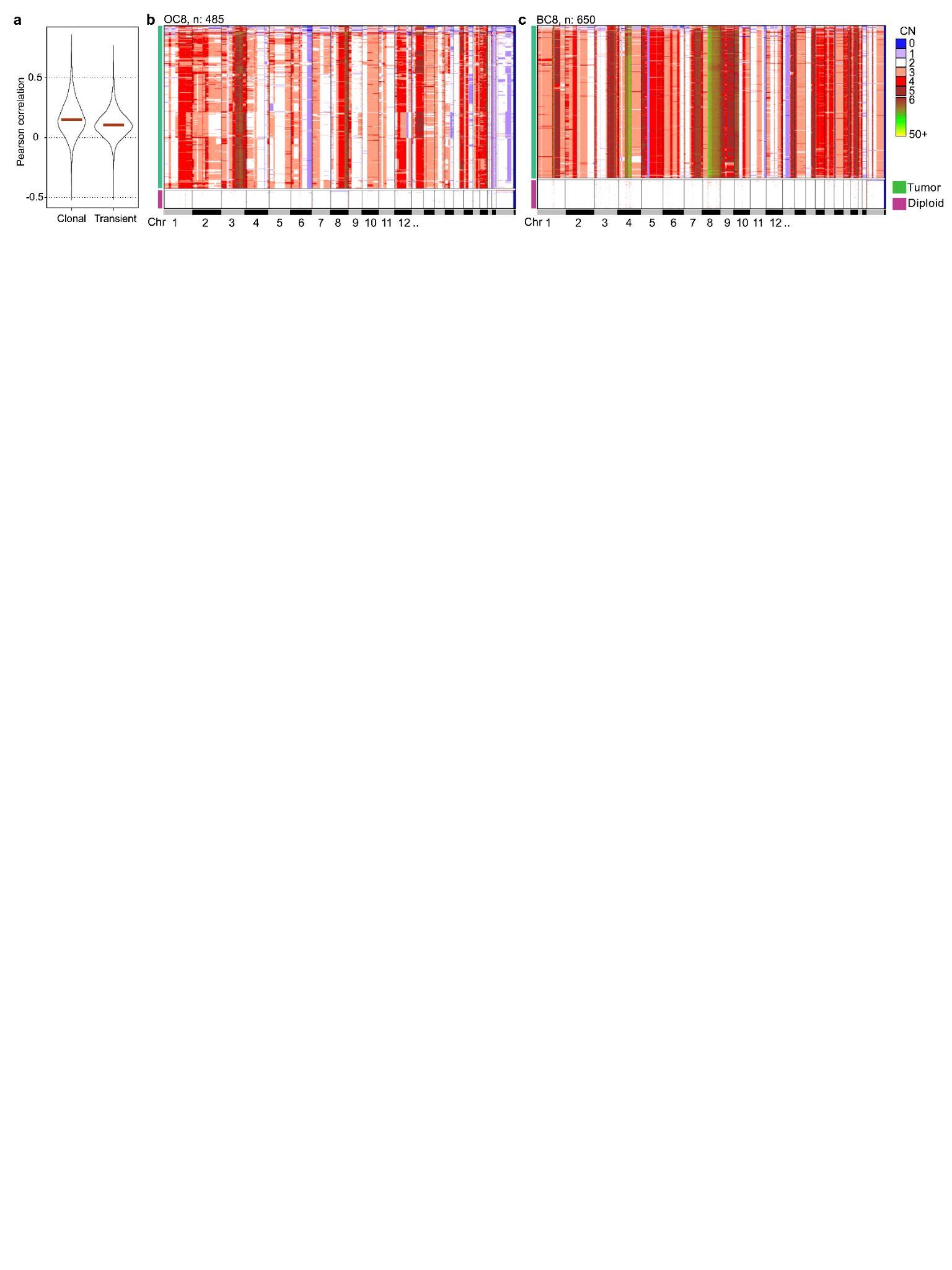

**Supplementary figure 6.Transient clonality. a**: Pearson correlation of dosage effect in clonal and transient samples. **b**: CN heatmap of OC8. Heatmap shows each single cell as a row and copy numbers are indicated by color. Aneuploid and diploid cells are separated as blocks. **c**: Same as b for BC8.

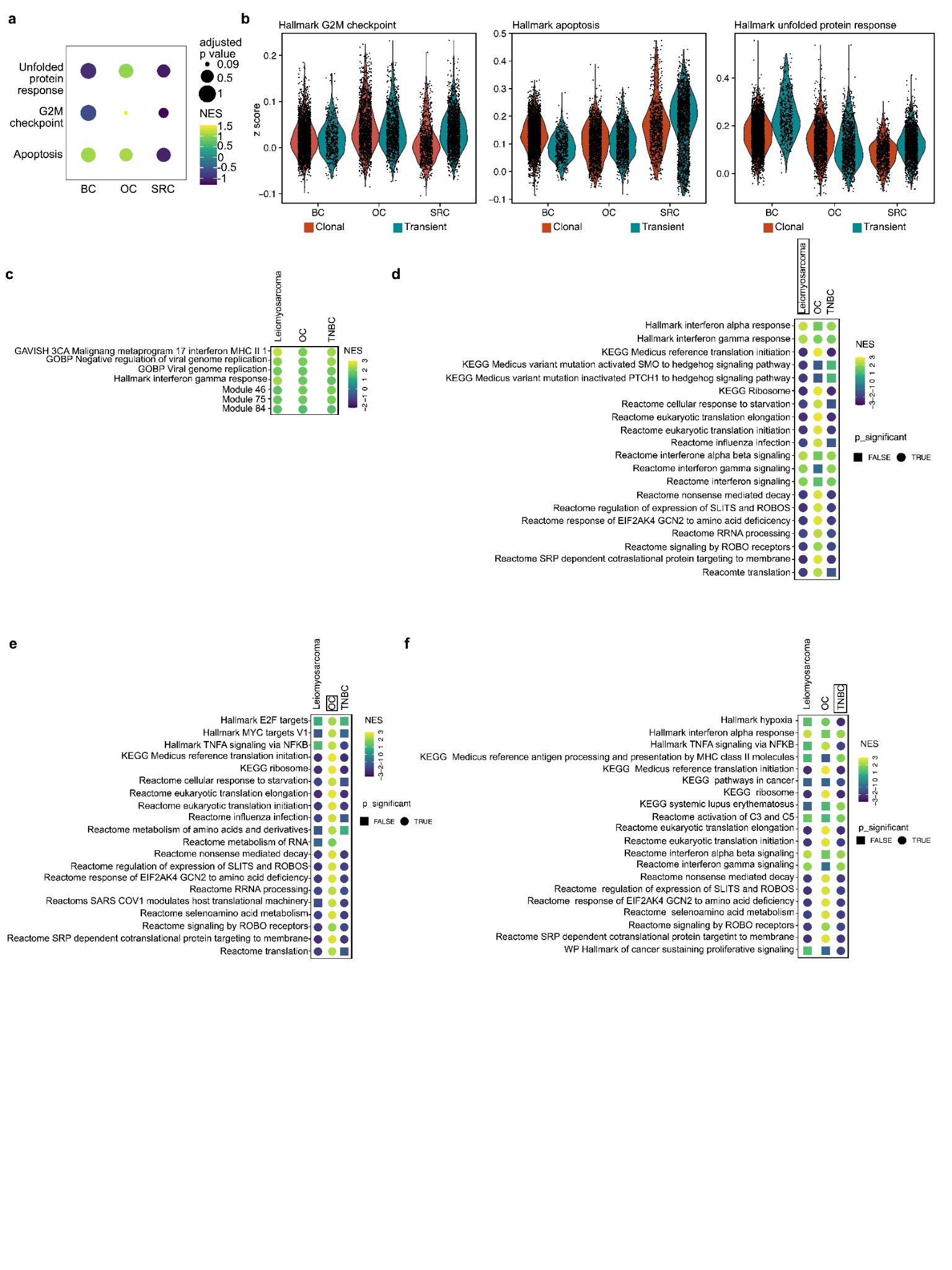

**Supplementary figure 7. Phenotypic effects of transient clonality in solid tumors. a**: Pathway analysis of BC, OC and SRC. Color shows normalized enrichment score (NES) clonal over transient cases. P value between the clonal and transient patients is shown as size. Pathways are hallmark pathways **b**: Z scores per tumor type for the three hallmark pathways from 6A split by genotype. **c**: All non-immune related pathways that have normalized enrichment scores (NES) in the same direction and are significant in all subtypes between transient and clonal samples. **d**: 20 most significantly enriched Reactome, Hallmark and KEGG pathways between transient and clonal leiomyosarcomas. Colored by NES, shape indicates significance (<0.05) **e**: Same as d for OC. **f**: Same as d and e for TNBC.
